# Dynamics of somatic L1 retrotransposition in the human brain

**DOI:** 10.64898/2026.11.16.767114

**Authors:** Ilke Demirci, Anton J. M. Larsson, Marta Paterlini, Ionut Dumitru, Michael Hagemann-Jensen, Kanar Alkass, Henrik Druid, Christoph Ziegenhain, Jakob Michaëlsson, Jeff E. Mold, Rickard Sandberg, Jonas Frisén

## Abstract

The sole source of genomic diversity in the human brain caused by autonomous retrotransposition is facilitated by the long interspersed nuclear element-1 (L1). However, assessing the rate, timing and source elements of this event has been challenging due to the lack of technologies that capture of L1s at sufficient scale in single cells. Here, we introduce LUSTRE, a single-cell assay that enables mapping L1 sites without whole-genome amplification, and profile ∼17,600 neurons from the dentate gyrus, cortex, cerebellum of the adult human brain. We report the presence of somatic L1s that define related lineages and conservatively estimate a validation-adjusted rate of 0.1-0.3 somatic events per neuron. Our analyses further indicate two distinct waves of retrotransposition during development and combined with the variability in their polyA tract lengths across single nuclei, suggest the formation of lineage hierarchies marked by new somatic retrotransposition events and cell divisions. Mapping the origin of L1s through 3’-transduction events identify the rare source L1s that are responsible for the majority of retrotransposition events. These results uncover the dynamics of L1-driven somatic mosaicism and establish L1s as endogenous recorders of cellular and lineage history in human tissues.

## INTRODUCTION

A large portion of the human genome is derived from transposable elements^1^, including DNA transposons, long-terminal repeat (LTR) retrotransposons and non-LTR retrotransposons. These are mostly inactive sources of structural variation that are silenced through mutations or epigenetic modifications^2,3^. The long interspersed nuclear element-1 (L1) family of transposable elements, a sub-class of non-LTR retrotransposons, is an exception and retains the ability to autonomously retrotranspose to new genomic loci via an RNA intermediate^4–6^. Approximately 0.02% of L1 copies have been reported to be retrotransposition-competent with functional open reading frames (ORFs)^7^, resulting in a small pool of mobile L1s with widely variable activity across individuals and cell types which continue to create diversity in the human genome^7–9^. This pool consists of L1s that show large variation in the human population and are differentially present in an individual or in individual genomes but absent from the human reference genome^7,10^.

Although it has been previously established that retrotransposition activity results in new polymorphic events in the germline and during early development in humans^11,12,13^, and in many cancers^14,15^, L1s are also a distinctive source of somatic mosaicism during later life. The potential of L1s to create somatic diversity have been largely explored *in vitro* through engineered L1 reporters in cultured cells, including neural progenitors^16–18^. Several strategies have further identified L1 retrotransposition events in the human brain through new methods that profile somatic L1 insertion sites in single neurons at variable scales^19–22^. However, the rate and prevalence of somatic retrotransposition have been under debate, where discrepancies have been attributed to whole-genome amplification (WGA) artifacts, sequencing data analysis, and validation procedures^23,24^. Recent whole-genome sequencing (WGS) approaches involving *in vitro* clonal expansion of single cells circumvent the requirement of WGA^25^. Such approaches are not applicable to mature postmitotic neurons, and sequencing costs may limit the number of cells analyzed. Consequently, somatic L1 retrotransposition in the human brain remains unresolved, where its developmental dynamics and its potential as an endogenous marker of cell lineage^20^ have not been investigated at scale.

Here, we developed LUSTRE (Lineages Using Somatic Transposable Elements), a novel molecular biology assay for the targeted discovery of L1s at a throughput of thousands of single cells. Our approach involves fluorescence activated cell sorting of single cells or nuclei into micro-well plates containing lysis buffer, followed by direct amplification of target sequences in single cells and detection of human specific L1 events by short-read sequencing. Using LUSTRE, we profiled neuronal nuclei from multiple regions of the human brain and validated a subset of the detected somatic L1s by long-range PCR and whole-genome sequencing. Based on thousands of identified germline and somatic L1 loci, we estimate a rate for L1 retrotransposition per neuron, infer a developmental period during which they arise, trace 3’ transductions to their source L1s. Finally, based on shared L1s and their polyA tract lengths, we explore L1s as a marker that reflects lineages and division histories of neurons in the human brain.

## RESULTS

### LUSTRE detects L1s in single cells with high sensitivity and throughput

We developed LUSTRE to prevent amplification errors and drop-out that arise from WGA by instead using single DNA molecules as input templates, and facilitate the targeted sequencing of thousands of single cells. We target the retrotransposition-competent L1s by exclusively priming for the subfamily L1Hs-Ta, which we hereafter refer to as L1. We leverage these consensus bases in the L1 3’ end for robust priming of template DNA, followed by tagmentation of the double-stranded DNA fragments via Tn5 transposase. Tn5 inserts a stretch of oligonucleotides into the target DNA upon tagmentation^26^. This sequence is used for exponential amplification together with a primer containing additional consensus bases downstream the initial priming site, which enables the capture of sequences containing the L1 insertion site together with the polyA tract and the genomic flanking region. This strategy allows for the amplification of unknown loci in the genome and circumvents the need for whole-genome amplification or whole-genome sequencing-based approaches (Figure 1A, STAR Methods).

**Figure 1.**
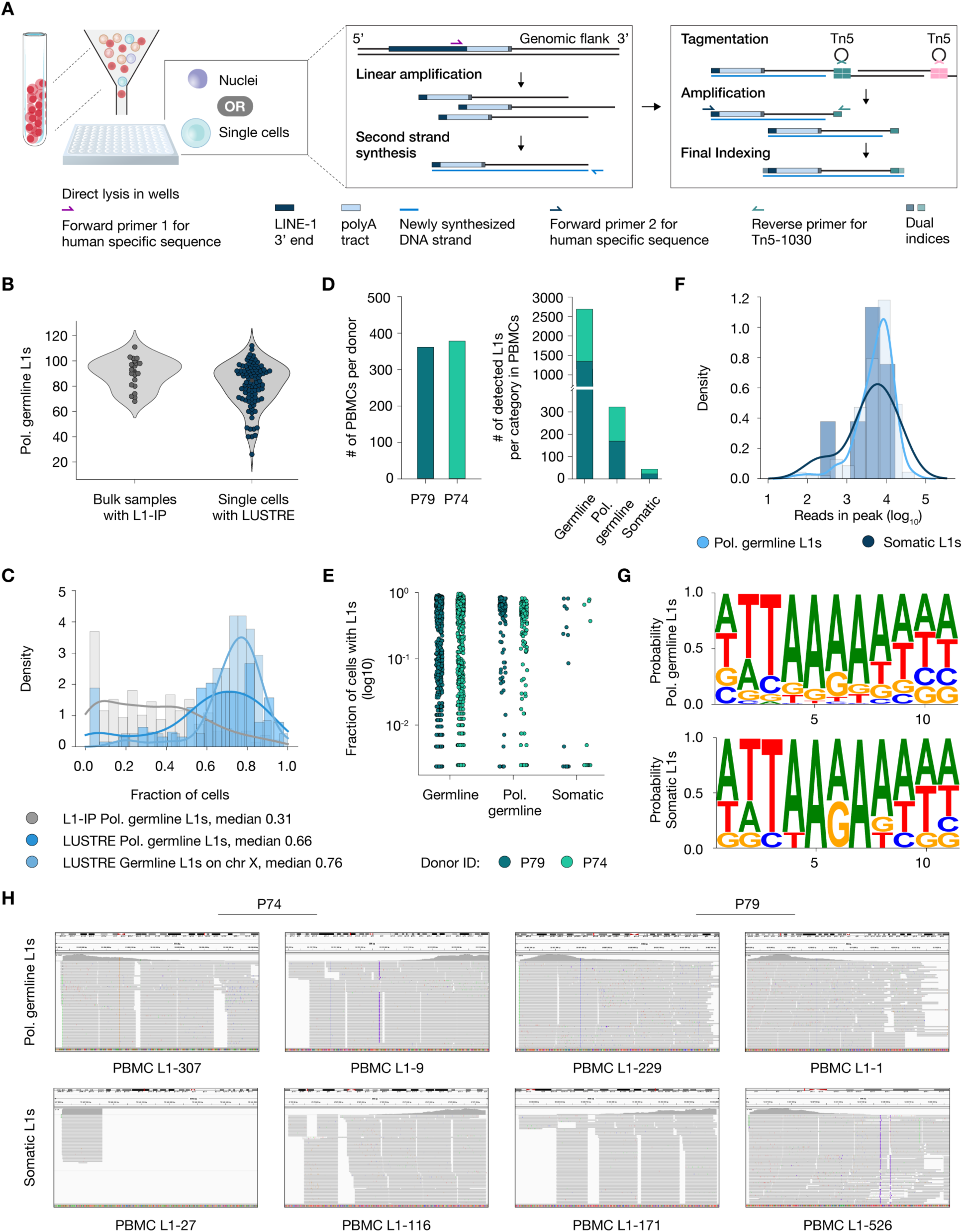
LUSTRE for targeted discovery of human specific L1 insertion sites in the single cells. (A) Overview of the method for amplification of L1 insertion sites together with the genomic flanking region. (B) Number of polymorphic germline insertion sites detected using L1-IP on bulk DNA (n = 21 samples) and LUSTRE on single cells (n = 96 cells). (C) Sensitivity to detect L1s in single cells with LUSTRE and L1-IP (n = 96 cells for both datasets). (D) Left: Number of peripheral blood mononuclear cells (PBMCs) sequenced per donor (n = 360 and 377 cells for P79 and P74 respectively). Right: Number of detected events per donor and category: germline, polymorphic germline and putative somatic events. (E) The fraction of cells where a given L1 event was detected, separated by donor and category (n = 1,344 germline, 153 polymorphic germline and 21 putative somatic L1s for P79 and 1,344 germline, 169 polymorphic germline and 23 putative somatic L1s for P74). (F) The read depth of detected insertion sites, for polymorphic germline and putative somatic L1s, *p = 0.678, independent t-test*. (G) Combined position weight matrices of sequence logos for the insertion site motifs in donors for polymorphic germline L1s (top) and putative somatic L1s (bottom). (H) Selected Integrated Genome Viewer (IGV) images for two somatic L1 and two polymorphic germline L1 for P74 (Left) and P79 (Right).

We first optimized reagents and experimental conditions that influence the performance of the method (Figure S1A). As artifactual DNA fragments can potentially arise from linear and exponential amplification^27^, we used a polymerase with a high 3’-5’ exonuclease proofreading and overall processivity (STAR Methods) compared to *Taq* polymerases previously demonstrated to cause chimeras resulting from template-switching^28^. As the rate of template switching is proportional to polymerase dissociation frequency, artifactual DNA can be reduced by higher processivity proofreading polymerases during amplification^29^. Moreover, we developed a custom analysis pipeline for LUSTRE which enabled stringent data analyses to account for such artifacts (Figure S2), where multiple independent criteria are required to call each putative L1 and recapitulate the overall molecular signature of a bona fide retrotransposition event (STAR Methods). We confirmed this in peripheral blood mononuclear cells (PBMCs) from a healthy male human donor (P79) and detected similar numbers of polymorphic germline elements in single cells using LUSTRE compared to a bulk PBMC sample prepared using a previously established short-read sequencing approach (Figure 1B, L1-IP with non-WGA bulk sample, reproduced from^11,19^). We next compared LUSTRE to L1-IP applied to single cells and found an increase in sensitivity from 0.31 (L1-IP, MDA-amplified single cells) to 0.66, based on polymorphic germline L1s and 0.76 based on known germline L1s present on the X chromosome (Figure 1C). The X-linked estimate is based on a small set of events known to be present in each nucleus, whereas the polymorphic germline estimate includes lower frequency events with a cut-off of >1 nuclei. Therefore, the higher sensitivity for X-linked germline L1s (0.76) compared to autosomal polymorphic germline L1s (0.66) are not attributable to the copy number differences.

To enable higher cellular throughput, we adapted LUSTRE to the 384-well format and applied it to 384 PBMCs from two male donors (P79 and P74), where most cells passed quality filtering (94% and 98% of cells for P79 and P74 respectively). Most L1 insertions present in the reference genome were detected (90% for P79 and 88% for P74), in addition to the expected number of germline polymorphic L1s and ∼20 putative somatic events per donor (Figure 1D, Figure S1B-C). The reference germline L1s had higher detection variability compared to polymorphic germline L1s (Figure 1E), which we confirmed was due to the specific amplification of the retrotransposition-competent subfamily while amplifying other families much less efficiently (Figure S1D). Moreover, the putative somatic events in the PBMCs had read depths comparable to germline L1s (*p* = 0.678, independent t-test, Figure 1F). Additionally, the detected insertion motifs were consistent with target-primed reverse transcription for both germline and putative somatic insertion sites (Figure 1G). As expected, the sequencing reads covering the genomic flank were consistently oriented in the opposite orientation to the insertion for both polymorphic germline and putative somatic L1s (Figure 1H).

To independently validate the detected L1s, we amplified seven polymorphic germline and five putative somatic insertions using primers targeting the predicted flanking regions and analyzed using long-read sequencing (Figure S3A, B). Two out of five putative somatic L1s could be phased using a nearby SNP, where the SNP was also detected without the insertion present in a subset of the reads – suggesting these events are mosaic in the individual (Figure S3C). Lastly, the L1 insertion sizes, polyA tract lengths and target-site duplication (TSD) lengths of these somatic insertions were consistent with our expectation based on existing literature^20^ (Figure S3D). Therefore, we concluded that LUSTRE can be used for high sensitivity detection of somatic L1 retrotransposition events, which were validated by long-read sequencing.

### High-resolution mapping of somatic L1 retrotransposition in the human brain

We next applied LUSTRE to neurons of the adult human brain and chose three regions with different developmental histories: the dentate gyrus of the hippocampus and the cortex, which both derive from the dorsal telencephalon, and the cerebellum which derives from the hindbrain^30–33^. To investigate the prevalence of somatic L1 retrotransposition and establish more precise estimates in neuronal lineages, we biopsied these brain regions from 3 postmortem donors and subjected 18,816 NeuN^+^ nuclei to LUSTRE (Figure 2A), where the libraries were sequenced at a budget of at least 250,000 reads per nucleus (Figure S4A-I). After demultiplexing and quality-control (>10,000 reads, STAR Methods), 7787, 7557 and 2251 NeuN^+^ nuclei were included for downstream analyses where 172, 432 and 87 somatic L1s were detected in the dentate gyrus, cortex and cerebellum respectively (total 17,595 NeuN^+^ nuclei, Figure 2B).

**Figure 2.**
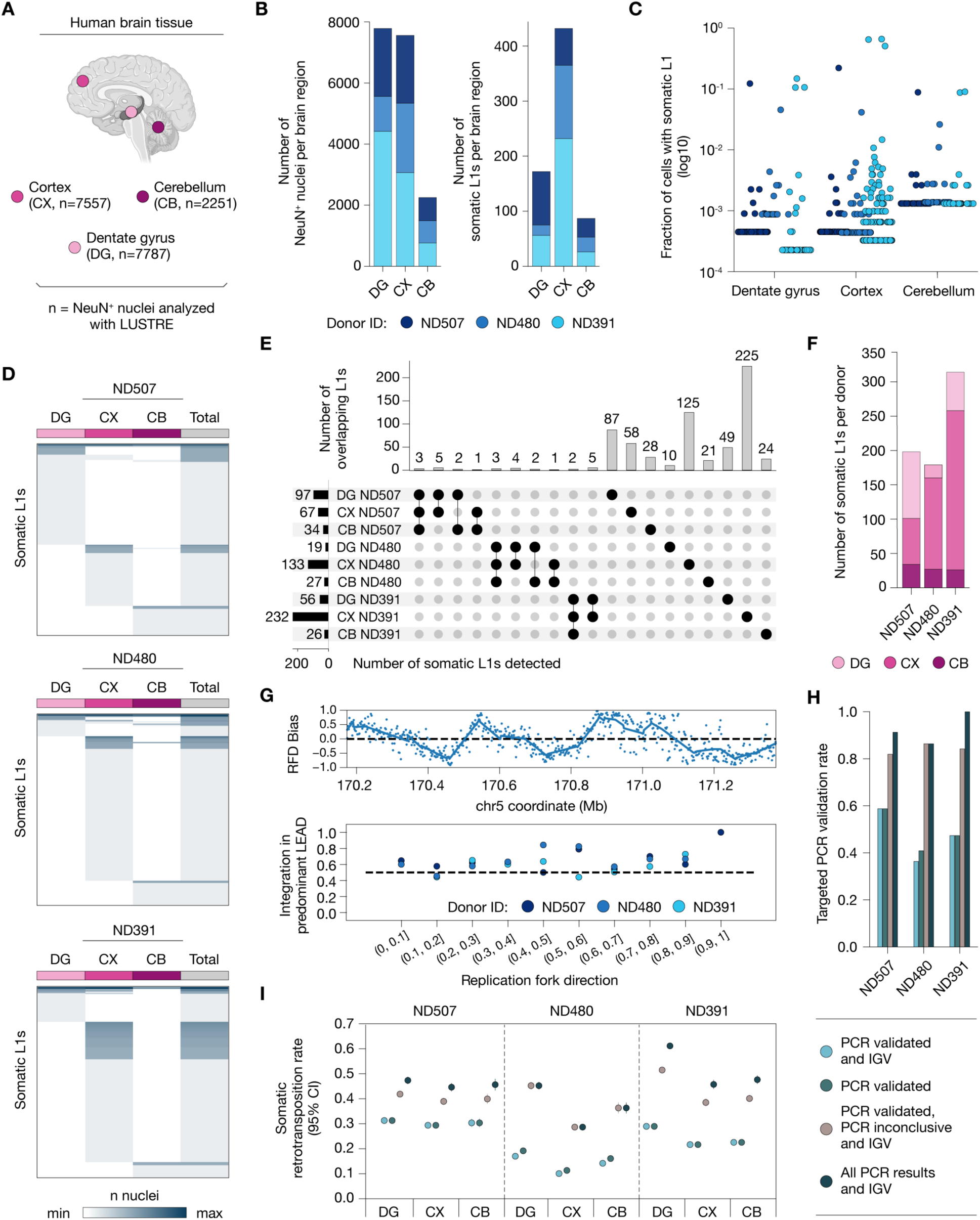
Large scale profiling of L1 retrotransposition events in the adult brain. (A) Experimental setup. (B) Number of total nuclei sequenced per region across all donors (left) and number of total putative somatic L1s detected per region across all donors (right). (C) Fraction of nuclei in which a given somatic L1 was detected, separated by region and donor. (D) Heatmap of the detection of somatic L1s for each region in the three donors. (E) Up-set plot showing the overlap of detected insertions across brain regions within each donor. (F) Number of total somatic L1s detected per donor across all regions (G) Top: Reproduction of replication fork direction (RFD) bias estimates based on the analysis described by Flasch et al. compared to Figure 5A in Flasch et al. Bottom: Fraction of L1s predominantly integrated in the leading strand (y-axis) as a function of replication fork direction bias (x-axis, absolute value). Data point from each donor shown separately. (H) Validation rates for selected L1s by PCR validation, based on different criteria for validation. PCR validated: PCR supports the existence of a larger DNA fragment. PCR inconclusive: No evidence by PCR that a larger DNA fragment was amplified. IGV: No indication of inconsistencies after manual inspection of read coverage using the Integrated Genome Viewer (IGV). See Note S1 for more details. (I) Sensitivity and validation adjusted mean rate estimate for each region and donor.

We first measured the sensitivity of LUSTRE in single neurons based on the number of polymorphic germline L1s detected and the number of X-linked germline L1s in male donors ND391 and ND507. We found that the performance was mainly uniform across donors and brain regions, and comparable to the previous application in PBMCs, except for the dentate gyrus in donor ND391, which may be due to tissue quality and processing (Figure S5A-D, Figure S6A). Next, to ensure the somatic L1s were not erroneously detected due to index switching^34^, we determined a read cutoff threshold which excludes the L1 calls that are based on reads likely to originate from another nuclei (STAR Methods, Figure S6B). The vast majority of reads close to known L1 were found in the correct orientation within 1 kb of the insertion site (98.9% below 1kb within 10 kb windows, Figure S6C and S6D). In agreement with the PBMC data, the somatic L1s were detected at read depths comparable to polymorphic germline L1s, for both somatic L1 found in multiple nuclei and in a single nucleus (Figure S6E). Furthermore, while a small number of somatic L1s were more widespread among different nuclei isolated from a single donor (L1-2285 detected in ND507, 16% of nuclei, L1-6480 detected in ND480, 4% of nuclei and L1-248, L1-133, L1-210 detected in ND391, both 30% of nuclei), the majority of the remaining somatic L1s were harbored by a small fraction of nuclei in each respective donor (found in only one nuclei: 156/184, 84.8% in ND507; 143/166, 86.1% in ND480; 229/305, 75.1% in ND391; 528/655, 80.6% across all donors, Figure 2C). A majority of the somatic L1s present in multiple nuclei were detected across different sample plates from the same donor and only four somatic L1s found in more than three nuclei were detected in only one plate, providing orthogonal evidence of biological origin as the probability of the same technical artifact arising at an identical genomic locus on independent sample plates is negligible (65 out of 127 somatic L1s, Figure S7A-D). Lastly, to understand whether there are any region-specific differences in retrotransposition, we considered the relationship between the number of detected somatic L1s and sequenced nuclei and found no differences between regions after controlling for this factor (ND391 DG was detected as an outlier using studentized residuals, STAR Methods, Figure S8A-D).

We next sought to identify neurons related through cell division, as retrotransposition occurs in a cell division dependent manner and resulting L1s propagate in the progeny through this process^35^. We first investigated the burden of somatic L1 events across brain regions and found that only 6% of somatic L1s for ND507 and ND480, and 2% of somatic L1s in ND391 were shared in multiple regions of the brain (Figure 2D) Conversely, most somatic L1s were detected within one brain region, with cortex and dentate gyrus more often sharing insertions compared to the cerebellum (Figure 2E). Overall, 184, 166 and 305 somatic L1 events were detected in the donors ND507, ND480 and ND391 across all brain regions (Figure 2F). Using this newly identified set of somatic L1s, we grouped neurons based on the L1s they harbor within this set and observed that these L1s defined separate groups of presumably related cells, where the presence of shared somatic L1s indicate a common telencephalic origin of the dentate gyrus and the cortex (Figure S9A). Taken together, these results indicate ongoing somatic L1 activity during neurogenesis which reflect developmental histories, in agreement with previous reports^20,25^.

We further explored the ability of somatic L1s identified through LUSTRE in reflecting biological processes. L1 retrotransposition have previously been shown to be correlated to replication fork directionality during cell division using *in vitro* experiments^36,37^. LUSTRE enables this analysis to be performed in our *in vivo* neuron datasets based on the somatic L1 events. We processed the same OK-seq^38^ dataset used by Flasch et al. and obtained the same underlying replication fork analysis they reported (Figure 2G, top, Figure S9B-C). We reproduced the same relationship between replication fork bias and somatic L1 insertions, where the orientation of somatic L1s was biased with respect to the fork direction, indicating that these events arose in replicating cells (Figure 2G, bottom). Moreover, somatic L1s were more likely to be present in the introns of genes, at higher abundance than expected by chance (Figure S9D). However, because of the requirement for high mapping quality in the LUSTRE analysis pipeline, this enrichment may partly reflect a detection bias favoring unique-sequence contexts rather than a strict biological preference for genic retrotransposition. The Shannon diversity was similar across regions and donors, with higher variability in dentate gyrus samples (Figure S9E). Despite the technical effects originating from human brain samples, this variability may reflect developmental heterogeneity among sampled neurons in the dentate gyrus, consistent with distinct progenitor dynamics and postnatal neurogenesis^39,40^.

To further confirm the validity of the somatic L1s identified in this study, we selected many L1s detected in multiple nuclei and other L1s found in only one nucleus, totaling 12 polymorphic germline and 63 somatic L1 events (n = 22, 22 and 19 loci for ND507, ND480 and ND391 respectively, Note S1). Since the lack of whole genome amplification step in LUSTRE precludes additional experiments on the sequenced nuclei, primer pairs were designed for each selected L1 and amplified using high-molecular weight DNA extracted from the same donor and sample region by long-range PCR in 125 samples across all brain regions (STAR Methods, Note S1). The primer pairs were designed to span approximately 1500-2000 bp upstream and downstream of the 5’ and 3’ ends of the identified L1s. Therefore, resulting fragments larger than the expected size were indicative of somatic L1 sequences. After agarose gel visualizations, we categorized each insertion according to multiple criteria, based on the PCR results and manual inspection of LUSTRE sequencing reads using the Integrated Genome Viewer (IGV). Based on this categorization, we calculated four different validation rates based on increasingly stringent requirements (STAR Methods). Using the most stringent requirements, we calculated a validation rate of 59%, 36% and 47% for ND507, ND480 and ND391 respectively (Figure 2H, Note S1). The overall validation rate was identical for L1s detected in a single nucleus and L1s detected in multiple nuclei (39% for both). Because L1s detected in multiple nuclei are additionally supported by their detection across independent sample plates and are therefore likely to be true events, this shared rate may reflect the false-negative rate of the long-range PCR rather than the specificity of LUSTRE calls. Notably, L1-2285 and L1-5439 in ND507 and L1-4062 in ND480 were identified as full length somatic L1s, providing further evidence that full-length, and therefore potentially retrotransposition-competent, somatic L1s arise in the human brain. The majority of the remaining somatic L1s were identified as non-full length, whereas some did not yield a conclusive PCR product. Some of the L1s indicated technical limitations to the validation procedure. For example, the most abundant L1 found in ND480 (L1-6480) did not yield a conclusive PCR product despite its detection in approximately 150 nuclei across all three brain regions. At a calculated allele frequency of approximately 2%, this is unlikely to be explained by a spatial mismatch between the tissue profiled with LUSTRE and the one used for PCR validations alone. Several locus-specific factors may instead contribute to PCR failure for this L1 event, including repeat-rich flanking sequences that impede priming or formation of secondary structures in the amplicon. Several remaining L1s highlighted a possible limitation of validating rare events using a bulk DNA-based approach. Notably, L1-10286 in ND480 and L1-258 in ND391 were initially detected as higher prevalence events in single brain regions and supported by high-quality sequencing reads. However, neither event provided conclusive support by long-range PCR, possibly due to differences between the initially selected tissue section of the donor sample and the one used for PCR validation. Because targeted PCR can fail for reasons unrelated to the presence of an insertion, we sought independent support for L1s by WGS. We sequenced bulk DNA from each of the three regions of each donor, and from a fourth donor who was not profiled by LUSTRE (NT022) as a negative control (Note S1). Given the presence of L1s across a given number of nuclei as measured by LUSTRE and the local sequencing depth, only 11 of the 655 somatic L1s were initially expected to yield sufficient supporting reads. We detected 8 of these L1s by WGS (Note S1; Figure 4-6). 4 additional somatic L1s initially identified by LUSTRE in a single nucleus each were also supported by WGS, which suggests a larger clone size in the tissue than expected based on their initial prevalence. None of the somatic L1s detected by LUSTRE had supporting reads in NT022. Moreover, out of the 11 L1s expected to be detected by WGS, 9 were initially tested by targeted long-range PCR. 8 L1s were confirmed by WGS, including 5 that PCR had not fully validated. 3 of these were the most abundant L1s: L1-6480 in ND480, and L1-248 and L1-210 in ND391; therefore, a lack of PCR support may not exclude the presence of an L1. Conversely, most L1s validated by PCR could not be validated by WGS, where the majority of these L1s were below the detection limit of WGS and could not be assessed (Note S1, Figure 6). Overall, 35 of the 63 tested L1s (56%) had conclusive support by PCR, WGS or both. Notably, bulk WGS could assess fewer than 2% of the L1s identified by LUSTRE, which indicates that most somatic L1 events can be detected only by single-cell analyses.

**Figure 3.**
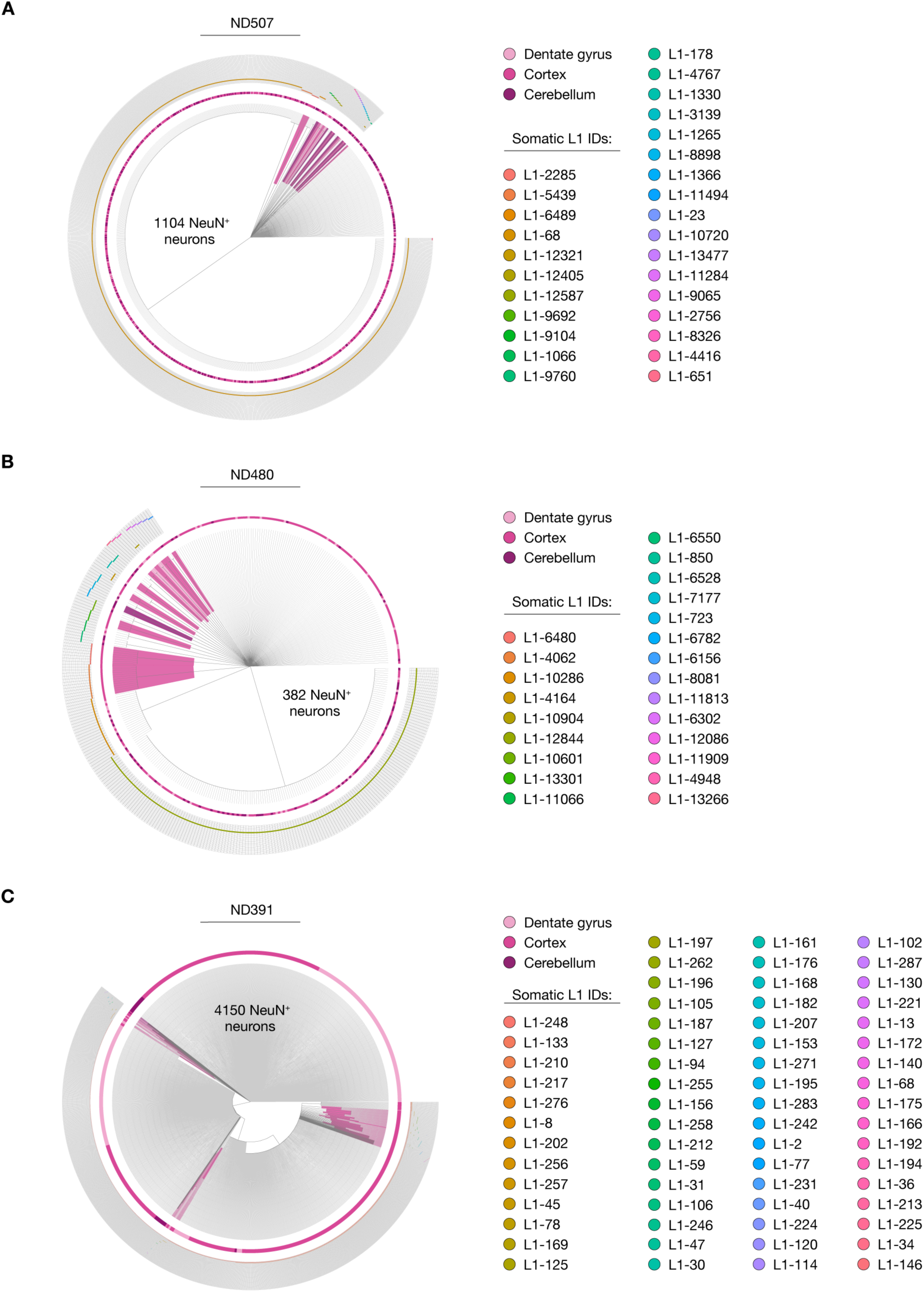
Phylogenetic trees based on somatic L1Hs events. (A-C) Visualizations of trees obtained using ScisTree and tskit, based on somatic L1 events, with branch lengths defined by somatic L1 divergence. Each NeuN^+^ nuclei have been colored by brain region of origin in ND507 (A), ND480 (B), and ND391 (C). Branches marked by a color indicate clades originating from a common ancestor in a single brain region.

**Figure 4.**
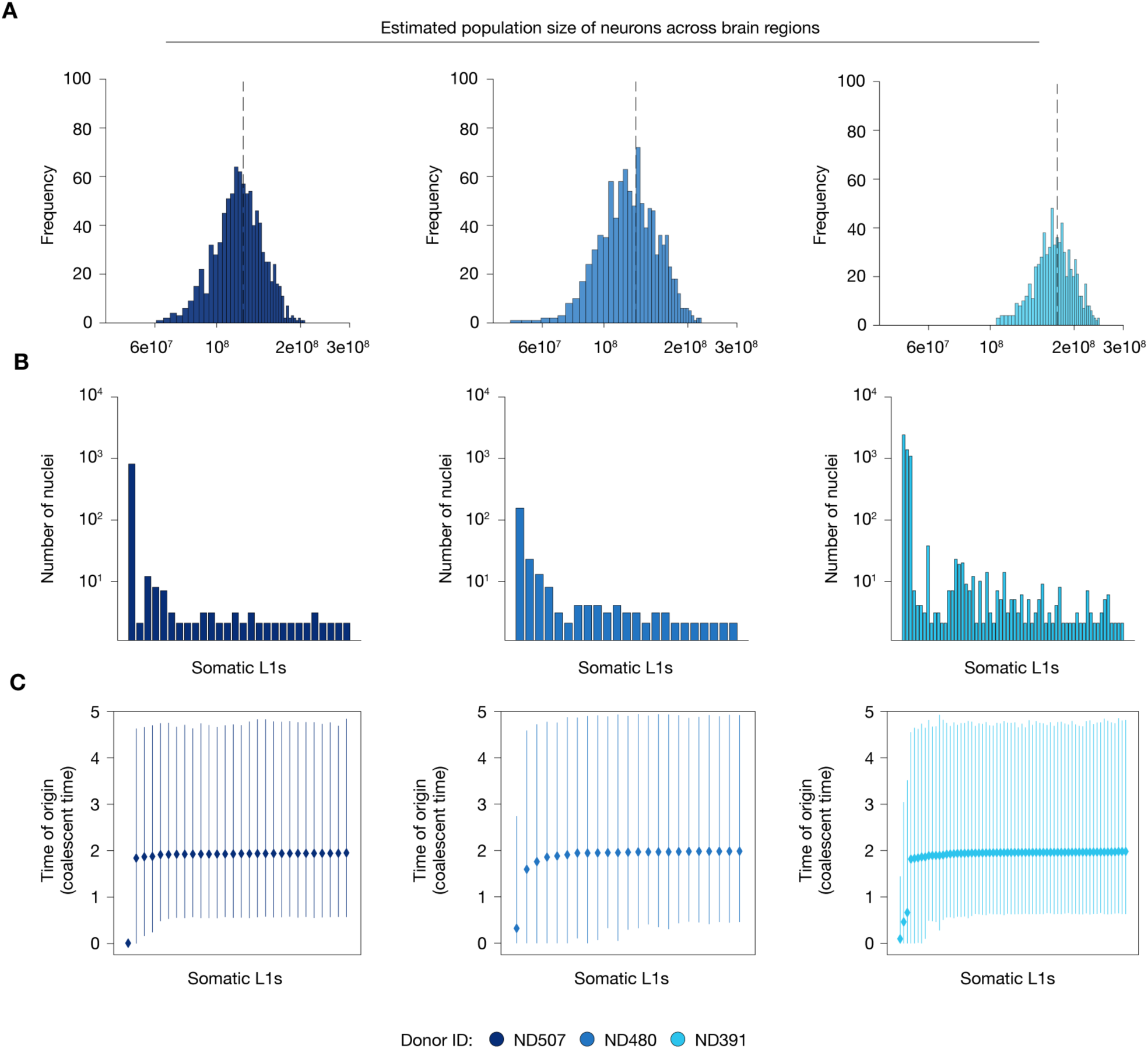
Population size and timing estimation of somatic L1 events. (A) Population size estimates after the rejection-sampling procedure based on the Kingman’s coalescent as the underlying model using the somatic L1s. Please see STAR Methods and Note S2 for more information. (B) Bar plot showing the number of nuclei where a somatic L1 was detected that is present in the trees shown in Figure 3. Each bar corresponds to a somatic L1 detected in >1 neuron (n = 28, 23, 64 somatic L1s for ND507, ND480 and ND391 respectively). (C) The estimated time of origin for somatic L1s since the MRCA based on the coalescent model. The estimates for each somatic L1 are shown in the same order as presented in (B).

**Figure 5.**
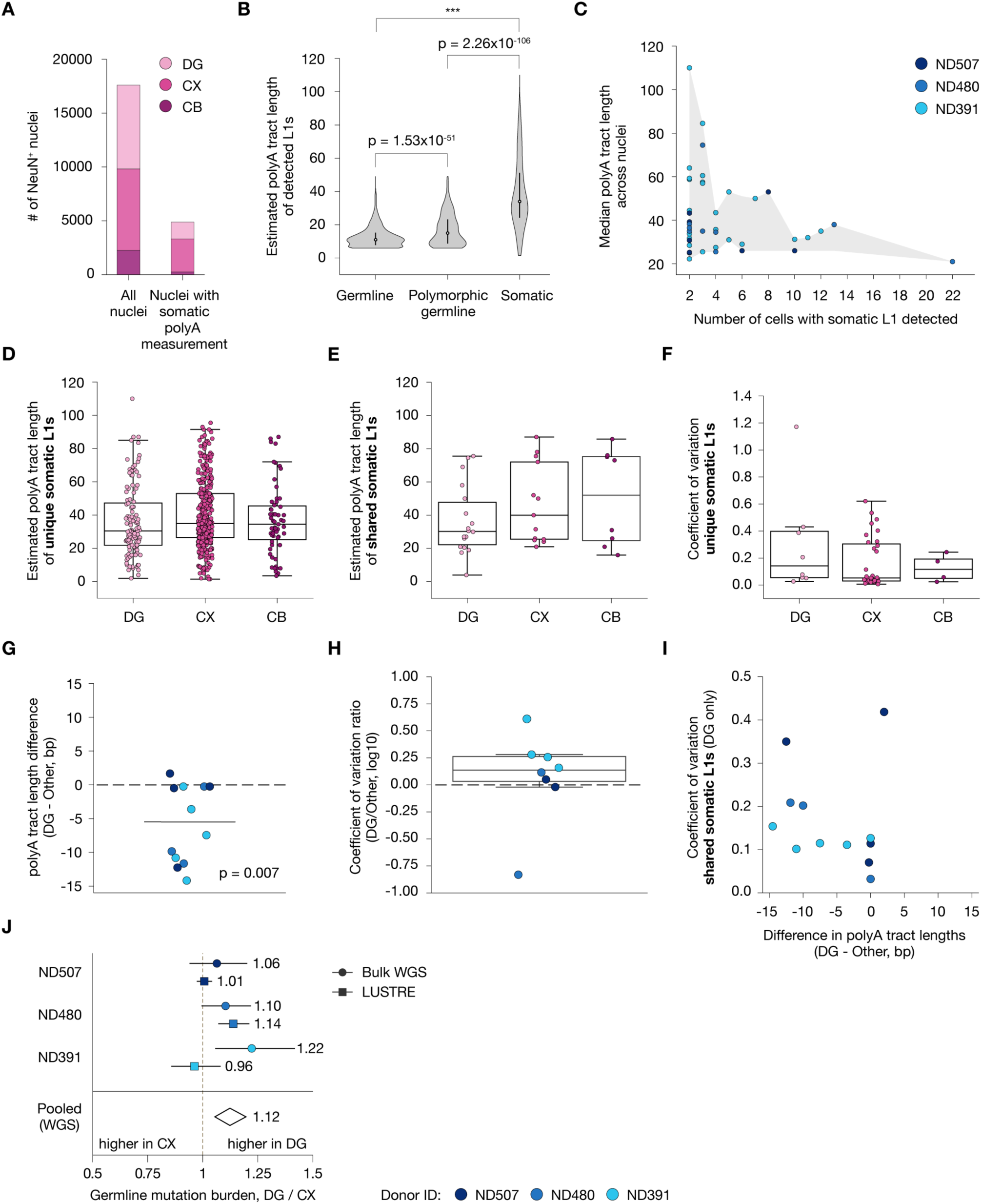
Variation in polyA tract lengths harbored by L1. (A) Total number of nuclei sequenced in this experiment, separated by region, and the number of nuclei with at least one somatic L1 with polyA tract quantified. (B) The median (across cells) estimated polyA tract lengths for germline, polymorphic germline and somatic insertions (all donors, n = 5,923, 698 and 525 germline, polymorphic germline and somatic polyA tracts respectively, independent t-test). (C) The relationship between the polyA tract length and number of detected cells (n = 44 tracts). (D-E) Estimated median polyA tract lengths of insertions found in only one region (unique) (D) and detected across multiple regions (shared) (E). (F) Coefficient of variation of polyA tract estimates of somatic insertions unique to a region. (G) Difference of median polyA tract lengths between the dentate gyrus (DG) and cortex (CX)/cerebellum (CB) for insertions shared between DG and at least one of CX/CB. Paired t-test (n = 12 tracts). (H) Ratio of coefficient of variation between DG and CX/CB insertions. (I) Relationship between polyA tract difference and the coefficient of variation of L1s detected across regions (shared). (J) Relative difference in germline polyA mutation burden between the DG and CX regions for LUSTRE (square data points) and bulk WGS (round data points).

**Figure 6.**
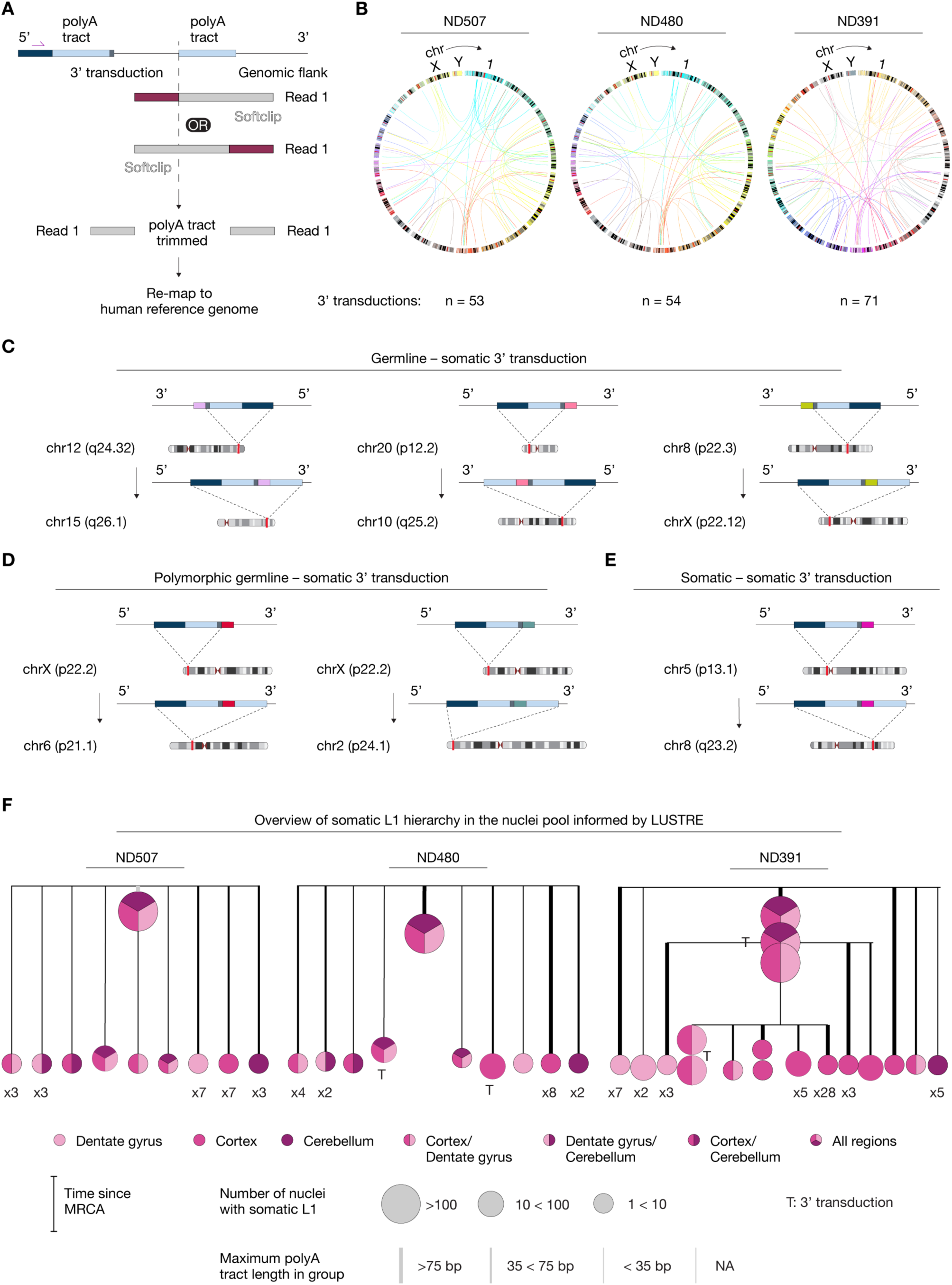
L1 transduction detection and phylogeny using LUSTRE. (A) Detection of 3’-transduction events through computational analysis of sequencing reads. (B) Detected 3’-transduction events for the germline and somatic insertions for each donor shown with a Circos plot. The color of the line denotes the source insertion chromosome. (C-E) Detected somatic retrotransposition events with observed 3’-transduction, where the source insertion is a germline (C), polymorphic germline (D) or somatic (E) insertion. (F) Mutation trees for each donor, where each insertion has been scaled by cell number, colored by detected region(s) and placed relative to the root based on the mutation dating estimate. The number of insertions which share the same characteristics are shown (e.g. x2, x5). MRCA: Most recent common ancestor.

We then used a sensitivity and validation rate adjusted procedure to estimate the number of somatic L1s harbored by each neuron that occurred during neurogenesis (STAR Methods) and therefore excluded the three most abundant insertions found in ND391 as they may have occurred in developmental stages preceding gastrulation based on their detection in the fraction of nuclei in the sampled population. We found an average of 0.1-0.3 somatic L1 events per neuron across three brain regions using the most stringent validation estimates, with a small but consistent increase in the dentate gyrus compared to the cortex and cerebellum (Figure 2I). However, because 5 of the L1s that PCR failed to validate were confirmed by WGS, this estimate is conservative. In conclusion, this result is within the range of previous retrotransposition estimates of 0.07 and 13.7 somatic L1 events per single-neuron^19,22^.

### Inferring the timing of somatic L1s in single neurons

The somatic retrotransposition events found by LUSTRE may further elucidate the timing of L1 retrotransposition during development in humans. By analyzing the number of somatic L1s detected in a given set of nuclei, we observed a log-log relationship up to around 10 NeuN^+^ nuclei, suggesting a power law relationship commonly found in statistical genetics^41^ (Figure S10A).

We thus reasoned that the somatic L1s inform the relationships among sequenced nuclei and can be used to estimate a genealogical tree where neurons are ordered across molecular time based on the specific somatic L1s they harbor which are shared between multiple nuclei (Figure 3A, 3B, 3C). Interestingly, we consistently observed that the cortex contained clades defined by somatic L1s shared among neurons at earlier molecular times more often compared to the other two regions across all three donors (highlighted clades, Figure 3A-C). We analyzed these trees using population genetics models and estimated the population size and time from the most recent common ancestor (MRCA) for each somatic L1 separately (STAR Methods and Note S2). All donors had similar estimated population sizes, with a posterior estimate of 174, 130 and 125 million cells for ND391, ND480 and ND507 respectively (Figure 4A). Moreover, we investigated the robustness of the obtained estimates and found that the specified mutation rate determines the estimated effective population size (Figure S10B) but importantly does not affect the estimated population mutation rate (Figure S10C). These estimates were not sensitive to the chosen acceptance window during rejection sampling (Figure S10D and S10E). Moreover, we found that the widely observed somatic events occurred close in generations to the MRCA, while most somatic insertion events were suggested to occur much later in development (Figure 4B, 4C). This pattern was also consistent with model predictions when only considering the fraction of nuclei the somatic L1s were detected in (Figure S10F). We next evaluated whether somatic L1s harbored by only a small fraction of nuclei approach the limit of detection due to assay sensitivity and read depth variation across the population, which can deplete the intermediate events and result in an observed gap between the retrotransposition windows. To ensure that this observation cannot arise due to incomplete sampling of somatic L1s across cells, we simulated a neutral coalescent site frequency spectrum subjected to binomial sampling at different sensitivity levels and found that incomplete detection of events did not reproduce the observed gap (Figure S10G). The coalescent model further assumes a mixed population of constant size (STAR Methods), whereas neurogenesis occurs through spatially structured clonal expansions and asymmetric divisions^42^. As this can bias phylogenies, we simulated genealogies under a constant retrotransposition rate in various possible scenarios and found that none fully reproduced the absence of intermediate frequency somatic retrotransposition events (Figure S11A-B). Taken together, the lack of detected somatic events between the early and late group suggests biological significance and the existence of two distinct time windows of active retrotransposition during neurogenesis, in addition to previously reported activity in embryonic stem cells^12^, and in colorectal epithelium^25^.

### Lineage histories are recorded in the polyA tracts of L1s

A bona fine L1 carries a 3’ polyA tract copied from its polyadenylated RNA, where a newly retrotransposed L1 typically harbors a longer tract at first and rapidly shorten over generations of cell divisions^37,43,44^. These tracts are prone to replication slippage like other homopolymers, and length changes in microsatellites have previously been described a molecular clock in tumorigenesis and demonstrated to reflect cell divisions^45–47^. Notably, variation in tract length at a single somatic L1 insertion distinguished sublineages within a clone of neurons harboring that insertion in the human brain^20^. We therefore reasoned that the polyA tracts sequenced in LUSTRE together with every L1 insertion site, may carry information on the division history of neurons. To understand the variation in polyA tract lengths across and within each L1, we developed two separate polyA tract estimation procedures due to the technical challenge of estimating very long homopolymers harbored by somatic insertions (STAR Methods and Figure S12). Using these two approaches in combination, we were able to estimate polyA tract lengths for germline and somatic L1s (Figure 5A). Consistent with previous findings, polymorphic germline L1s had longer tracts compared to germline L1s fixed in the human genome^49^ (median 15 and 11, respectively). The tracts harbored by somatic L1s were longer than both categories of germline L1s (median 34 bp), where some exceeded 100 bp as expected from the relative ages of the three categories of L1s (Figure 5B).

We first asked whether the tract length of a somatic event reflects its age, as the tracts of insertions arising later in development would have shortened over fewer cell divisions. The most abundant somatic L1s we detected (L1-2285 in ND507, L1-6480 in ND480 and L1-248, L1-133, L1-210 in ND391) had a median polyA tract length of 77 base pairs (76 in ND507; 84 in ND480; 72, 51 and 77 in ND391), whereas the median polyA tract length of polymorphic germline L1s was 15 base pairs. Interestingly, somatic L1s found in fewer cells tended to have longer tracts (Figure 5C). Therefore, these somatic L1s with longer tracts may indicate retrotransposition events occurring later in neuronal development. This finding prompted us to compare the polyA tract lengths across the three different regions we sampled, hypothesizing the lengths may inform cell division dynamics. While polyA tracts harbored by insertions unique to a region did not show any differences in lengths (Figure 5D), the nuclei in the dentate gyrus had shorter polyA tracts in L1 insertions shared across regions (Figure 5E). The variation in polyA tract lengths across nuclei were not different for L1s unique to a region, and we postulated that the timing of the events may confound such analyses (Figure 5F). Importantly, the insertions detected across regions enabled a paired analysis to exclude any confounders. Indeed, the polyA tracts in the dentate gyrus were shorter by an average of 5.5 bp compared to the same polyA tract measured in the other two regions and was consistent in all three donors (*p* = 0.007, paired t-test, *n* = 12 somatic L1s, Figure 5G). Notably, tract lengths were also more variable between nuclei in the dentate gyrus for 6 of the 8 somatic L1s for which variability could be compared (Figure 5H), and the most variable polyA tracts in the dentate gyrus tended to have shorter tracts compared to the other two regions (Figure 5I). We additionally analyzed the germline polyA tracts in the LUSTRE and bulk WGS data, and quantified the mutation burden of each region as the deviation of tract lengths from the constitutional length (STAR Methods). This mutation burden was higher in the dentate gyrus than in the cortex, with ratios of 1.06, 1.10 and 1.22 for ND507, ND480 and ND391, respectively, and 1.12 when pooled (95% CI), whereas the corresponding ratios from the LUSTRE data were 1.01, 1.14 and 0.96 (Figure 5J). Taken together, the shorter and more variable tracts harbored by the same L1s in the dentate gyrus suggests that more frequent slippage events separate dentate granule neurons from the ancestral L1 rather than the cortical or cerebellar neurons. This may reflect a higher number of cell divisions in the lineages giving rise to dentate granule neurons compared to the cortex and cerebellum as measured from the somatic events to the established non-dividing population of neurons we sampled in adult age.

### Tracing L1 activity in the soma based on 3’ transductions

Which L1s cause somatic retrotransposition is an outstanding question, with most studies reporting retrotransposition activity in the germline or *in vitro*. Importantly, L1 is known to commonly transduce parts of the flanking genomic DNA to the new genomic location during retrotransposition, which occurs when the 3’ processing complex bypass the polyadenylation signal harbored by the L1 and instead relies on another signal present in the flanking DNA. These 3’ transductions provide a clue as to which L1s are active at different stages of development *in vivo.* To demonstrate the capability of LUSTRE to detect 3’-transductions we extracted reads which span the transduced DNA and the new flanking region and conservatively assigned source elements by considering uniquely mapping reads in both directions (Figure 6A). This conservative approach discovered 53, 54 and 71 transduction events in donors ND507, ND480 and ND391 respectively (Figure 6B). Out of these 3’-transduction events, 6 were found in somatic insertions, two in ND480 and four in ND391. All fixed or polymorphic germline source elements were full-length L1 sequences. Three of these events were from fixed germline sources (Figure 6C). One source, chr12(q24.32), had previously documented *in vitro* retrotransposition activity^7,50^, while the other two, chr20(p12.2) and chr8(p22.3), had not. Both 3’-transduction events detected from polymorphic germline sources were from the same L1 present on chrX (p22.2), which had both previously documented *in vitro* retrotransposition activity and 3’-transduction events^7,14,50^ (Figure 6D). Interestingly, the somatic L1 with the chrX (p22.2) 3’-transduction found in ND391 was detected in all three regions, while the somatic L1 found in ND480 with the same source was only detected in the cortex of that donor, suggesting this source element is active during more than one stage of human brain development. Another possibility is a highly stable L1 ribonucleoparticles persisting across cell divisions^13^. Moreover, two of the source elements, chrX (p22.2) and chr12(q24.32), have evidence of evolutionarily recent germline activity based on sequence distance from other full-length L1^50^. We additionally detected a somatic L1 on chr5(p13.1) which in turn acted as a source for another somatic event present on chr8(q23.4) (189bp transduction, Figure 6E). These two somatic L1s were significantly overlapping across nuclei (*p* = 2.1 × 10^−8^, Fisher’s exact test). Finally, we combined the genealogical tree, mutation dating, region information and transduction detection to build an overview of the somatic L1 hierarchy as informed by LUSTRE in the three studied individuals (Figure 6F).

## DISCUSSION

Genomic mosaicism is introduced by L1 retrotransposition in human tissues, particularly the human brain. However, critical understanding of somatic L1 activity have been hampered by the limited scalability or sensitivity of existing methods. We developed LUSTRE, a tagmentation-based method that uses few consensus bases in the DNA to detect insertion sites at previously unknown loci, which we demonstrate at a scale previously unachieved in the human brain.

The rates of retrotransposition present in the human brain has been debated largely due to differences in protocols, analyses stringency or the size of datasets. Moreover, analyses have been limited in throughput. We applied LUSTRE to ∼17,500 single neuronal nuclei from the adult human brain across three donors and detected hundreds of somatic L1 events across cells in each donor. We subjected a subset of those L1 events to validation with PCR. Since LUSTRE directly amplifies the 3’ end of the L1 without prior whole genome amplification, DNA was extracted from a separate area of the same tissue section which may have affected the validation results of rare L1 events. Additionally, the limited input of high molecular weight DNA where rare clones contribute few or no template molecules and poor amplification of longer or full-length L1s compared to the shorter wild-type allele may limit our calculated retrotransposition rates to a lower bound. Similarly, because 5’ truncation does not affect 3’ detection design of LUSTRE and single copy loci are efficiently detected, the somatic event detection probability may be overestimated when normalized using polymorphic germline L1s. These effects may bias the overall somatic retrotransposition rate estimates, indicating that our reported rate of 0.1-0.3 somatic L1s per neuron is conservative. While the number of events found per cell is in between previous substantially deviating estimates, the sheer number of cells sequenced with LUSTRE allowed us to detect many more retrotransposition events within the individuals we profiled than previously described. This confirms the hypothesis that L1 does introduce substantial genomic mosaicism in the human brain. We found no large differences in retrotransposition activity across regions, and many somatic L1 events private to a specific brain region, suggesting retrotransposition is active across the brain during development. Our timing inference uses a coalescent model whose assumptions on population dynamics may approximate neurodevelopment, the two-wave pattern we demonstrate was robust to the structured and constant-rate scenarios we tested. However, this remains to be a model-based interpretation which would be strengthened by developmental timing studies where more developmental population structures are included.

Several lines of evidence support the biological origin of the detected L1 events in addition to validation by long-range PCR, as the molecular design of LUSTRE further mitigates the risk of chimeric artifacts. The biological signatures presented in this study demonstrate the L1 endonuclease motif at detected insertion sites, concordance with replication fork directionality, 3’-transductions from known active source elements, including a somatic-to-somatic retrotransposition event. These signatures are collectively consistent with the known mechanisms of retrotransposition and not expected from stochastic chimeric rearrangements. Notably, the core conclusions of this study are based on somatic L1s shared across multiple nuclei and independent sample plates, where several validation approaches are applicable.

Importantly, LUSTRE enabled us to investigate specific aspects of L1 biology. For example, we found that the detected somatic L1 preferentially inserts in the same orientation as the leading strand during DNA replication. This finding supports the claim that somatic L1 retrotransposition is coupled to cell division *in vivo*, and we anticipate LUSTRE will be a useful tool to study these dynamics further. 3’-transductions reveal the genomic origin of retrotransposition and allow us to better understand how retrotransposition is regulated in humans. Previous studies have either studied retrotransposition potential from germline 3’-transduction events or using *in vitro* assays^14,51,52^. In our data, 2 germline sources had been reported to be active according to previous studies, while 2 sources had not been^53^. Interestingly, the two previously reported source L1s had widely different reported activity. In fact, while the source insertion present on chrX is highly active in *in vitro* assays and in the germline, the chr12 source had only been reported to be weakly active *in vitro* by one study. If *in vitro* studies were representative of *in vivo* retrotransposition potential, the likelihood of observing this source amongst all other possible sources would be low. It is likely that the activity of individual L1s is dependent on epigenetic factors which change with cell type, possibly enabling the retrotransposition by L1s whose 3’-transductions are not found in the germline or by *in vitro* assays^12^. Another explanation is allelic variation among the population, where the same L1 locus may appear active or inactive based on the assayed allele^24,43^. This is previously demonstrated in the chr13D31L1 locus, where three allelic variants differ in retrotransposition activity due to the presence or absence of an intact ORF, indicating that an inactive locus *in vitro* may be retrotransposition competent *in vivo*. Such allelic differences therefore provide a basis for the retrotransposition activity of the chr12 source element, which had only been reported as weakly active previously. In contrast, the source present on chrX was found to give rise to 3’-transduction across donors and was either region restricted or across all regions, suggesting its activity is resistant to global epigenetic changes.

The polyA tract homopolymers harbored by L1s are readily detected by LUSTRE and presumably mutate frequently during cell division due to polymerase slippage^42^. This is particularly the case for somatic events, which initially contain much longer tracts than the germline events. However, during library preparation this polymerase slippage also occurs during PCR and linear amplification^54,55^, and the analysis is further complicated by the difficulty of current sequencing technology to sequence polyA tracts with many repeat units^56,57^. To overcome this, we developed two bioinformatic approaches to account for this technical variability. We found that the polyA tracts of somatic L1s in the dentate gyrus are shorter than the same tracts in the cortex and cerebellum, indicating a higher number of cell divisions among the dentate gyrus lineages. This may reflect the neurogenic program that forms the dentate gyrus, where granule neurons are generated from successive rounds of intermediate progenitor proliferation continued into postnatal life^58,59^, increasing the number of divisions that separate the sampled neurons from their ancestral L1 event. Ongoing adult neurogenesis in the dentate gyrus^39,40,60^ may suggest additional divisions in postnatal born neurons in a small subpopulation of granule neurons given the age of sample donors. Therefore, the overall length of shorter polyA tracts in the dentate gyrus is likely to be driven by the developmental program, with adult neurogenesis being a region-specific contributor. Furthermore, the observed polyA tract differences are compatible with multi-base-pair deletions over tens of cell divisions, consistent with previous studies of long polyA tracts^20,44^. While these results are based on aggregate estimates across nuclei, this does suggest there is valuable information in these data to investigate the phylogeny and cell depth of populations of individual cells. However, multiple improvements to the molecular biology and computational analysis may be required to unlock these insights to decrease and account for the technical variation in polyA tract length estimates of individual cells.

While this study focused mainly on neurons, LUSTRE can be applied to other tissues in the human body. For example, previous studies have reported extensive L1 retrotransposition in normal colorectal epithelium^25^. They are particularly implicated in cancer biology, where L1 is known to be retrotranspose in about half of cancers^15^. LUSTRE would enable the study of L1s in individual cancer cells at much larger scales than previously achievable. Additionally, the LUSTRE protocol can easily be modified to target any sequence where the integration site is unknown, for example other transposable elements or viral integration sequences in gene therapy applications. We envision LUSTRE to be a readily applicable and scalable method to detect integration sites in single cells. Collectively, our work further highlights the intrinsic properties of L1s for high-resolution lineage analyses in humans, where changes in polyA tract lengths represent a function of time over an evolutionary scale, and transduced sequences point to the parent and offspring relationships between a subset of L1s and their progeny.

## STAR METHODS

### KEY RESOURCES TABLE

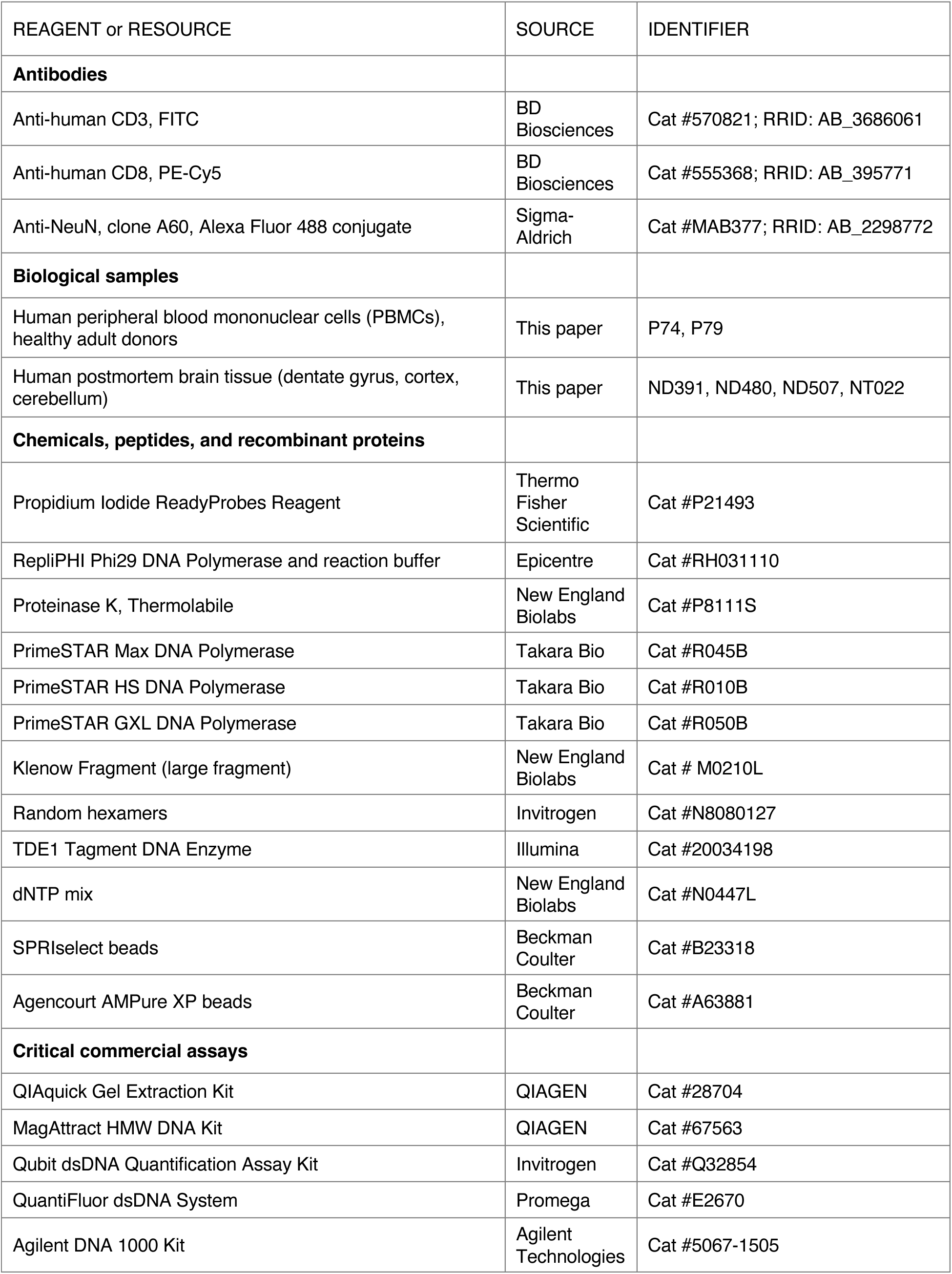
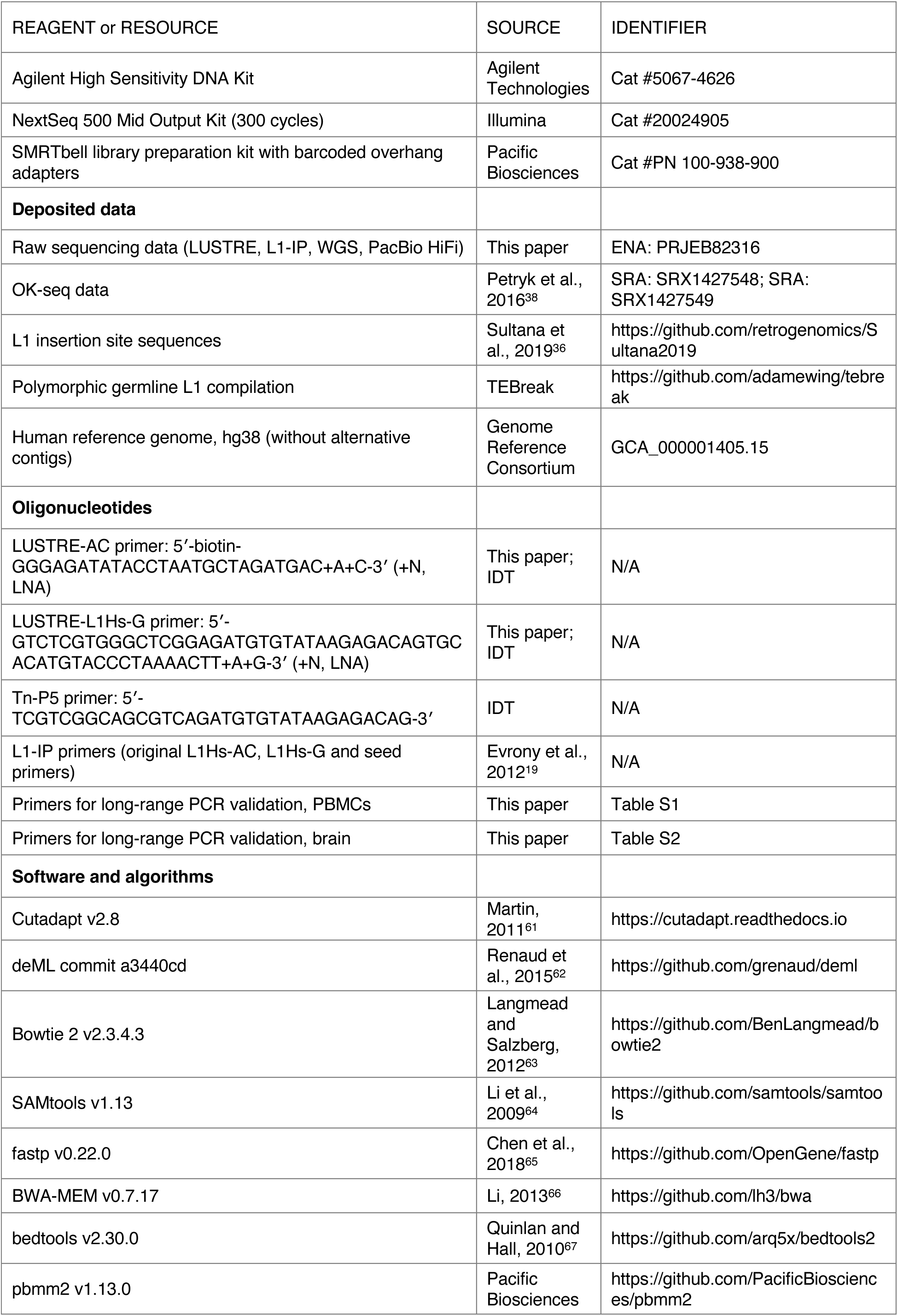
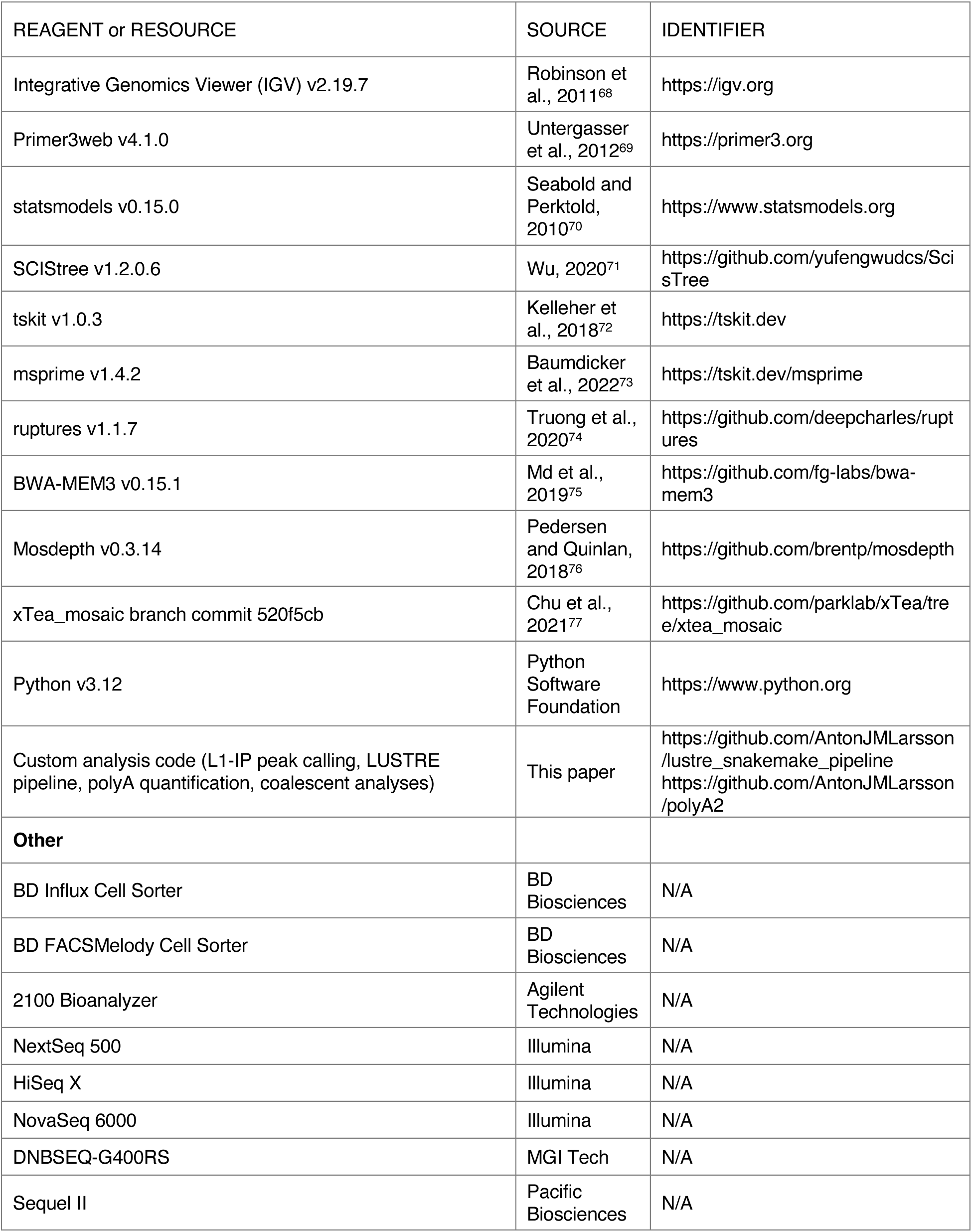

### RESOURCE AVAILABILITY

#### Lead contact

Requests for further information and resources should be directed to and will be fulfilled by the lead contact:

Jonas Frisén

#### Materials availability

This study did not generate new unique reagents. Sequences of all primers generated in this study are provided in the key resources table and in Tables S1 and S2.

#### Data and code availability

- Sequencing data have been deposited at the European Nucleotide Archive (ENA) under accession number PRJEB82316 and are publicly available as of the date of publication.
- All original code has been deposited at GitHub and is publicly available as of the date of publication. The information is listed in the key resources table.
- Any additional information required to reanalyze the data reported in this paper is available from the lead contact upon request.

### EXPERIMENTAL MODEL AND STUDY PARTICIPANT DETAILS

#### Human peripheral blood mononuclear cell samples

Peripheral blood samples were obtained from two healthy male donors aged 41 and 46 after informed consent. Collection of blood samples was approved by the Swedish Ethical Review Authority (2006/229-31/3, 2016/1415-32 and 2020-02604). Mononuclear cells were isolated by density centrifugation and cryopreserved in fetal bovine serum (FBS) with 10% DMSO and stored in liquid nitrogen for subsequent flow cytometry sorting.

#### Human brain samples

Adult dentate gyrus, cortex and cerebellum samples were from a postmortem male donor aged 26 (ND507), female donor aged 25 (ND480) and male donor aged 26 (ND391). A fourth male donor (NT022) aged 24 was used as a control for WGS validation experiments. The tissue was obtained from the Swedish National Board of Forensic Medicine through the KI Donatum tissue collection program. The collection process and all subsequent experiments were approved by the Swedish Ethical Review Authority (2010/313-31/3, 2012/699-32, 2016/2337-32, 2017-1129-32) as well as the ethical committees of the above-mentioned tissue collection bank. The tissue was collected in agreement with national law in Sweden and included informed consent. Tissue samples were dissected immediately after the autopsy, frozen on dry ice upon collection and stored at -80 °C until further processing. Due to the small number of donors, the influence of sex on somatic L1 retrotransposition could not be assessed.

### METHOD DETAILS

#### Flow cytometry analysis and sorting

Frozen PBMC samples were thawed, washed in FACS buffer (PBS, 2% FBS, 1 mM EDTA) and filtered through a 40 µm strainer. Cells were stained with CD3-FITC (BD Biosciences) or CD8-PECy5 (BD Biosciences) together with dyes for live/dead cell discrimination (DAPI or Propidium Iodide ReadyProbes Reagent (Thermo Scientific)). Cells were sorted into 96- or 384-well plates using a BD Influx Cell Sorter or BD FACSMelody instrument with a 100 µm nozzle.

Human brain samples were examined under a stereomicroscope, and 1 mm^3^ pieces from the respective brain regions were dissected. The dissected tissue pieces were homogenized using a glass Dounce homogenizer, in 10 ml of ice-cold lysis buffer (0.32 M RNase and DNase free-sucrose, 5 mM CaCl2, 3 mM Magnesium Acetate, 2.0 mM EDTA, 10 mM Tris-HCl pH 8.0, 0.1% Triton X-100 (Sigma), 1 mM DTT (Sigma) and nuclease-free water). The homogenized tissue solution was then resuspended in 20 ml of ice cold 1.8 M sucrose solution (3 mM Magnesium Acetate, 10 mM Tris-HCl pH 8.0 and Nuclease-free water). The tissue homogenate was slowly layered onto a cushion of 10 ml of the above-described sucrose solution and was centrifuged at 26,500 × g for 2 hours at 4°C. After centrifugation, the supernatant was discarded, the pellet in each tube was resuspended in 1.5 ml of nuclei resuspension buffer (1X PBS, 0.5% BSA) and was filtered through a 70-μm strainer. After washing, the nuclei pellet was resuspended in 100 µl of nuclei resuspension buffer. The quality and purity of the nuclei were evaluated visually using a cell culture light microscope and the concentration of nuclei in the solution was determined with the help of a glass hemocytometer and coverslip. The purified nuclei were incubated for one hour with anti-NeuN Antibody, clone A60 from Sigma-Aldrich conjugated with Alexa Fluor 488 fluorophore (1:500) and for 5 minutes with 0.1 µg/mL DAPI in PBS containing 0.5% bovine serum albumin. Single-nucleus sorting was performed on a BD Influx Cell Sorter with a 100 µm nozzle. All steps were performed on ice, at low temperature.

#### Generation of L1 insertion profiling (L1-IP) libraries

L1-IP libraries were generated according to a previously described protocol^19^. This method requires whole-genome amplified DNA material from single cells as input. Therefore, we have performed multiple displacement amplification (MDA) according to the protocol in the same publication. Briefly, single-cell FACS was performed to sort T cells into 96-well plates containing 2.8 µl lysis buffer (200 mM KOH, 5 mM EDTA, 40 mM DTT) in each well and neutralized with 1.4 µl neutralization buffer (400 mM HCl, 600 mM Tris pH 7.5). MDA was performed by adding 15.8 µl of reaction mix, containing 1X RepliPHI phi29 reaction buffer (Epicentre), 50 µM random hexamers, 2 mM dNTPs, 40 U RepliPHI phi29 polymerase (Epicentre) and H_2_O. The reaction was carried out for 10 hours at 30 °C, followed by heat inactivation for 10 min at 65 °C. After quality control, the samples were stored at -20 °C. Upon thawing, the MDA products were diluted to 100 ng µl^−1^ to be used in PCR. Genomic DNA from PBMCs was used as a bulk sample at sufficient concentration for L1-IP. First PCR of L1-IP involves the pairing of a human-specific L1 primer (L1Hs-AC) complementary to L1 3’ UTR and 8 arbitrary seed primers containing 6-base barcodes along with 5 degenerate N nucleotides. Therefore, each sample was divided into 8 wells of 96-well plates, and one set of seed primers was used for each sample. After 5 cycles of linear amplification with L1Hs-AC, seed primers were added into each reaction to enable exponential amplification of samples. Next, the PCR products were purified with SPRI Ampure XP beads with a 1:1.8 ratio. The second round of PCR involves the pairing of L1Hs-G primer complementary to L1 3’ UTR and an oligo complementary to seed sequences which incorporates an Illumina sequencing adapter. Libraries were subsequently visualized by gel electrophoresis, and size selection was performed by excising bands between 200-500 bp. Gel bands were cut using sterile scalpels for each sample set (8 seed primer products). DNA was extracted in individual columns by using a gel extraction kit (QIAGEN) according to manufacturer’s instructions. Each sample set consisting of 8 seed reactions were pooled equimolarly and DNA concentrations were measured using Qubit dsDNA Quantification Assay Kit (Invitrogen). Library quality and size distribution were checked on a DNA 1000 chip. In each sequencing library, 32 single-cell libraries (32 cells x 8 seeds = 256 samples) were pooled equimolarly and sequenced at 150-bp single-end using Nextseq Mid Output Cartridge kits and a Nextseq 500 sequencer (Illumina).

### LUSTRE

Single cells were sorted into 96-well or 384-well plates containing 3 µl lysis buffer (60 mM Tris-Ac pH 8.3, 2 mM EDTA pH 8, 15mM DTT) and immediately spun down for storage in -80 °C. If nuclear separation was previously performed, nuclei lysates were completed with 2 µl of the lysis buffer upon thawing. Samples were incubated at 75 °C for 30 min. Next, 0.5 µl of thermolabile Proteinase K (1:1000 dilution, NEB) was added to samples for incubation at 37 °C for 15 min, followed by 55 °C for 10 min and immediately placed on ice afterwards. Linear amplification was performed by adding 7 µl of PCR mix, containing 1X PrimeStar Max (Takara) and 0.5 µM LUSTRE-AC primer (5’-biotin-GGGAGATATACCTAATGCTAGATGAC+A+C-3’, HPLC desalted; IDT) into 3.5 µl DNA lysate. LUSTRE-AC primer targets human-specific L1 at positions 5930-5931 as previously described^2,14^. We have made modifications to this primer to ensure robust amplification, where 5’ biotinylation and 3’ locked nucleic acid technology provide directionality and inhibit 3’ degradation of the primer. Linear amplification was carried out as follows: 3 min at 95 °C for initial denaturation, 40 cycles of 20 sec at 98 °C, 15 sec at 63 °C, 45 sec at 72 °C and 5 min at 72 °C for final elongation. Samples were purified with SPRIselect beads (Beckman Coulter) to size select for fragments > 300 bp. To generate double-stranded DNA from the linear amplification product, 2 µl of hexanucleotide priming mix containing 1X Buffer 2 (NEB), dNTPs 0.1 mM (NEB), 0.25 µM random hexamers (Invitrogen), 0.125 µl Klenow large fragment (NEB), 0.5 µl H_2_O was added to samples and incubated for 1 hour at 37 °C followed by 20 sec at 75 °C for heat inactivation. Samples were purified again with SPRIselect beads (Beckman Coulter) to size select for fragments > 200 bp. Purified samples were immediately subjected to tagmentation in 10 µl, consisting of 1X tagmentation buffer (10 mM Tris pH 7.5, 5 mM MgCl_2_, 5% DMF) and 0.02 µl TDE1 (Illumina). Tagmentation was carried out for 12 min at 55 °C, followed by the addition of 0.5 µl of 0.2% SDS. The release of Tn5 from the DNA was enabled through incubation for 30 min at 68 °C. Exponential amplification of tagmented DNA was performed by adding 6 µl of PCR mix, containing 1X PrimeStar HS Buffer (Takara), 0.25 mM dNTPs (Takara), 0.04 U µl^−1^ PrimeStar HS polymerase (Takara), 1 µM LUSTRE-L1Hs-G primer (5’-GTCTCGTGGGCTCGGAGAT GTGTATAAGAGACAGTGCACATGTACCCTAAAACTT+A+G-3’, HPLC desalted; IDT) and 1 µM Tn-P5 (5’-TCGTCGGCAGCGTCAGATGTGTATAAGAGACAG-3’, HPLC desalted; IDT).

LUSTRE-L1Hs-G primer targets human-specific L1 at position 6015 as previously described^11,19^ and contains the adapter for Illumina-Nextera Index 1, allowing for paired-end sequencing and dual-indexing. Tn-P5 is complementary to the synthetic oligo inserted by Tn5 transposase upon tagmentation^78^ and contains the adapter for Illumina-Nextera Index 2. Exponential PCR was carried out as follows: 3 min at 95 °C for initial denaturation, 22 cycles of 20 sec at 98 °C, 15 sec at 65 °C, 1 min at 72 °C and 5 min at 72 °C for final elongation. Resulting LUSTRE libraries were purified with SPRI beads (Beckman Coulter) or in-house produced 22% PEG beads with 1:0.8 ratio. Library quality and size distribution were checked on a high-sensitivity DNA chip (Agilent Bioanalyzer). Libraries were quantified using the QuantiFluor dsDNA System (Promega) and normalized to 1 ng µl^−1^ per well.

#### Validation of somatic L1s in genomic DNA

For the PBMC sample, 7 polymorphic germline and 5 putative somatic L1s together with 2 germline L1s were selected for validation. Primers targeting these loci were designed by using Primer3^69^ (Table S1). High molecular weight (HMW) DNA was extracted from 2 million PBMCs from donor P79 using MagAttract HMW DNA kit (QIAGEN) according to manufacturer’s instructions to be used as template for long-range PCR.

For the brain samples, 12 polymorphic germline and 63 putative somatic L1s were selected for validation (Note S1). Primers targeting these loci were designed by using Primer3 (Table S2). For donors ND507, ND480 and ND391, approximately 25 mg of tissue was excised from the cortex and cerebellum samples. 24 mg, 11 mg and 13 mg of tissue were excised from the dentate gyrus samples for ND507, ND480 and ND391 respectively due to availability. For WGS experiments, an additional 30 mg of tissue was excised from the donor NT022. HMW DNA was extracted using the MagAttract HMW DNA Kit (QIAGEN) according to manufacturer’s instructions to be used as template for targeted long-range PCR and WGS.

We used PrimeStar GXL (Takara) for performing long-range PCR as this enzyme performs best for targets of variable size and primers with variable Tm values under identical conditions^79^. Long-range PCR of L1 targets was performed by adding 1X PrimeStar GXL Buffer (Takara), 0.4 mM dNTPs (Takara), 0.5 U PrimeStar GXL polymerase (Takara), 0.4 µM of forward and reverse primers each to 40 ng of HMW DNA, and cycled as follows for PBMC and brain samples respectively: 30 cycles of 10 sec at 98 °C followed by 10 min at optimized temperatures ranging between 66-70 °C for each primer pair used; 32 cycles of 10 sec at 98 °C, 10 min at optimized temperatures ranging between 60-64 °C for each primer pair used, followed by 12 min at 68 °C. PCR products from the brain samples were visualized on 0.8% agarose gels. PCR products from the PBMC samples were visualized on a 1% gel and individual bands were excised by using sterile scalpels. DNA was extracted in columns by using a gel extraction kit (QIAGEN) according to manufacturer’s instructions.

SMRTbell libraries were prepared for the PBMCs by using barcoded overhang adapters according to manufacturer’s instructions (PacBio). 4-10 kb libraries were pooled and sequenced on a SMRT cell using a PacBio Sequel II.

#### Library preparation and sequencing

Dual indexing of LUSTRE libraries was performed by using custom-designed Nextera index primers containing 10 bp indexes at 0.1 µM concentration each, differing with a minimum Levenshtein distance of 2 between any two indices. For LUSTRE libraries, 1X PrimeStar Max (Takara) and 3 µl Nextera indices were added to each well containing 3 µl of DNA, and cycled as follows: 3 min at 95 °C, 12 cycles of 20 sec at 98 °C, 15 sec at 55 °C, 30 sec at 72 °C, and 5 min at 72 °C. Library quality and size distribution were checked on a high-sensitivity DNA chip (Agilent Bioanalyzer).

Experiments involving the optimization of LUSTRE for experimental conditions and PBMC libraries were sequenced at 150-bp paired-end using Nextseq Mid Output Cartridge kits and a Nextseq 500 sequencer (Illumina). Remaining PBMC libraries were sequenced at 200-bp paired- end on a high-output flow cell using a DNBSEQ-G400RS (MGI). The adult dentate gyrus libraries were sequenced 150-bp paired-end on an S4 flow cell using a NovaSeq 6000 (Illumina). Remaining neuron libraries were sequenced 150-bp paired-end using a DNBSEQ-G400RS (MGI). The target number of reads for LUSTRE libraries was selected to 250,000 paired-end reads per cell to provide read coverage for detection of germline and somatic insertions. Final WGS library preparation for short-read bulk WGS and sequencing at a minimum of 60X coverage on DNBSEQ-T7 (MGI) were performed by Xpress Genomics AB, Solna, Sweden.

### QUANTIFICATION AND STATISTICAL ANALYSIS

#### Data processing and analysis of L1-IP data

##### Demultiplexing and mapping

The first 21 bases of the read give information regarding the cell, and which seed primer the read belongs to. The cell barcode is present at bases 5-11, while the seed barcode is present at base 16-21.

*Cutadapt*^61^ was used to identify these sequences and the software *deML*^62^ was used to assign the most likely cell and seed combination. The reads corresponding to the same cell were then identified based on the demultiplexing.

Before mapping, repetitive G sequences present on the end of the read were trimmed with *cutadapt*. The reads were mapped with bowtie2^63^ to a custom version of hg38 where human specific L1 sequences have been N-masked to ensure unbiased mapping. The resulting .sam file was then turned into a .bam file and sorted using *samtools*^64^.

##### Peak calling insertions

The peak calling was done with an in-house script, which is a re-implementation of the peak calling method described by Evrony et al., 2012. Briefly, a peak is started when the read pileup above a given mapping quality on a particular mapping position is above a specified threshold. In our case, and for Evrony et al., 2012, that threshold is 10 reads and a mapping quality of 25. The peak is extended until there are not any more reads covering a position, with an allowed gap up to 500 bp, same as Evrony et al., 2012. Peak statistics were then collected: read depth, strand and unique read start sites.

##### Softclipping reads and insertion site profiling

During the mapping process, the 5’ or 3’ part of the read (or both) is allowed to not match the reference sequence which the read is assigned to. This part of the read (if present) is called the softclipping part of the read. For the L1-IP-seq method, this occurs if we sequence into the human specific L1 site, but there is no support for the insertion in the reference genome. Therefore, we extracted the softclipping sequences from reads and mapped them to the human-specific L1 sequence. If the read maps to the sequence, the peak which that read is a part of has softclip support. Furthermore, it allows us to obtain information on the exact insertion site sequence of that peak.

##### Assessing sensitivity by polymorphic germline insertions

The sensitivity of L1-IP to detect heterozygous insertions was measured by considering polymorphic germline L1s. These polymorphic L1s are predominantly heterozygous, and each polymorphic L1s found in at least one cell or sample was used to estimate sensitivity. The fraction of detection was calculated for each polymorphic germline insertion, and the median fraction of detection across samples was used as a measure of sensitivity.

#### Data processing and read alignment for LUSTRE

The reads were trimmed using fastp^65^ and mapped to the human reference genome (hg38, no alternative contigs) using bwa mem^66^. The first read mate (read 1) contains the flanking sequence, while the second read mate (read 2) contains the 3’ L1 sequence, poly-A tract, potential 3’-transduction sequence and additional poly-A tract, and potentially a portion of the immediate flanking sequence.

To discover L1s from combined single-cell LUSTRE data, the read 2 alignments which serve as anchoring reads were first identified. All the read 2 alignments were collected and subjected to four requirements for further consideration: the alignment is not a supplementary alignment, is paired, is mapped, and the first 23 bases of the forward sequence correspond to the expected L1 sequence up to a Hamming distance of 6 and the 8 following bases correspond to the non-primer sequence AGTATAAT within a hamming distance of 1. These reads were then separated into three different groups: reads within 10kb of a germline insertion, reads within 10kb of a polymorphic germline insertion (based on the compilation distributed with TEBreak; see KEY RESOURCES), and the remaining read clusters (i.e. putative somatic). The polymorphic germline and putative somatic reads were further separated based on whether the read was discordant or not. In this context, a read pair is discordant if any of these 3 conditions are true: the read-pair maps to different chromosomes, the read-pair maps to the same chromosome at least 15 kb away from each other, or the read-pair is properly paired but the L1 portion of read 2 does not map to the loci (i.e. softclipped). The germline insertion group was directly quantified per insertion with no additional requirements. The polymorphic germline and putative somatic insertion groups were further filtered to exclude reads mapping to any position where at least 1% of reads were concordantly mapping. The genomic intervals containing reads were then identified in each cell and merged within 1kb across cells using *bedtools*^67^. The number of unique tagmentation sites (no softclipping and mapping quality above 0) with more than 10 supporting reads were then counted, and all intervals with no such tagmentation sites were discarded. The intervals were then further merged within 2 kb using *bedtools*. Each interval was then analyzed for softclipping reads oriented toward the insertion site, to determine the most likely insertion position. The read intervals were then subjected to further filtering, based on the characteristics of the polymorphic germline insertion group: the mean mapping quality is above 25, at least one cell has over 10 reads detected in the interval, and the consensus sequence of the softclipping portion of reads at the insertion site needs to contain at least 8 As in the first 10 bases of the sequence since this sequence is expected to correspond to the polyA tract. If the detected consensus sequence is shorter than 10 base pairs, all bases in the sequence must be As. Moreover, if the putative somatic L1 call was observed across multiple donors, including other datasets generated through LUSTRE but not included in this study, the interval was discarded.

To investigate whether the insertion site sequence as a group is consistent with previous knowledge, we constructed a position weight matrix (PWM) from insertion site sequence data from another study (data found at https://github.com/retrogenomics/Sultana2019). The insertion site sequences were scored based on this PWM and a Logo plot was made for the putative somatic insertions present in that sample. A logo plot shows the probability of each letter (A, T, C, G) for each position in the insertion site sequence.

Reads which aligned to a long polyA tract instead of the flanking sequence were recovered by extracting the softclipped portions of reads whose aligned sequences contained at least 90% As. The softclipped regions of these reads were then aligned to a custom reference genome containing all sequences 1kb upstream and downstream of all germline and polymorphic germline insertions and unfiltered putative somatic intervals. The aligned reads were not included in the discovery but used in the final quantification for polymorphic germline and putative somatic insertions. For cell-level analyses, an insertion was considered detected in an individual cell if at least 3 supporting reads were found.

In summary, the following characteristics were required for a somatic L1 call:

- Not within 10kb of a known germline insertion.
- Not identified in any other donor across all generated LUSTRE data.
- At least 99% of reads are discordantly mapping.
- At least one identified tagmentation site with >10 reads.
- Mean mapping quality above 25.
- At least one cell barcode with >10 reads detected in the genomic interval.
- The consensus sequence of the called insertion site contains at least 8 As in the first 10 bases of the sequence. If the sequence is shorter than 10 base pairs, all bases in the sequence must be As.

#### Data analysis of L1 insertion sites with long-read sequencing

The HiFi reads were demultiplexed using the known primer sequences together with the ligation adapter sequences to separate reads by sample (Table S1). Reads which did not contain any expected combination of sequences were discarded at this step. Reads were then aligned to the hg38 build of the human reference genome, using pbmm2 with the HiFi alignment mode. The reads of each locus in the experiment were inspected using the software Integrative Genomics Viewer (IGV)^68^ to find large insertions or softclipped reads at the expected insertion site. If found, characteristics of the sequences were collected: TSD, polyA tract length, and allelic mutations by inspecting the reads in IGV and their specific insertion sequence. The allelic mutations were then quantified using a python script.

#### Analysis of replication fork direction bias using OK-seq

For analyzing replication fork direction bias, the OK-seq data was obtained from Petryk et al^38^. Specifically, SRX1427548 and SRX1427549 were used to calculate replication fork direction (RFD) bias. The OK-seq processing was performed exactly as previously described, with the exception that the reads were mapped to the hg38 reference genome instead of hg19. The difference in reference genomes explains the difference in coordinates between Figure 2G in this work and Figure 5A in Flasch et al^37^. The analysis of insertion direction preference as a function of RFD bias was also performed in the same manner as described in Flasch et al., 2019.

#### Criteria for validation for L1 somatic insertions subjected to PCR validation

To calculate the validation rate, we assigned the status of each L1 somatic insertion subjected to PCR validation accordingly:

- PCR validated: PCR supports the existence of a larger DNA fragment than the expected empty-site amplicon. The larger band may be separated from the main band to varying degrees.
- PCR inconclusive: No evidence by PCR that a larger DNA fragment was amplified. The band is consistent with the expected size and there is no second larger band observed.
- PCR missing: No observable band at all or only clear off-target priming.
- IGV: No indication of inconsistencies after manual inspection of read coverage using the Integrated Genome Viewer (IGV). Note that very few insertions fail this criterion.

We then assigned each insertion a combination of PCR validated/PCR inconclusive and IGV status. Four different grouping criteria was then used to calculate a validation rate:

- PCR validated and IGV
- PCR validated (meaning PCR supports the insertions but IGV has indications of inconsistencies)
- PCR validated/PCR inconclusive and IGV
- All PCR (including PCR missing) and IGV

See Note S1 for an overview of all validations, all PCR results for each insertion together with IGV screenshots, and a summary of the classification for each group of criteria.

#### Analysis of bulk neuron WGS

WGS data was trimmed using fastp^65^, aligned using bwa-mem3^75^ to the hg38 reference genome with alternative contigs. The aligned file was then sorted and duplicate-marked using samtools^64^. Coverage depth was calculated using mosdepth^76^. The data was analyzed for somatic L1 discovery using the xTea^77^ software using the mosaic mode.

To assess the WGS support of insertions discovered by LUSTRE, and to avoid tool-specific filtering that may cause false negatives, we used a supervised approach which analyzed clipped and discordant reads in the specific regions. Softclipped reads were indicated to support an L1 event if the breakpoint of the clipped region was consistent with a polyA tract (15 bases 80% A/T) or the sequence otherwise shared at least one 15-mer with the consensus L1 sequence. Discordant reads were considered to support an L1 event if the mate was found mapped to a L1HS region found in the reference genome.

To determine the expected number of WGS reads for a somatic L1, we calculated:

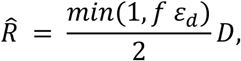

where *f* is the fraction of nuclei the L1 was detected in and *ε_d_* is the detection rate for the donor *d* and *D* is the WGS depth at that region of the library.

#### Sensitivity and validation rate adjusted L1 insertion estimation

Given the number of *detected* somatic insertions and the fraction of detected polymorphic germline insertions, the sensitivity-adjusted number of somatic insertions per cell was estimated as follows. The number of *k* detected somatic insertions was assumed to follow a binomial distribution B(*n, p*) with *n* representing the actual number of somatic insertions and *p* the detection probability. Formulating the problem in this manner, we see that *k* is known and we can use the fraction of detected polymorphic insertions as an estimate of *p*. *n* is estimated by calculating the log likelihood of observing *k* insertions at the nuclei specific detection probability *p* for *n = 0,..,100*, and applying the softmax function to the resulting log likelihood. The range for *n* can be considered a discrete uniform prior with finite support (0 to 100), and *p* is determined using the estimated sensitivity per nuclei. The expected value of this probability was then calculated for each nucleus – corresponding to a sensitivity adjusted estimate. The mean and 95% CI for nuclei in each region and donor was then calculated and visualized.

To adjust this value further by validation rate, the insertions subjected to validation were grouped into “validated” and “non-validated” according to four different grouping criteria for each donor (see previous methods section). The non-validated insertions were dropped from the initial table used to calculate the number of detected somatic insertions, and the fraction of validated insertions was used to scale each nucleus-specific sensitivity adjusted estimate before calculating the mean and 95% CI for each region and donor.

#### Ordinary Least Squares (OLS) with outlier detection to assess region and donor specific L1 detection

The OLS class in *statsmodels*^70^ was fitted to the number of sequenced nuclei and total number of detected somatic insertions, counted separately for each donor and region. The outlier test function of the OLSResults class was used with Bonferroni significance correction, which classified the DG region in donor ND391 as an outlier.

#### L1 mutation dating and lineage tree construction

The software *scistree*^71^ was used to obtain a genotype table based on the single-cell somatic insertion table. This genotype table and resulting lineage tree were then analyzed using *tskit*^72^. The overall population dynamics were estimated using a rejection-sampling approach to the standard Kingman’s coalescent model described previously^80^. To date mutations, another previously described rejection-sampling approach was used with a slight modification to the acceptance step, where the generated estimates are accepted if

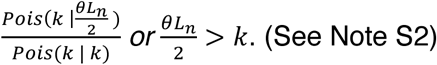

We additionally used the Kingman’s coalescent model to simulate mutation dating estimates given the number of nuclei with the mutation and the total number of nuclei. The procedure was repeated under acceptance windows of the Poisson CDF of 0.025–0.975, 0.05–0.95, 0.1–0.9 and 0.2–0.8 for all three donors, and the effect on the posterior N and θ was assessed. The observed detected-frequency spectrum was compared with the spectrum expected under a single wave of retrotransposition subject to LUSTRE’s detection characteristics. Detection was modelled as binomial thinning. An insertion truly present in *k* cells was detected in *Binomial*(*k*, *p_det_*) of them, with the true frequency spectrum drawn from the neutral coalescent expectation, read depth sampled from the empirical per-cell read-count distribution, and *p_det_* spanning 0.30–0.66. Expected detected spectra were obtained from 400 Monte-Carlo replicates per donor, and the fraction of insertions in the intermediate band (10–100 detected cells) was compared between observed and simulated data. To test whether a constant retrotransposition rate over a non-trivial scenario could reproduce the observed spectrum, genealogies were simulated with *msprime*^73^ for n = 4150 cells (30 replicates each) under six demographies. Constant size, exponential (transit-amplifying) expansion, mild and strong bottlenecks, spatially structured progenitor pools (8 demes with limited migration), and a symmetric to asymmetric division shift, all under a single constant mutation rate. Simulated spectra were passed through the same detection model (*p_det_* = 0.66), and the fraction of insertions in the intermediate band was compared with the observed value.

#### Identification of transduction events

To detect transduction events, all softclipped portions of mapped reads were extracted, trimmed of uninterrupted runs of Ts at both ends and re-mapped to the human reference genome (hg38 without alternative contigs). Moreover, the orientation and mapping quality of the read were propagated for each re-mapped softclipped portion. All re-mapped portions were then analyzed together with all detected germline, polymorphic germline, and putative somatic insertions. A transduction event was considered detected if more than 10 reads with mapping quality above 25 (both for the re-mapped portion and the original read) were found at both the source and target insertion.

#### Quantification of polyA tract lengths

For the germline elements, we developed a computational framework to identify the flanking region and count the length of the polyA tract in between. However, due to the difficulty in sequencing homopolymers with short-read technology of the long tracts harbored by somatic insertions, we employed a change-point detection method to estimate the length of very long tracts based on their base calling quality scores. For germline insertions, the sequence of each read 2 was summarized with polyA runs truncated and scanned for possible allelic variation. Summarized sequences close to allelic variants were then detected and corrected. Then, histograms of polyA tract lengths were calculated for each cell and tagmentation event, and the median tract length across reads was calculated for each cell. Due to the drop-off in sequencing quality of long polyA tracts found in somatic insertions, the above strategy was not used for somatic insertion events. Instead, we used the drop-off in sequencing quality after the polyA tract to quantify its length. We applied a binary segmentation algorithm (Binseg) using the python package *ruptures*^74^, with model ‘l1’ and jump size 1. If the algorithm found no change-points or the number of detected As in the sequence was below 85%, we did not keep the estimate. If the read passed these criteria, we saved the length of the sequenced tract, and the median value across all quantified reads per cell was considered the estimated length.

## ACKNOWLEDGEMENTS

This work was supported by grants from Göran Gustafsson Foundation (R. S.), the Strategic Research Program in Stem Cells and Regenerative Medicine at Karolinska Institutet StratRegen (J. F.), the Knut and Alice Wallenberg Foundation (J. F.), the Swedish Cancer Foundation (R. S., J. F.) and the Swedish Research Council (J. Michaëlsson, R. S., J. F.).

## AUTHOR CONTRIBUTIONS

I. Demirci and J. F. conceived the project. I. Demirci performed the experiments, prepared the sequencing libraries, analyzed the data, interpreted the results and wrote the manuscript. A. J. M. L. developed the data analysis pipelines for LUSTRE, analyzed the data, interpreted the results and wrote the manuscript. M.P. and I. Dumitru performed the human brain dissections with help from I. Demirci. M. H. J. assisted with LUSTRE method development, final library preparation and sequencing. K. A. and H.D. performed the tissue excisions in the postmortem donors. C. Z. assisted with final library preparation and sequencing. J. Michaëlsson and J. Mold isolated peripheral mononuclear cells. R. S. and J. F. interpreted the results and wrote the manuscript. J. F. supervised the study.

