## Supplemental Figures for "Dynamics of somatic L1 retrotransposition in the human brain"

### Supplemental Figures 1-12

#### Figure S1

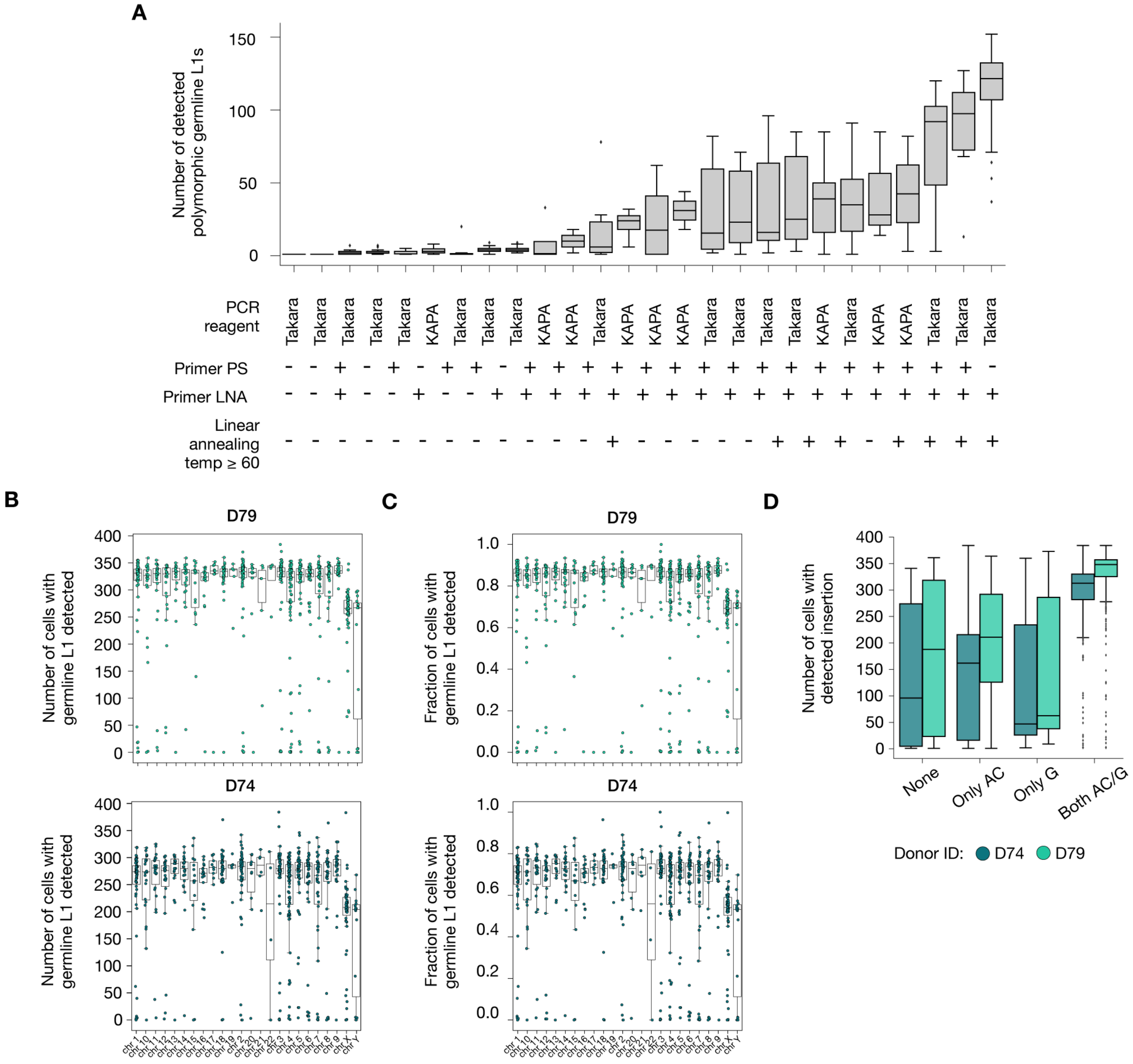


**Figure S1. Method development and detection of diagnostic nucleotides for L1Hs.**
(A) Sequenced experimental conditions of LUSTRE based on key changes to the reagents and protocol parameters.
(B) Number of cells with germline L1s detected with the diagnostic nucleotides (AC/G) for donor D79 (top) and D74 (bottom), separated by chromosome of origin.
(C) Fraction of cells with germline L1s detected with the diagnostic nucleotides (AC/G) for donor D79 (top) and D74 (bottom), separated by chromosome of origin.
(D) Number of cells with a given insertion detected, grouped by the diagnostic nucleotides specific for L1HS-Ta subfamily for donor D74 and D79 respectively.

**Figure S2**

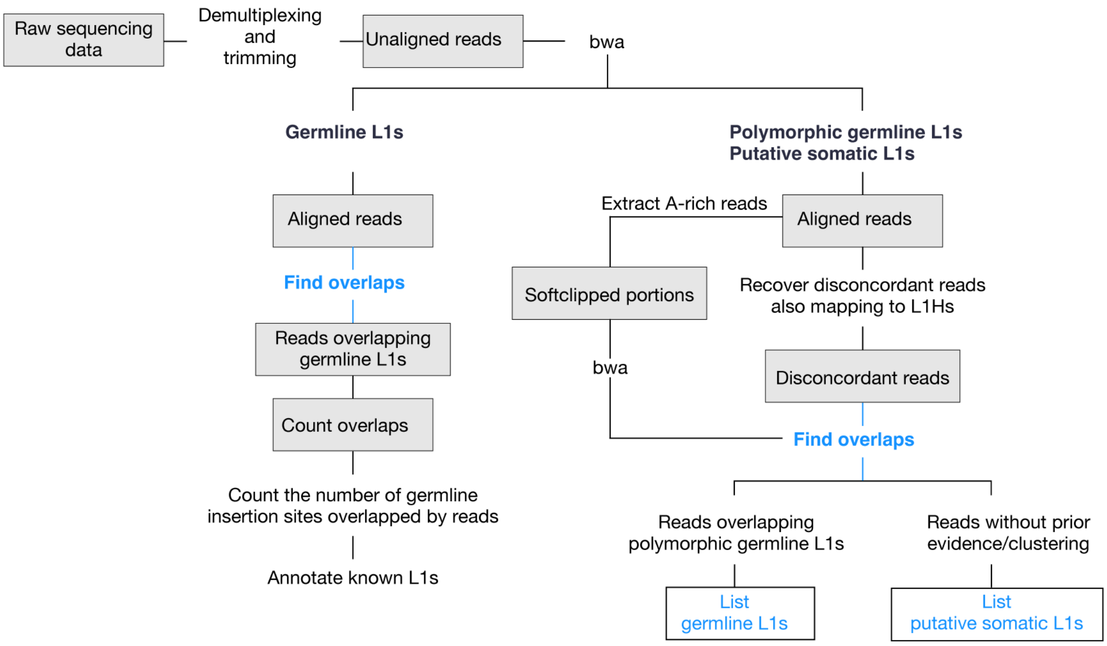


**Figure S2. Overview of the computational strategy used to detect L1Hs from single-cell sequencing data.** See STAR Methods.

**Figure S3**

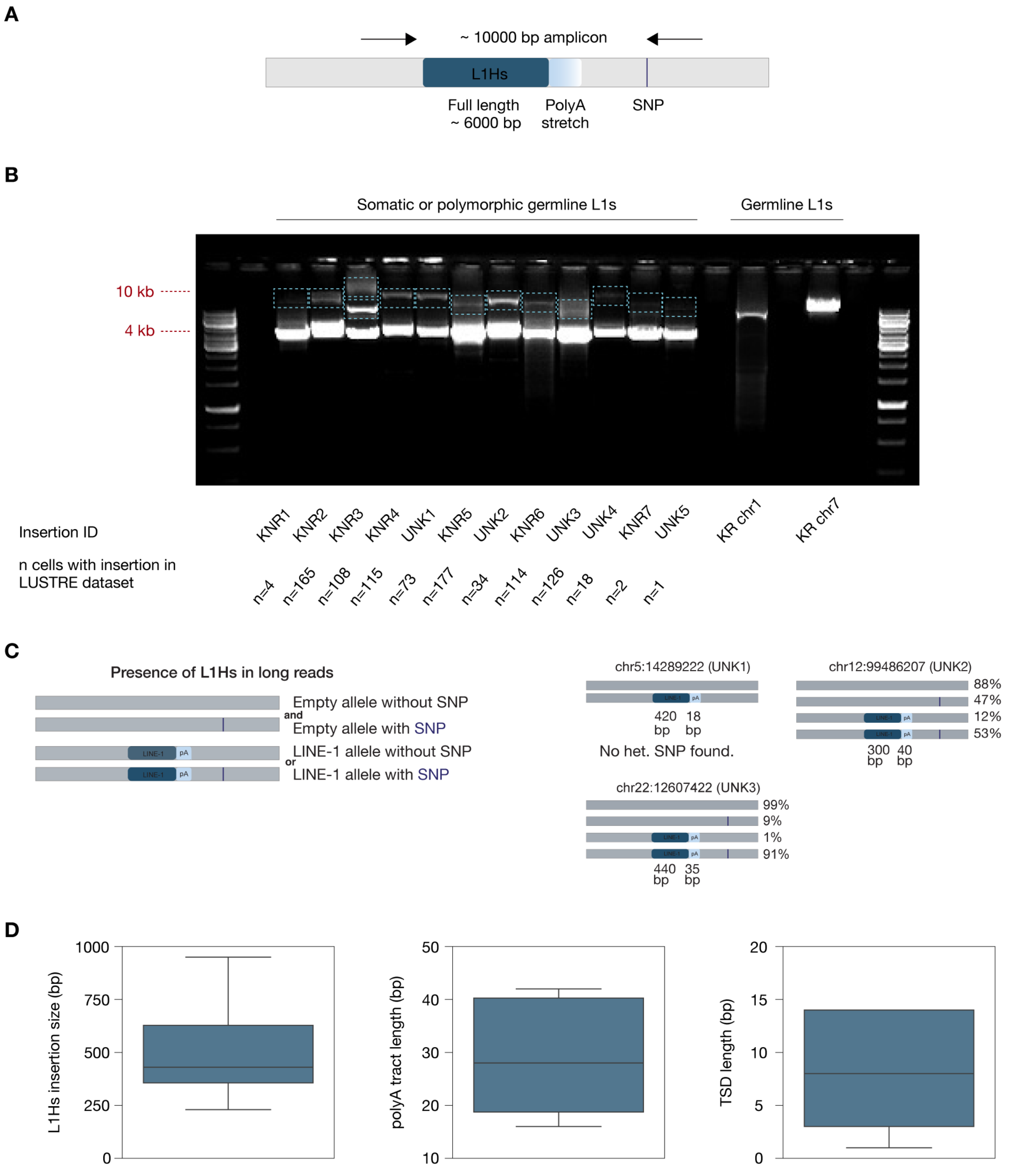


**Figure S3. Validation of putative somatic L1s using long-read sequencing.**
(A) PCR strategy for long read amplicon sequencing, 40 ng of HMW DNA was used for PCR.
(B) Image of gel electrophoresis of PCR amplified putative somatic (UNK) and polymorphic germline (KNR) L1s, including positive and negative controls. Labels highlighting the top gel bands indicate the PCR products used for PacBio sequencing to validate polymorphic or somatic L1s found in corresponding long read sequencing data: KNR2, KNR4, UNK1, UNK2, KNR6, UNK3.
(C) Validating the presence of putative somatic L1s using long-read sequencing and supporting heterozygous SNPs.
(D) Summary of insertion size, polyA tract length, and target-site duplication (TSD) size for long-read validated L1s.

**Figure S4**

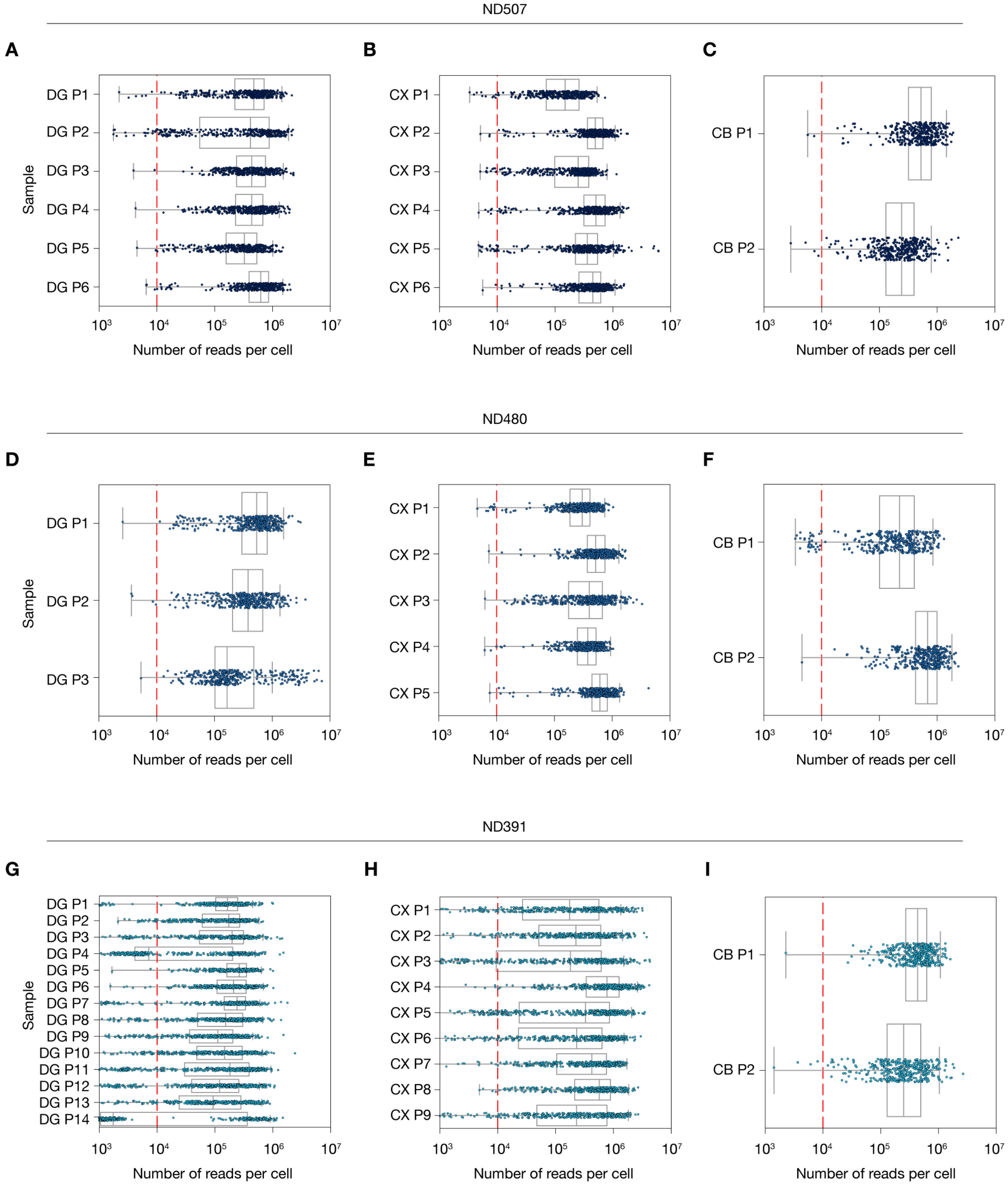


**Figure S4. Sequencing depth of nuclei for human brain datasets.**
(A-I) Number of reads assigned to each single nuclei separated by plate of origin and region for donor ND507 (A-C), ND480 (D-F), and ND391 (G-I). Dashed red line indicates 10,000 reads.

**Figure S5**

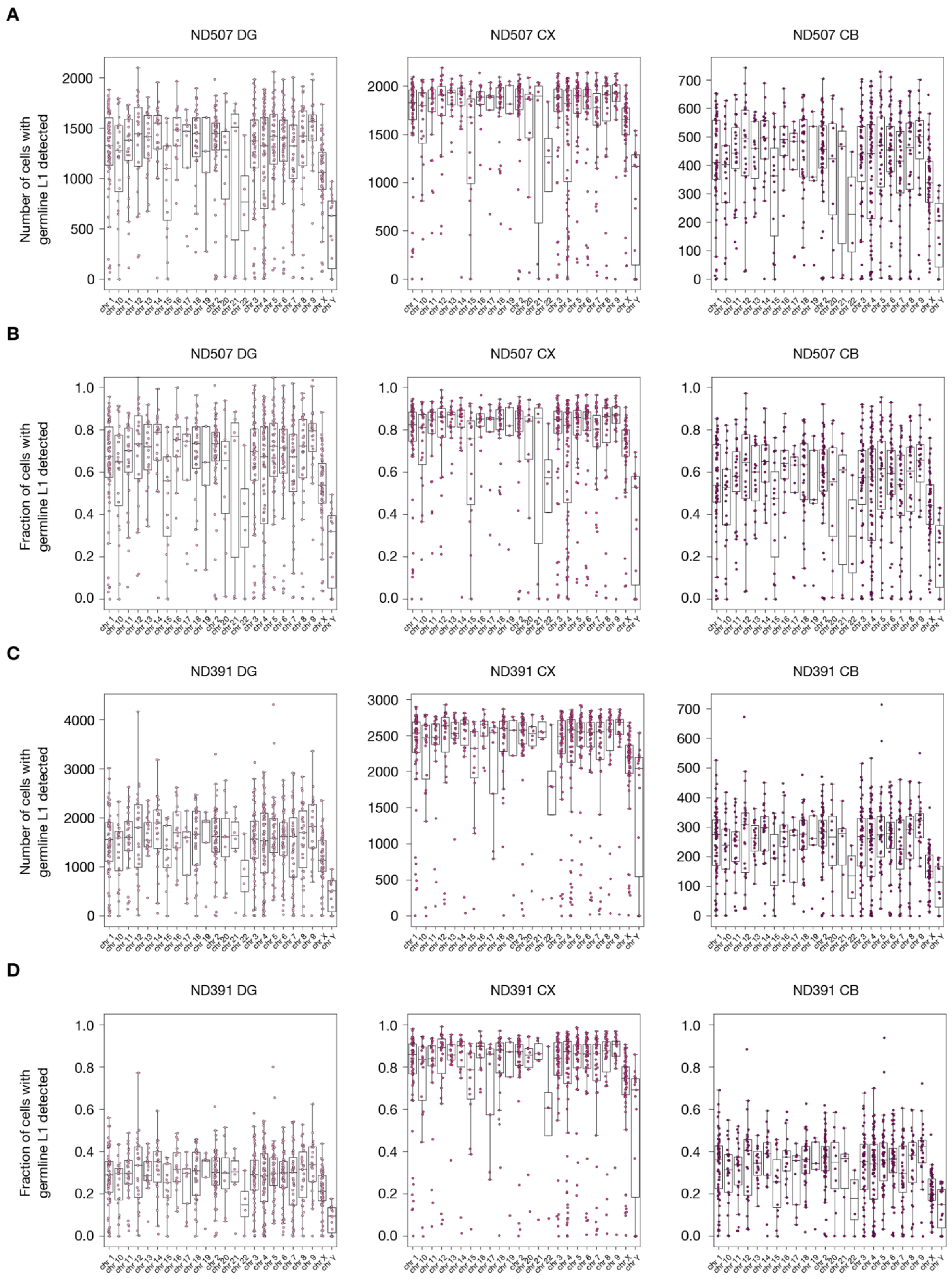


**Figure S5. Number and fraction of germline L1 in neurons with the diagnostic nucleotides (AC/G) in male donors.** ND507 (A and B) and ND391 (C and D), separated by chromosome of origin where the X and Y chromosome exists in only one copy. ND480 was not analyzed in this manner since this donor was female.

**Figure S6**

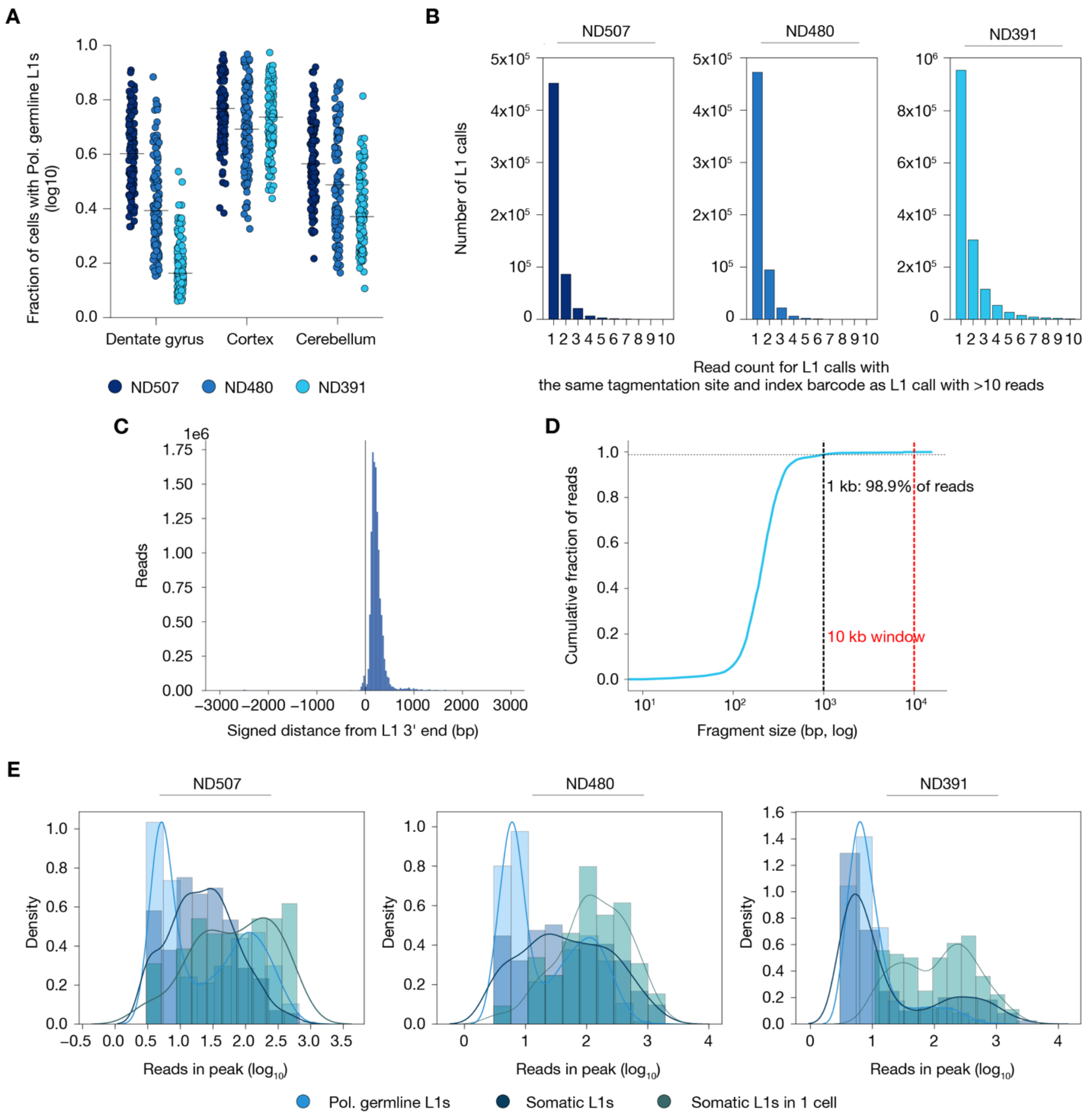


**Figure S6. The relationship between cells sequenced and L1Hs detection.**
(A) Fraction of polymorphic germline L1Hs detected per nuclei, separated by region and colored by donor.
(B) Read count for L1 calls which share a tagmentation site and one index barcode (reads are dual-indexed) with another L1 call which have >10 reads. The majority of these calls have < 3 reads and are filtered away by a read cutoff. Shown separately for ND507 (right), ND480 (middle) and ND391 (left).
(C) The signed distance of reads from annotated L1 3' ends gathered from 10kb windows surrounding known germline L1s across the genome.
(D) The cumulative distribution of read fragment sizes, showing that the vast majority of reads are within 1kb of the annotated L1 3' end within the defined 10kb windows (see Methods).
(E) Mean read depth for polymorphic germline L1, somatic L1, and somatic L1 only detected in one cell. Shown separately for ND507 (right), ND480 (middle) and ND391(left).

#### Figure S7

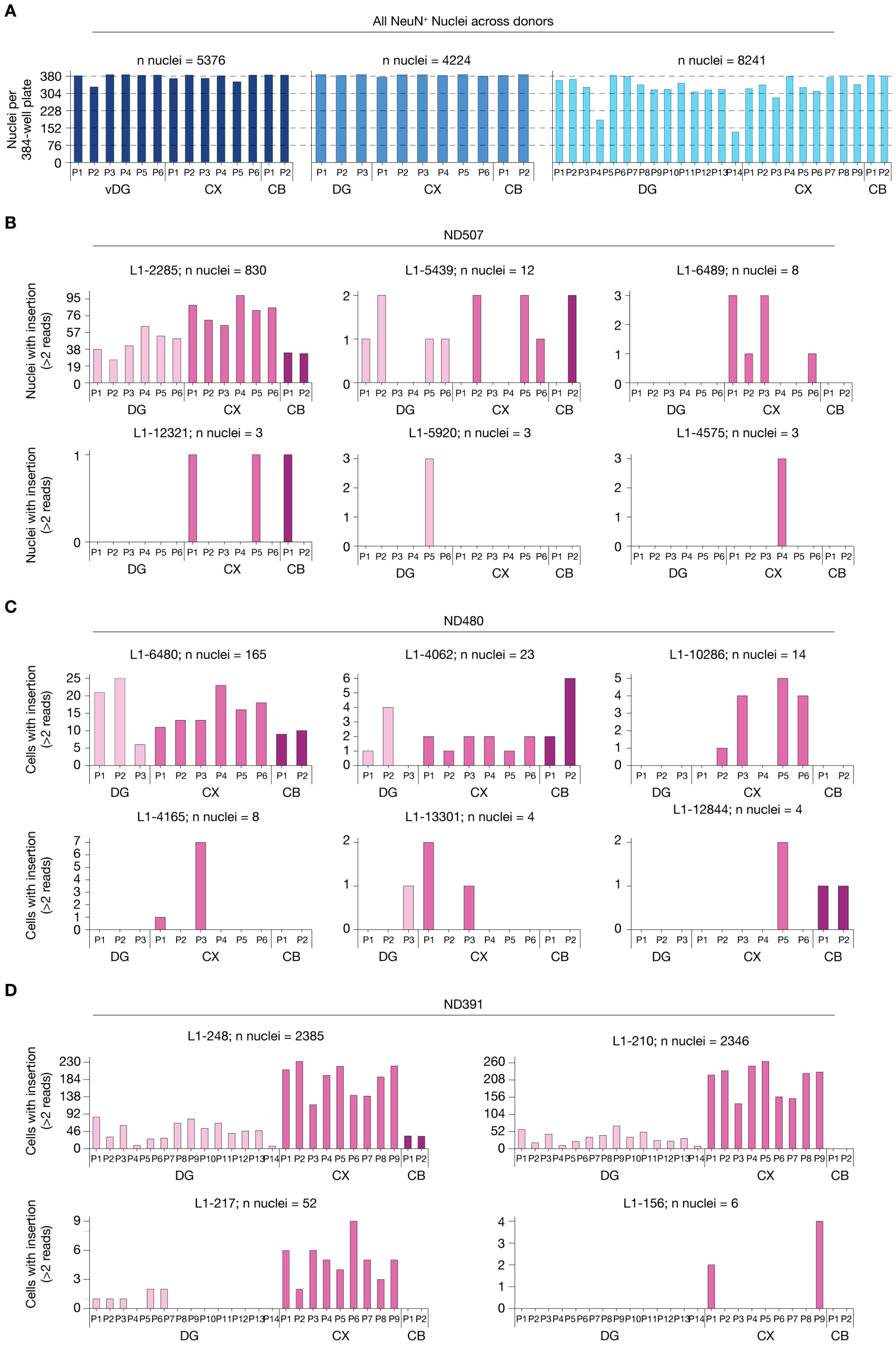
A
All NeuN+ Nuclei across donors
n nuclei = 5376
n nuclei = 4224
n nuclei = 8241
Nuclei per 384-well plate
380
304
228
152
76
0
vDG
CX
CB
DG
B
ND507
L1-2285; n nuclei = 830
L1-5439; n nuclei = 12
L1-6489; n nuclei = 8
L1-12321; n nuclei = 3
L1-5920; n nuclei = 3
L1-4575; n nuclei = 3
Nuclei with insertion (>2 reads)
C
ND480
L1-6480; n nuclei = 165
L1-4062; n nuclei = 23
L1-10286; n nuclei = 14
L1-4165; n nuclei = 8
L1-13301; n nuclei = 4
L1-12844; n nuclei = 4
Cells with insertion (>2 reads)
D
ND391
L1-248; n nuclei = 2385
L1-210; n nuclei = 2346
L1-217; n nuclei = 52
L1-156; n nuclei = 6
Cells with insertion (>2 reads)
DG
CX
CB

**Figure S7. Distribution of somatic L1s across brain regions and sequenced plates.**
(A) Number of cells included in analysis by plate, donor and region.
(B-D) Detection of selected somatic L1s in donor ND507 (B), ND480 (C) and ND391 (D) separated by plate and region. DG: Dentate Gyrus, CX: Cortex, CB: Cerebellum.
See **Figure S6B** for the justification to select read cutoff >2 reads.

**Figure S8**

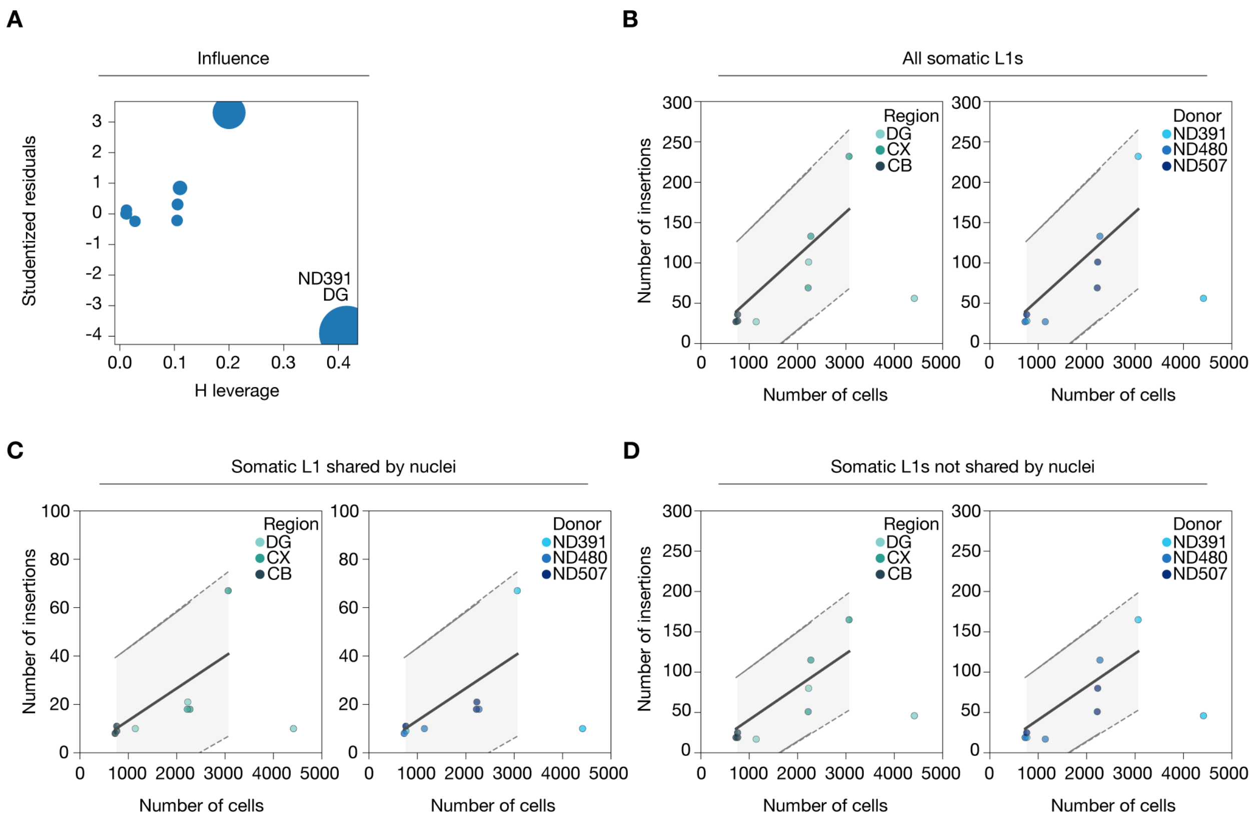


**Figure S8. The relationship between nuclei sequenced and L1 detection.**
(A) Influence plot of ordinary linear squares regression of number of sequenced nuclei and somatic L1 detection for each region and donor.
(B-D) Ordinary linear squares regression using all L1s (B), only L1s shared across nuclei (C) or L1s found in one nuclei (D).

**Figure S9**

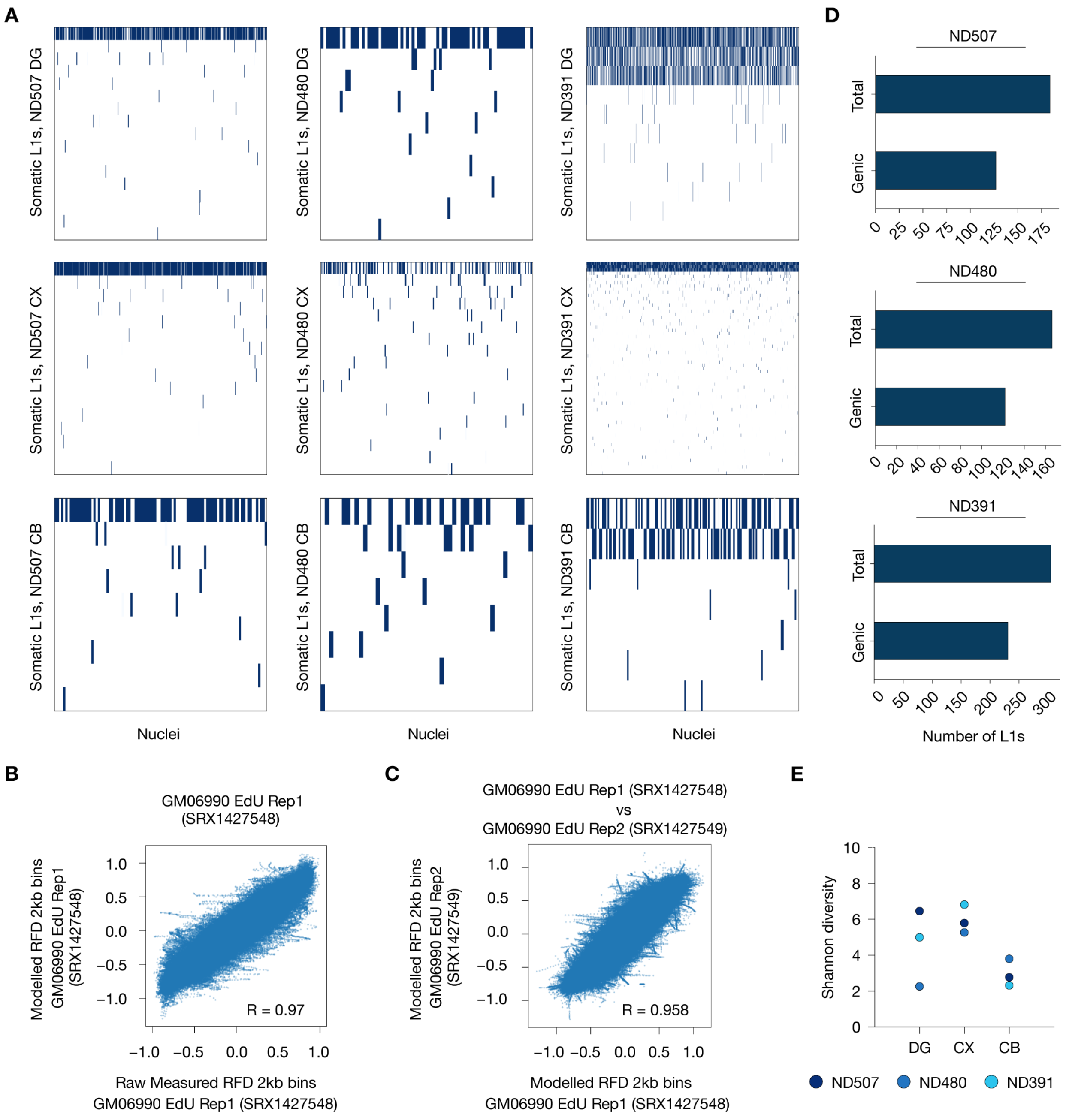


**Figure S9. Somatic L1 location, distribution and insertion preference.**
(A) Heatmap of somatic L1Hs detected in each single nuclei per region (DG: dentate gyrus top, CX: cortex middle, and CB: cerebellum bottom) and donor (ND507 left, ND480 middle, ND391 right).
(B) Correlation between modelled and raw measurements of replication fork direction (RFD) bias for the GM06990 cell line (SRX1427548, obtained from Petryk, N. et al., see References for complete citation).
(C) Correlation between replicates of modelled replication fork direction (RFD) bias for the GM06990 cell line (SRX1427548 and SRX1427549, obtained from Petryk, N. et al., see References for complete citation).
(D) Number of somatic L1Hs found in genic regions (introns) compared to the total number of insertions for donor ND507 (left) ND480 (middle) and ND391 (right).
(E) The estimated Shannon diversity index based on somatic insertion nuclei counts for each region and donor.

**Figure S10**

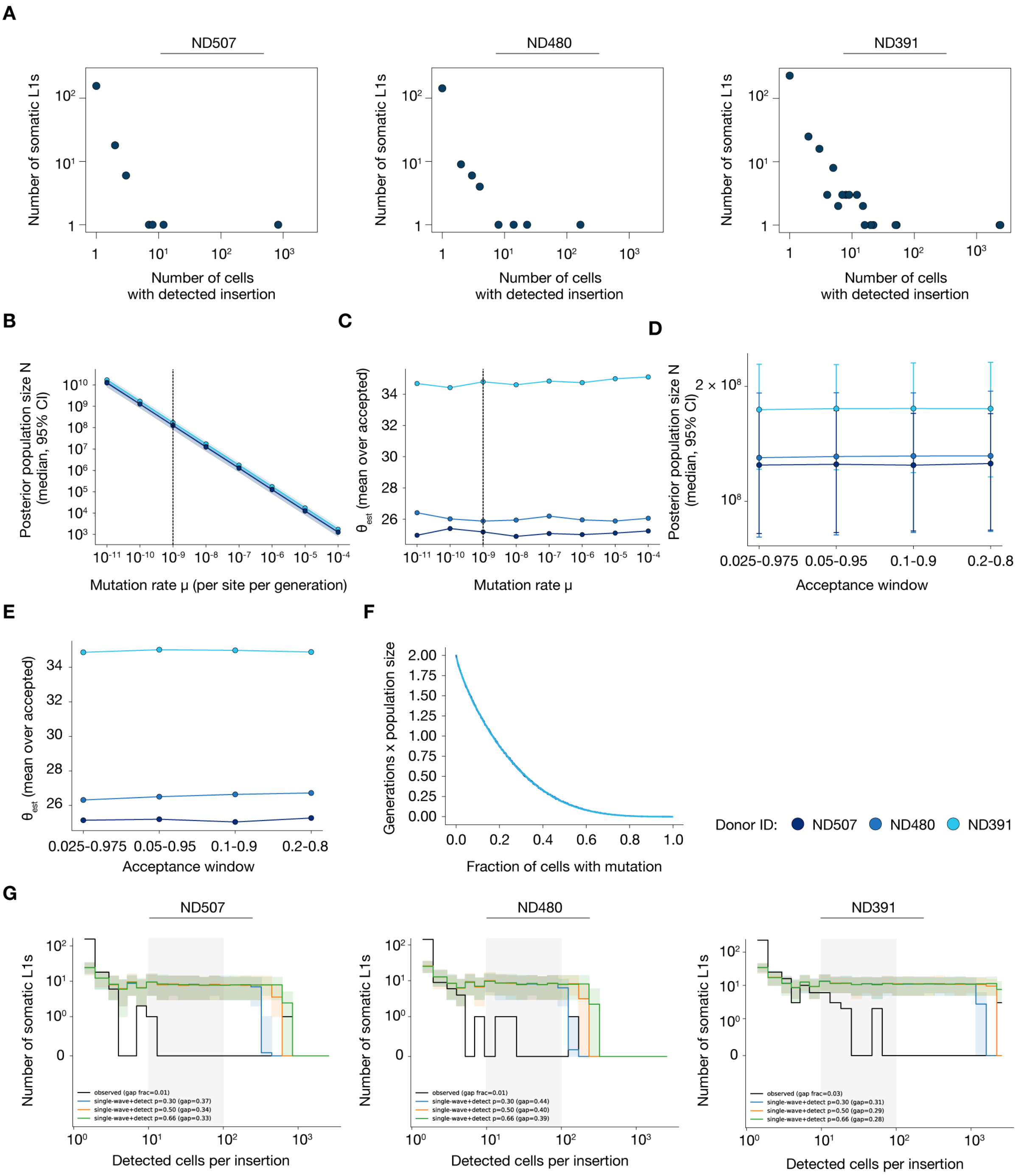


**Figure S10. Simulated mutation dating using the Kingman's coalescent model.**
(A) Log–log plot showing the relationship between the number of cells harboring a particular L1 and the total count of L1s detected in that many cells.
(B) The posterior population size N versus μ.
(C) The population mutation rate θ versus μ.
(D) Posterior population size N is unchanged across acceptance windows.
(E) The estimated θ is acceptance window-invariant.
(F) Mutation dating using the Kingman's coalescent. The x-axis shows the fraction of cells/nuclei which harbored the L1Hs in the simulation, and the y-axis shows the estimated number of generations normalized by population size since the most recent common ancestor (MRCA).
(G) Test of whether the gap at intermediate detected frequencies (≈10–100 cells, grey band) can be produced by detection limitations alone.

**Figure S11**

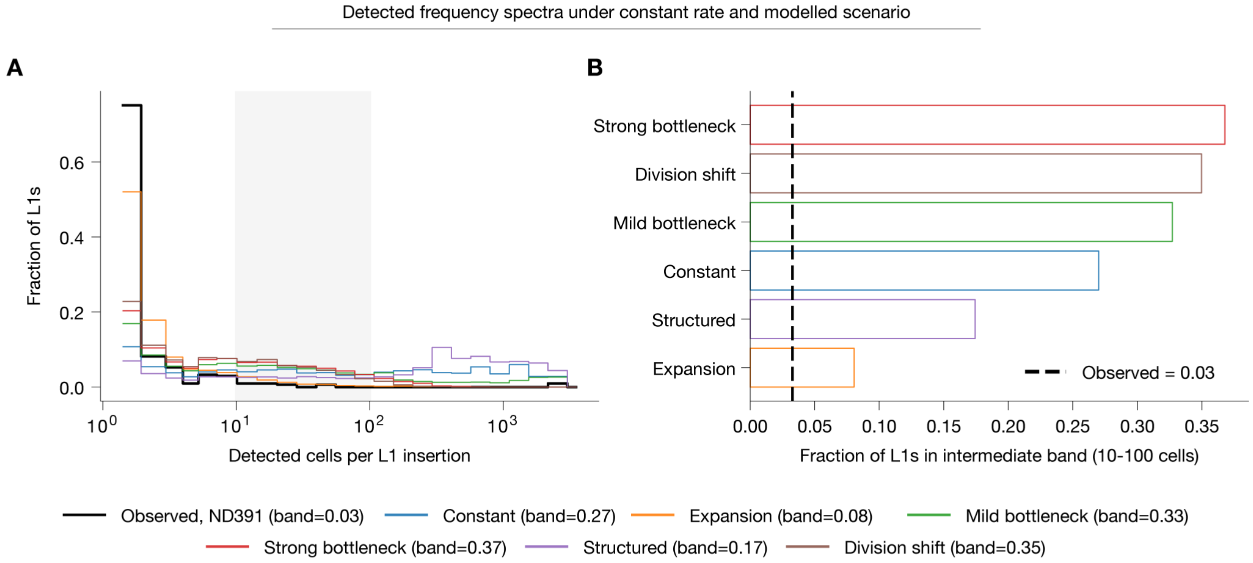


**Figure S11. Coalescent simulations testing whether a constant retrotransposition rate can explain the observed two-wave spectrum.**
(A) Detected frequency spectra (fraction of insertions versus detected cells per L1) for the observed ND391 data (black) and each modelled scenario. The intermediate band (10–100 cells) is shaded.
(B) Fraction of insertions falling in the intermediate band for each scenario, ranked, against the observed value (dashed line, 0.03).

**Figure S12**

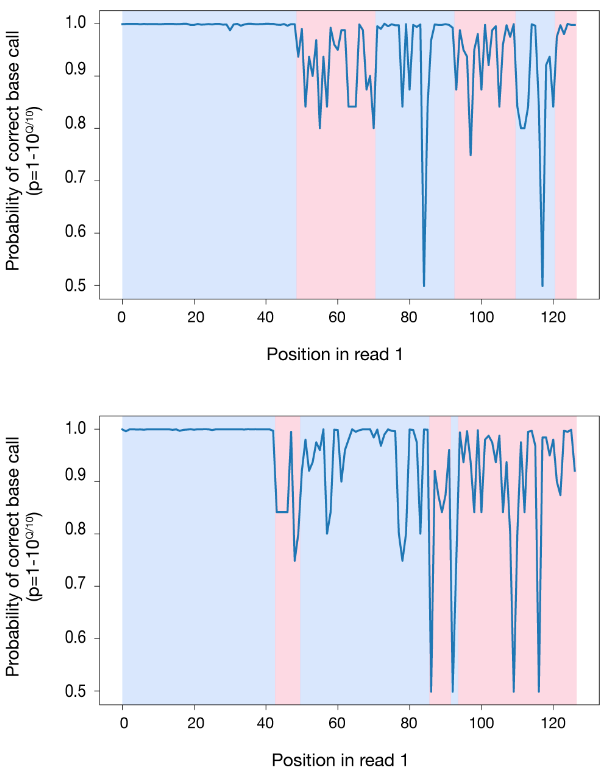


**Figure S12. Representative traces of base calling qualities for LUSTRE read 1 containing very long polyA tracts from somatic L1s.** The colored (blue/red) background shows the estimated break-points found by the implemented algorithm to estimate the length of long polyA tracts. Red indicates a detected decrease break-point in the quality score across the interval. Blue indicates a detected increase break-point in quality score across the interval.
