## Supplementary material for "Dynamics of somatic L1 retrotransposition in the human brain": Note S1

**Note S1**
for PCR and WGS validation of L1s detected by LUSTRE


| | |
|---|---|
| **Selection of L1s** | p. 2 |
| **Validation of L1 insertion sites by targeted long-range PCR** | p. 3 |
| **Categorization of L1s for post-PCR validation rate estimations** | p. 29 |
| **Validation of L1 insertion sites by bulk whole-genome sequencing** | p. 32 |

### Selection of L1s

A subset of discovered somatic L1 were subjected to PCR validation. 22, 22 and 19 L1s were selected for ND507, ND480 and ND391 respectively. The selection procedure was done semi-randomly where all the L1s detected across multiple nuclei were selected first. Then, L1s detected in one nuclei were randomly picked for validation to reach approximately 20 L1s per donor. In addition, 4 polymorphic germline insertions were selected for each donor. Below is the summary of putative somatic L1s subjected to PCR validation.

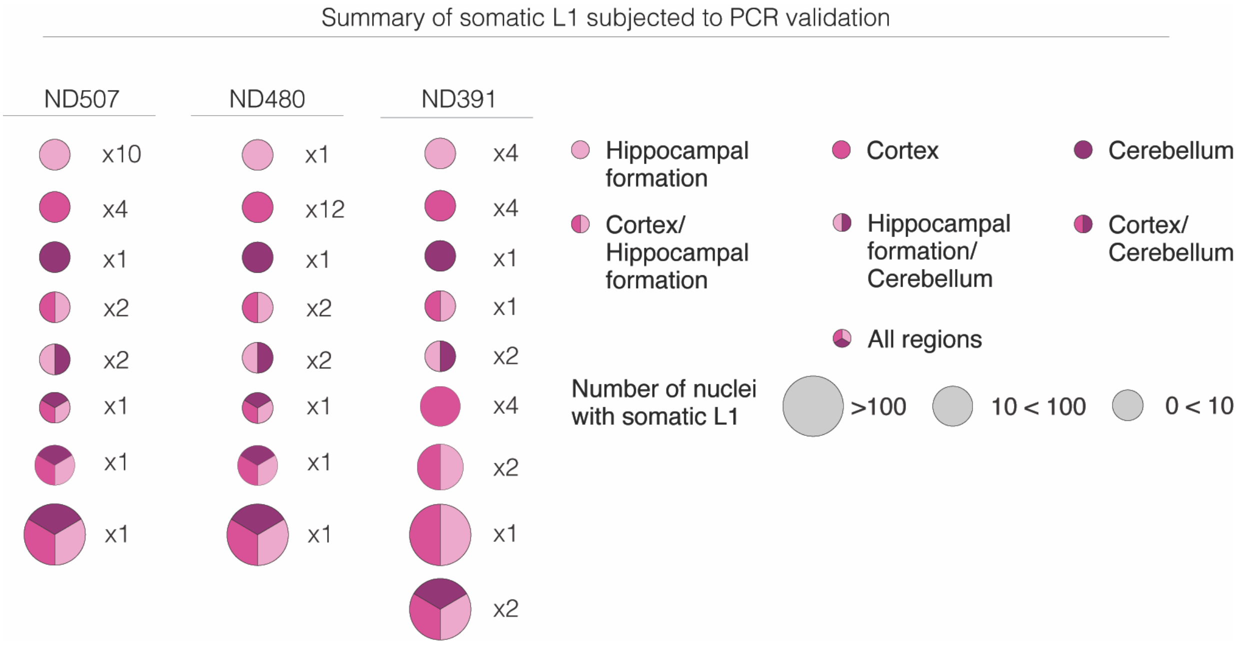

**Validation of L1 insertion sites by targeted long-range PCR**

Here, the number of tagmentation events indicate the number of unique read 1 start sites (excluding soft clipping reads) with a read count threshold of 10 reads per position. Each position is only counted once per insertions across all cells. Therefore, highly abundant insertions (e.g. polymorphic germline) will have considerably fewer tagmentation events than detected neurons. Moreover, the IGV screenshot may show more unique read 1 start sites than the statistic indicates, since a read count threshold has been applied to this value.

**Donor ND507**

<u>**L1-1366**</u> (Polymorphic Germline)

| Location | Strand | Number of tagmentation events | Number of single neurons | Number of sample plates |
|---|---|---|---|---|
| chr4:163751901 | + | 338 | 2469 | 14 |

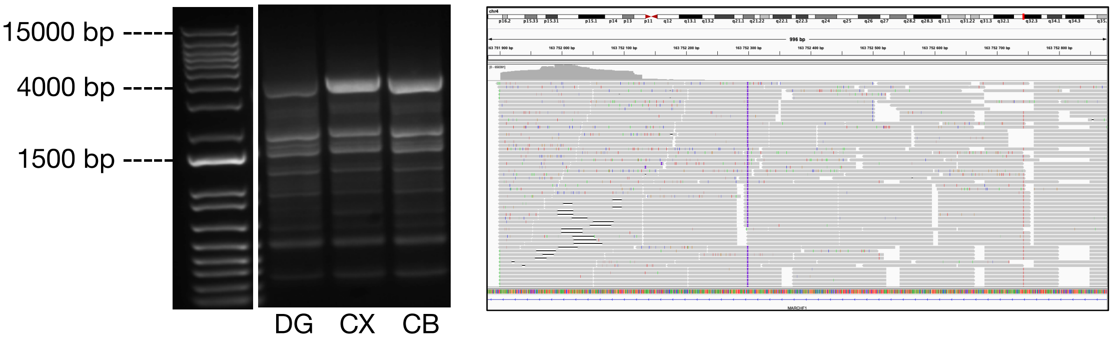


<u>**L1-1949**</u> (Polymorphic Germline)

| Location | Strand | Number of tagmentation events | Number of single neurons | Number of sample plates |
|---|---|---|---|---|
| chr9:8727676745 | - | 457 | 2863 | 14 |

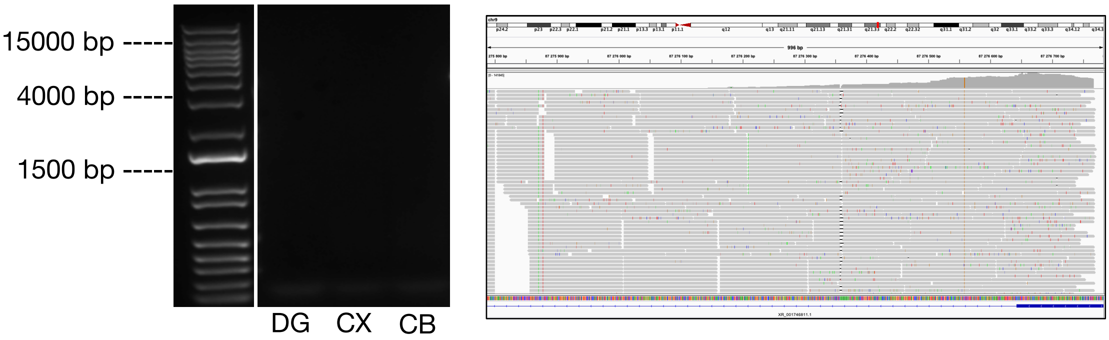

**L1-2285** (Somatic)

| Location | Strand | Number of tagmentation events | Number of single neurons | Number of sample plates |
|---|---|---|---|---|
| chr12:14150445 | + | 154 | 830 | 6 |

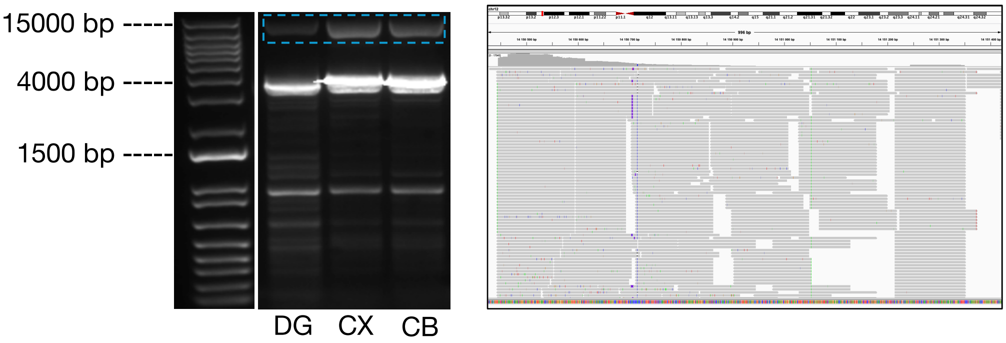


**L1-365** (Polymorphic Germline)

| Location | Strand | Number of tagmentation events | Number of single neurons | Number of sample plates |
|---|---|---|---|---|
| chr12:12780760 | - | 492 | 3955 | 14 |

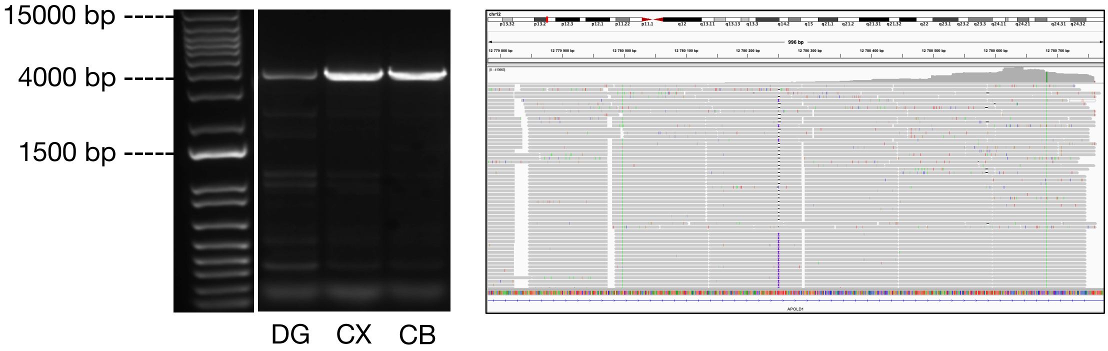


**L1-5439** (Somatic)

| Location | Strand | Number of tagmentation events | Number of single neurons | Number of sample plates |
|---|---|---|---|---|
| chr19:19987166 | + | 7 | 12 | 4 |

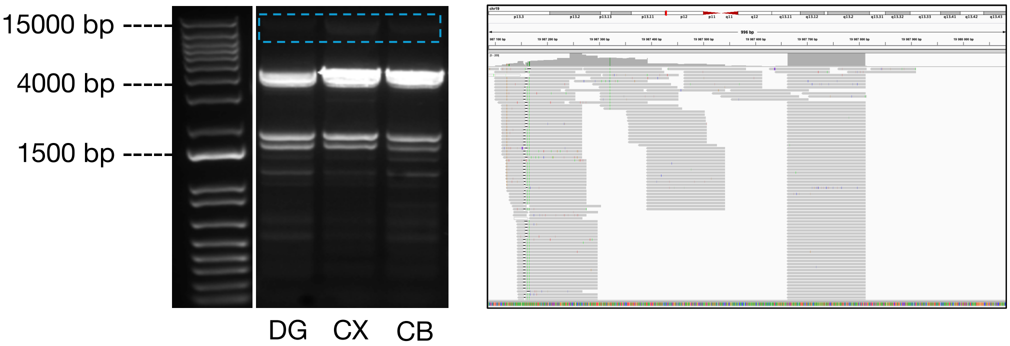

**L1-68** (Somatic)

| Location | Strand | Number of tagmentation events | Number of single neurons | Number of sample plates |
|---|---|---|---|---|
| chr1:12247502 | + | 3 | 7 | 5 |

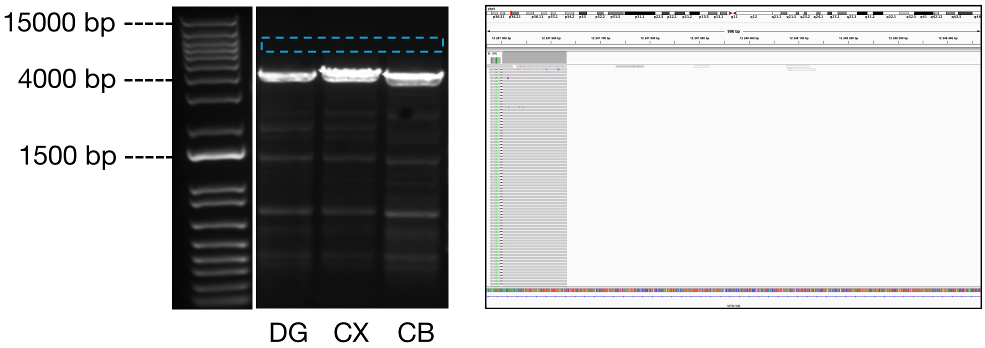


**L1-23** (Somatic)

| Location | Strand | Number of tagmentation events | Number of single neurons | Number of sample plates |
|---|---|---|---|---|
| chr1:4706506 | - | 2 | 2 | 2 |

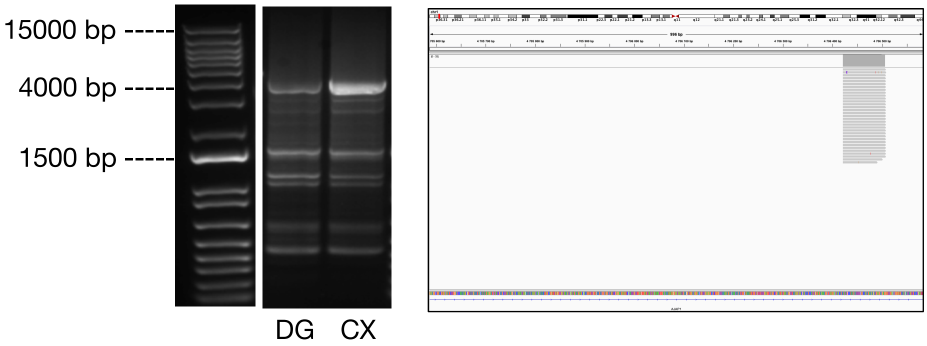


**L1-3139** (Somatic)

| Location | Strand | Number of tagmentation events | Number of single neurons | Number of sample plates |
|---|---|---|---|---|
| chr13:81793576 | - | 2 | 2 | 2 |

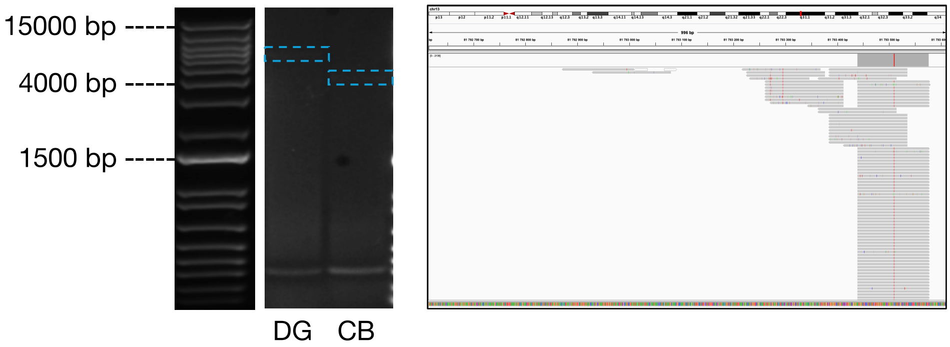

**L1-9065** (Somatic)

| Location | Strand | Number of tagmentation events | Number of single neurons | Number of sample plates |
|---|---|---|---|---|
| chr4:142141058 | - | 3 | 2 | 1 |

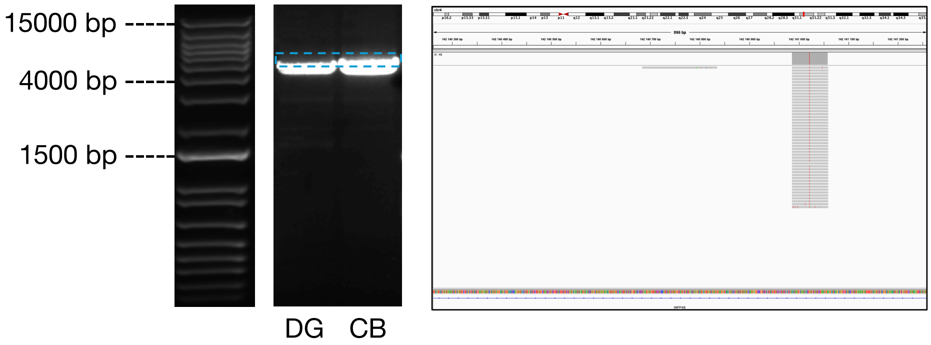


**L1-9104** (Somatic)

| Location | Strand | Number of tagmentation events | Number of single neurons | Number of sample plates |
|---|---|---|---|---|
| chr4:151041509 | - | 2 | 2 | 2 |

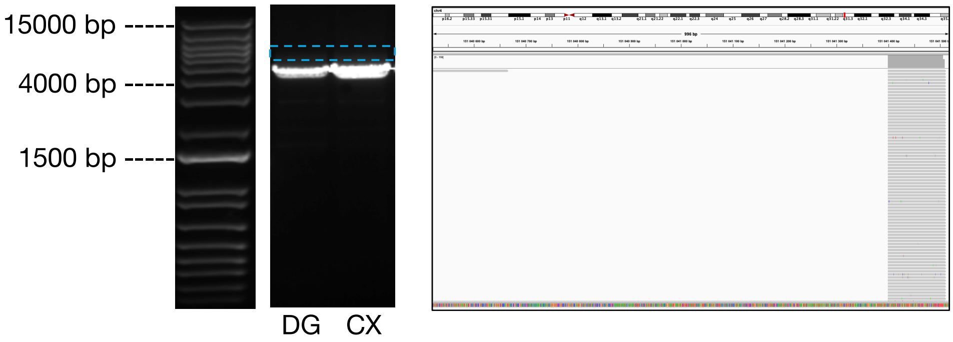


**L1-10422** (Somatic)

| Location | Strand | Number of tagmentation events | Number of single neurons | Number of sample plates |
|---|---|---|---|---|
| chr6:71399182 | + | 2 | 1 | 1 |

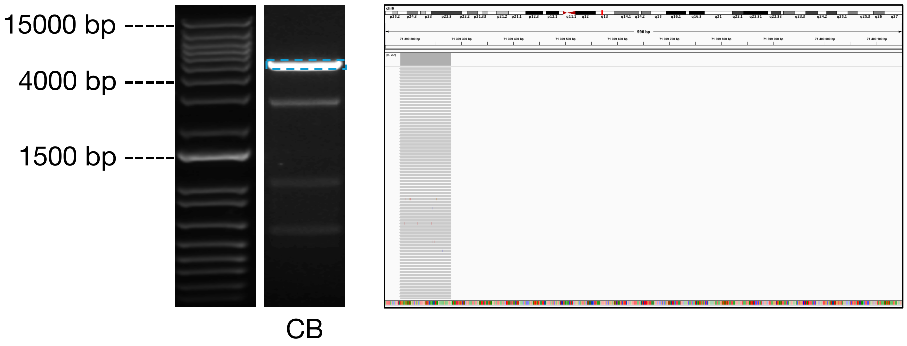

**L1-12405** (Somatic)

| Location | Strand | Number of tagmentation events | Number of single neurons | Number of sample plates |
|---|---|---|---|---|
| chr9:25584446 | + | 2 | 3 | 2 |

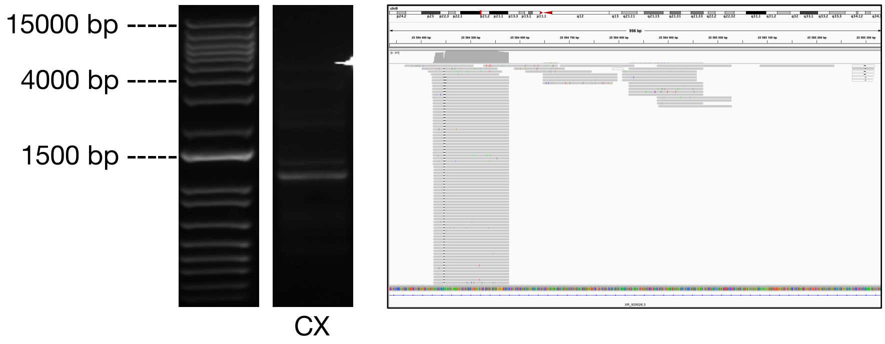


**L1-1633** (Somatic)

| Location | Strand | Number of tagmentation events | Number of single neurons | Number of sample plates |
|---|---|---|---|---|
| chr11:10550060 | - | 1 | 1 | 1 |

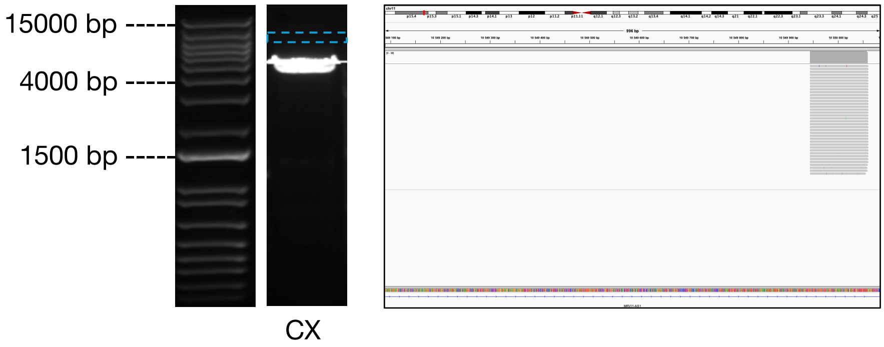


**L1-4348** (Somatic)

| Location | Strand | Number of tagmentation events | Number of single neurons | Number of sample plates |
|---|---|---|---|---|
| chr16:62292243 | - | 1 | 1 | 1 |

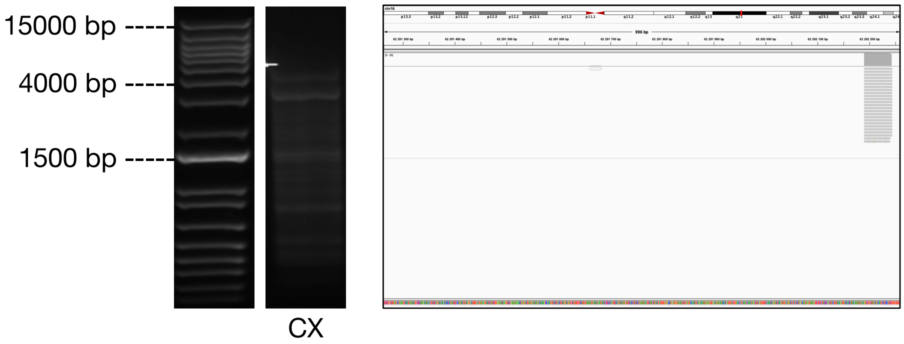

**L1-6489** (Somatic)

| Location | Strand | Number of tagmentation events | Number of single neurons | Number of sample plates |
|---|---|---|---|---|
| chr2:180110586 | + | 5 | 8 | 4 |

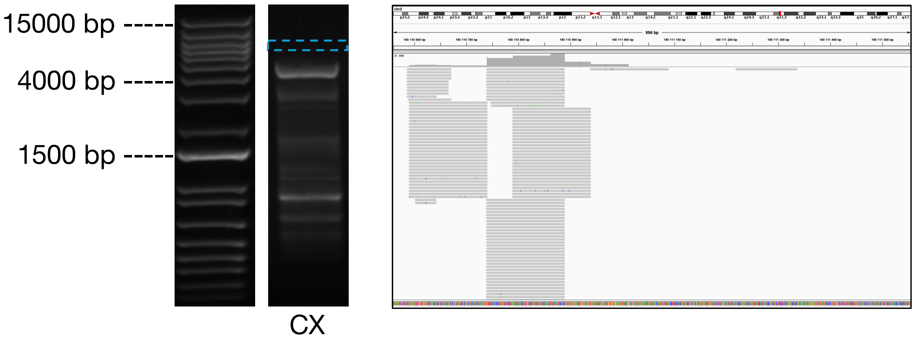


**L1-10069** (Somatic)

| Location | Strand | Number of tagmentation events | Number of single neurons | Number of sample plates |
|---|---|---|---|---|
| chr5:177123696 | + | 1 | 1 | 1 |

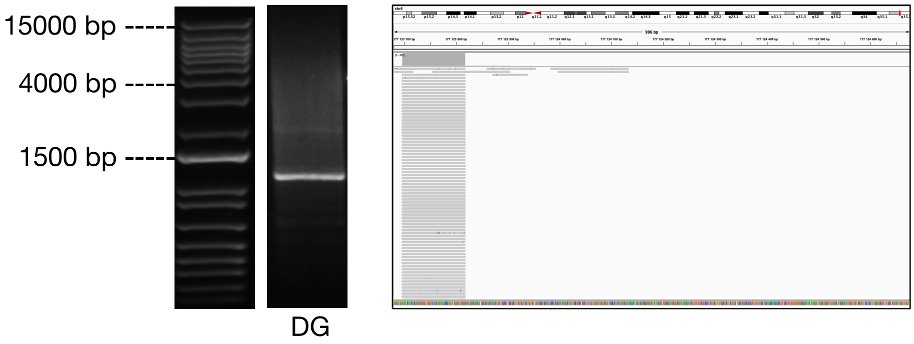


**L1-10580** (Somatic)

| Location | Strand | Number of tagmentation events | Number of single neurons | Number of sample plates |
|---|---|---|---|---|
| chr6:106826781 | + | 1 | 1 | 1 |

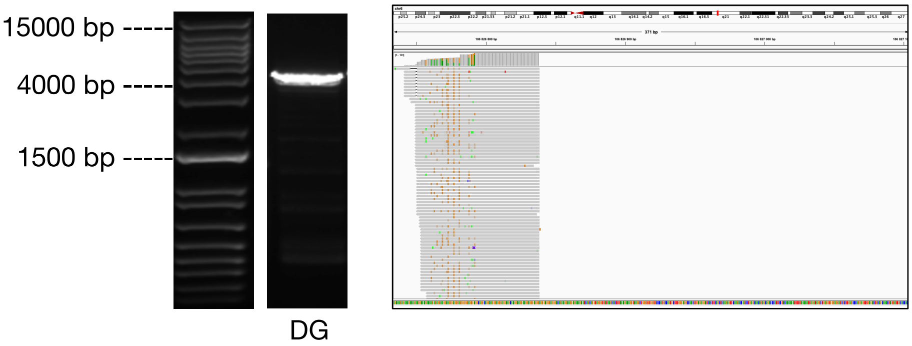

**L1-1153** (Somatic)

| Location | Strand | Number of tagmentation events | Number of single neurons | Number of sample plates |
|---|---|---|---|---|
| chr10:30105372 | - | 1 | 1 | 1 |

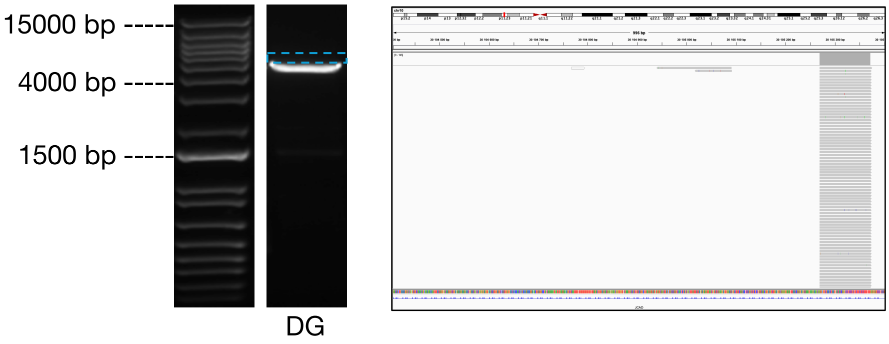


**L1-13029** (Somatic)

| Location | Strand | Number of tagmentation events | Number of single neurons | Number of sample plates |
|---|---|---|---|---|
| chrX:41262073 | - | 1 | 1 | 1 |

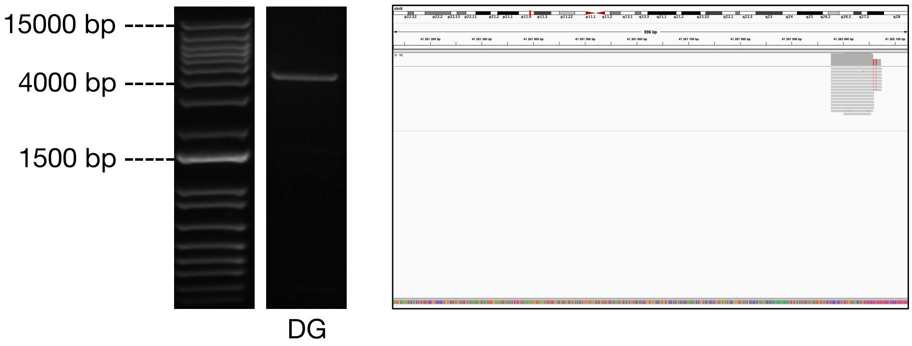


**L1-178** (Somatic)

| Location | Strand | Number of tagmentation events | Number of single neurons | Number of sample plates |
|---|---|---|---|---|
| chr1:35228618 | - | 2 | 2 | 2 |

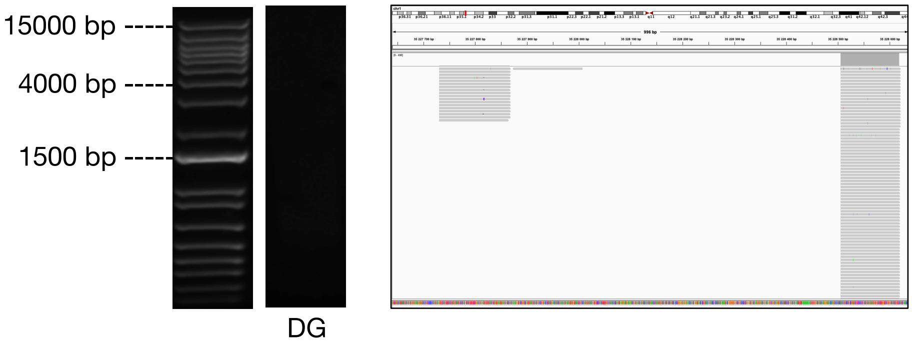

**L1-1972** (Polymorphic Germline)

| Location | Strand | Number of tagmentation events | Number of single neurons | Number of sample plates |
|---|---|---|---|---|
| chr9:115321601 | - | 481 | 3040 | 14 |

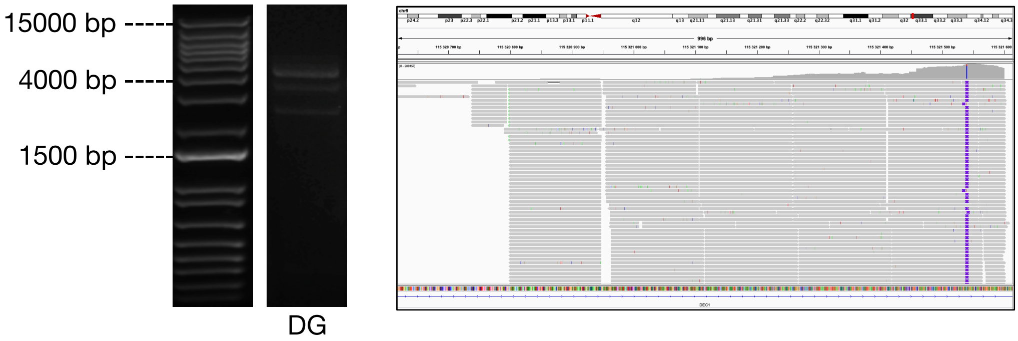


**L1-2300** (Somatic)

| Location | Strand | Number of tagmentation events | Number of single neurons | Number of sample plates |
|---|---|---|---|---|
| chr12:16405173 | + | 2 | 1 | 1 |

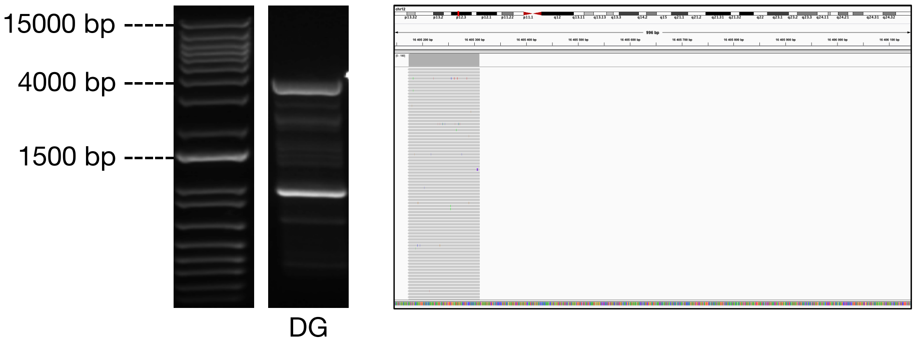


**L1-8148** (Somatic)

| Location | Strand | Number of tagmentation events | Number of single neurons | Number of sample plates |
|---|---|---|---|---|
| chr3:135376549 | + | 1 | 1 | 1 |

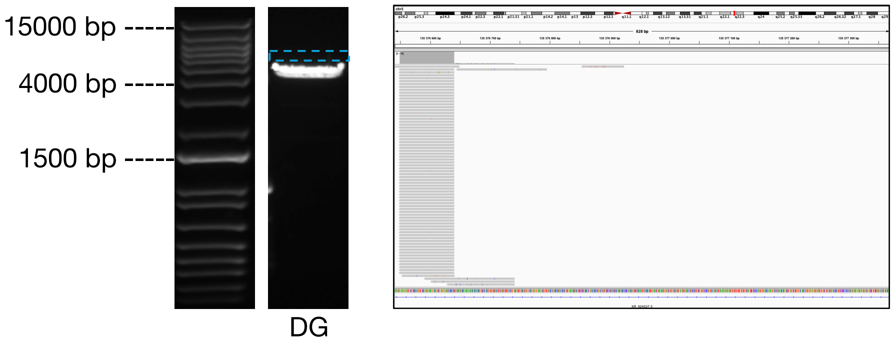

**L1-8435** (Somatic)

| Location | Strand | Number of tagmentation events | Number of single neurons | Number of sample plates |
|---|---|---|---|---|
| chr3:197206747 | + | 1 | 1 | 1 |

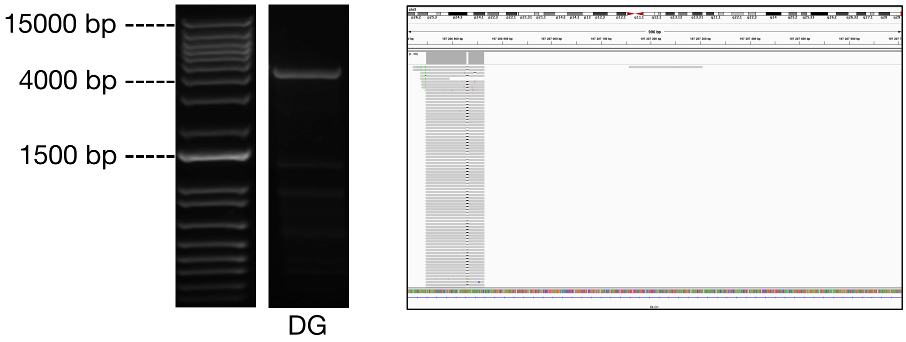


**L1-9692** (Somatic)

| Location | Strand | Number of tagmentation events | Number of single neurons | Number of sample plates |
|---|---|---|---|---|
| chr5:97424445 | - | 2 | 2 | 2 |

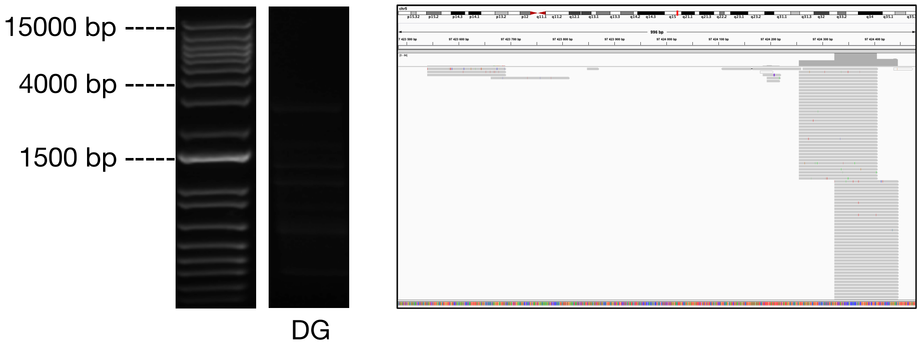


**L1-9734** (Somatic)

| Location | Strand | Number of tagmentation events | Number of single neurons | Number of sample plates |
|---|---|---|---|---|
| chr5:105697264 | + | 1 | 1 | 1 |

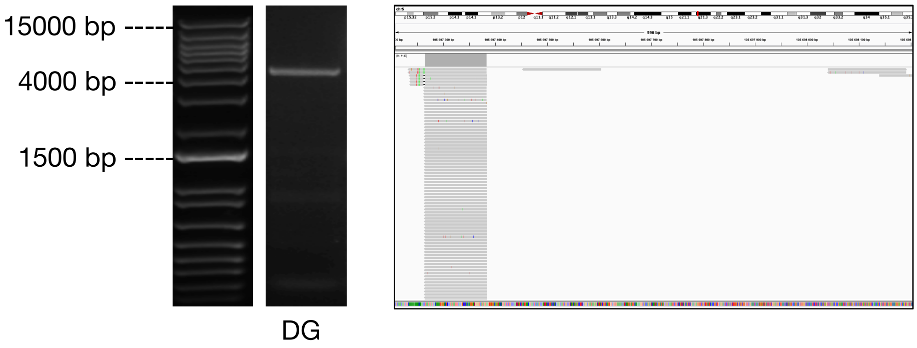

**Donor ND480**

**L1-1111** (Polymorphic germline)

| Location | Strand | Number of tagmentation events | Number of single neurons | Number of sample plates |
|---|---|---|---|---|
| chr3:5986713 | + | 581 | 3195 | 11 |

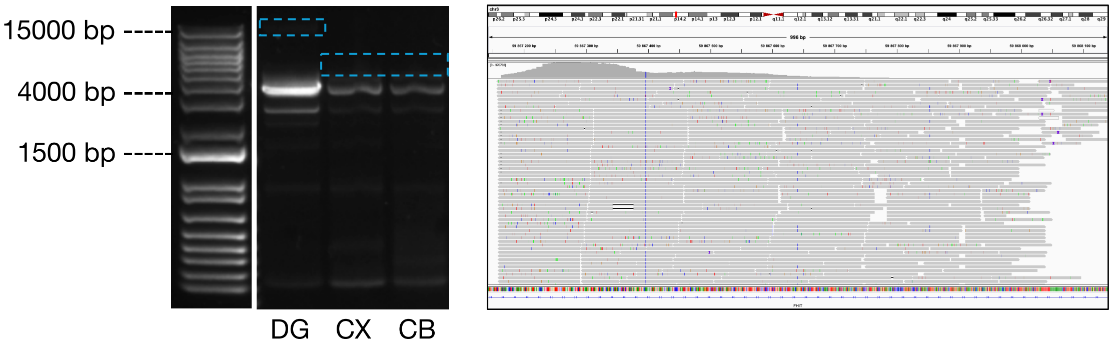


**L1-1800** (Polymorphic germline)

| Location | Strand | Number of tagmentation events | Number of single neurons | Number of sample plates |
|---|---|---|---|---|
| chr8:5971760 | - | 641 | 3615 | 11 |

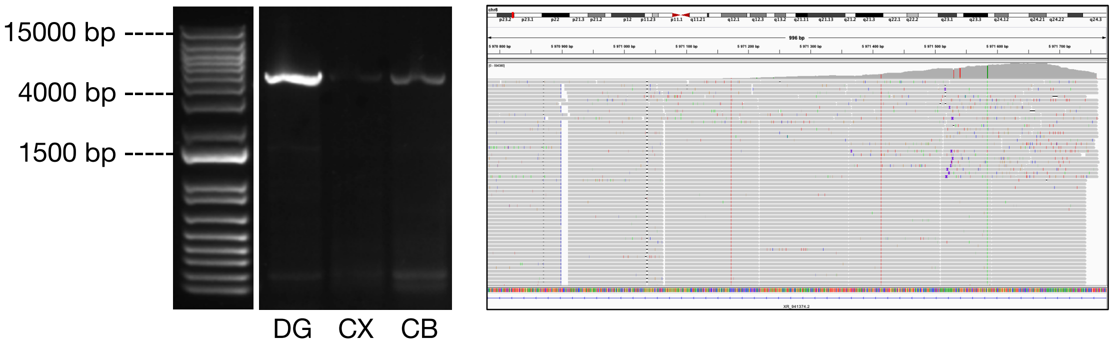


**L1-1938** (Polymorphic germline)

| Location | Strand | Number of tagmentation events | Number of single neurons | Number of sample plates |
|---|---|---|---|---|
| chr9:72290005 | - | 494 | 3122 | 11 |

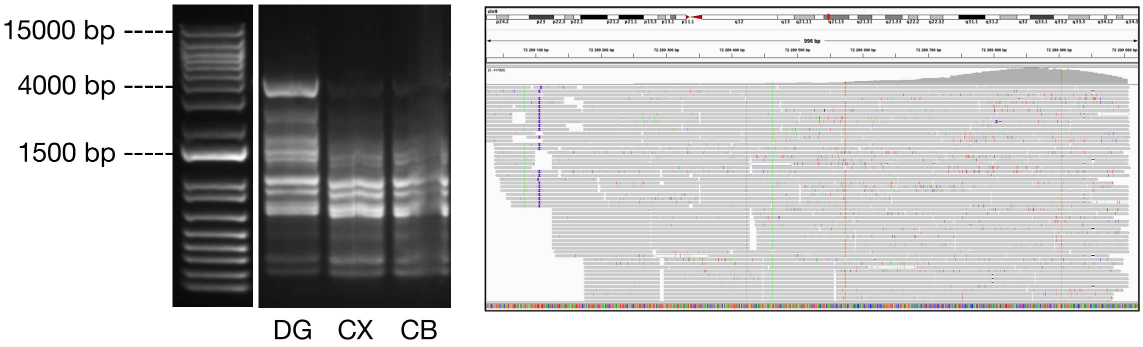

**L1-4062** (Somatic)

| Location | Strand | Number of tagmentation events | Number of single neurons | Number of sample plates |
|---|---|---|---|---|
| chr15:92479991 | - | 18 | 23 | 6 |

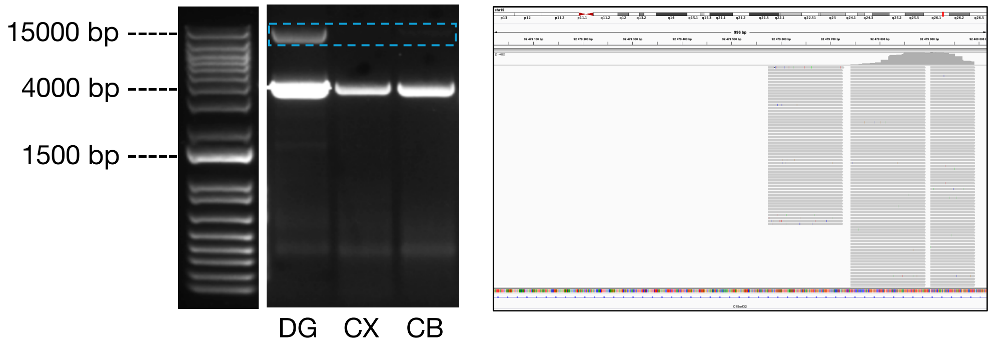


**L1-6480** (Somatic)

| Location | Strand | Number of tagmentation events | Number of single neurons | Number of sample plates |
|---|---|---|---|---|
| chr2:177932099 | + | 59 | 165 | 6 |

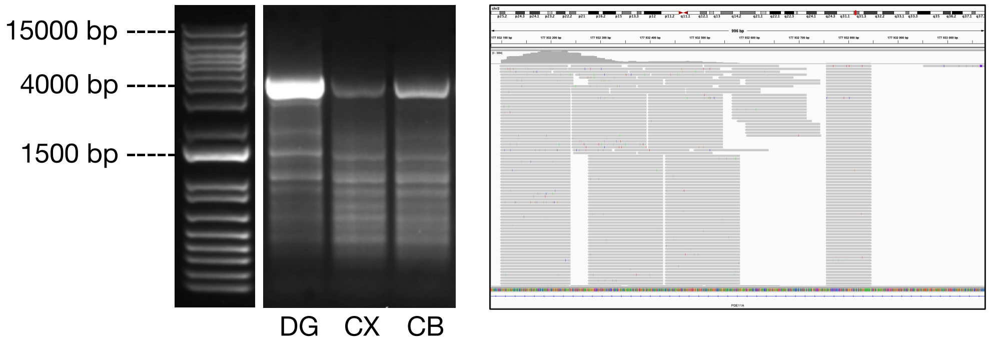


**L1-6528** (Somatic)

| Location | Strand | Number of tagmentation events | Number of single neurons | Number of sample plates |
|---|---|---|---|---|
| chr2:189228420 | + | 7 | 3 | 3 |

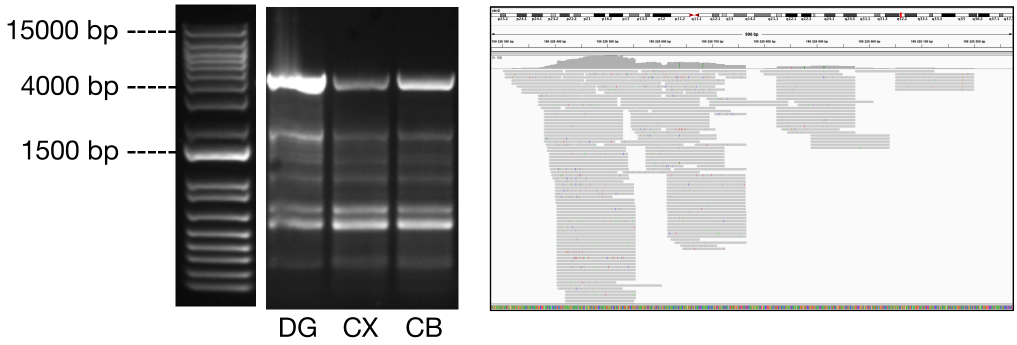

**L1-959** (Polymorphic germline)

| Location | Strand | Number of tagmentation events | Number of single neurons | Number of sample plates |
|---|---|---|---|---|
| chr2:198915174 | + | 563 | 3142 | 11 |

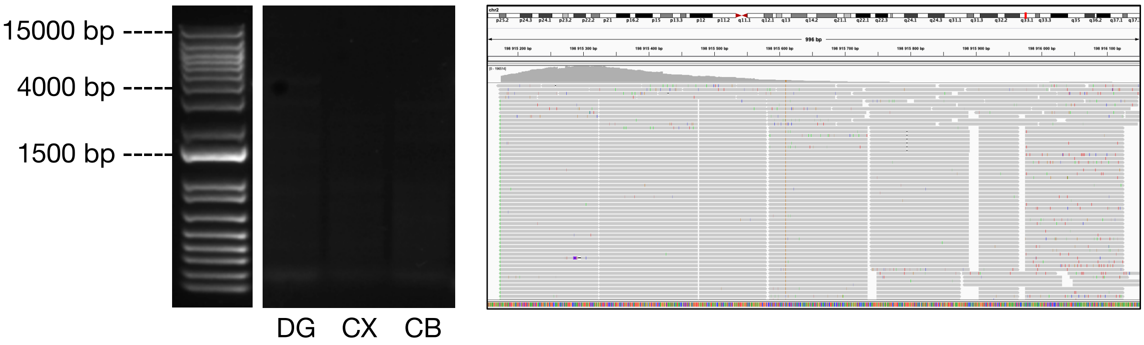


**L1-10904** (Somatic)

| Location | Strand | Number of tagmentation events | Number of single neurons | Number of sample plates |
|---|---|---|---|---|
| chr7:5209034 | - | 3 | 4 | 2 |

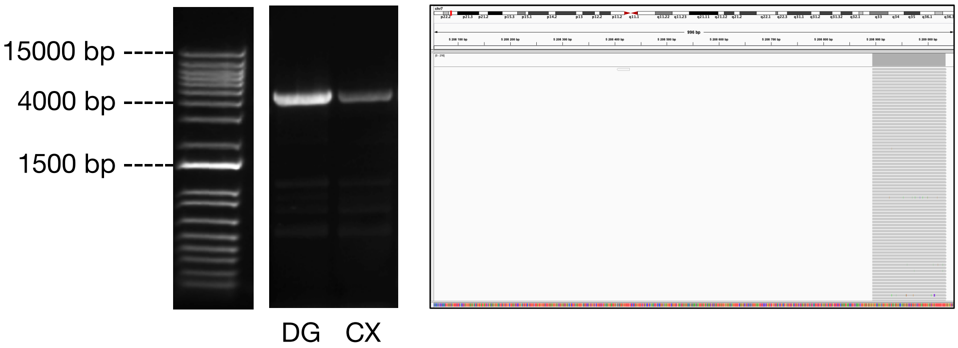


**L1-723** (Somatic)

| Location | Strand | Number of tagmentation events | Number of single neurons | Number of sample plates |
|---|---|---|---|---|
| chr1:178223717 | - | 3 | 3 | 3 |

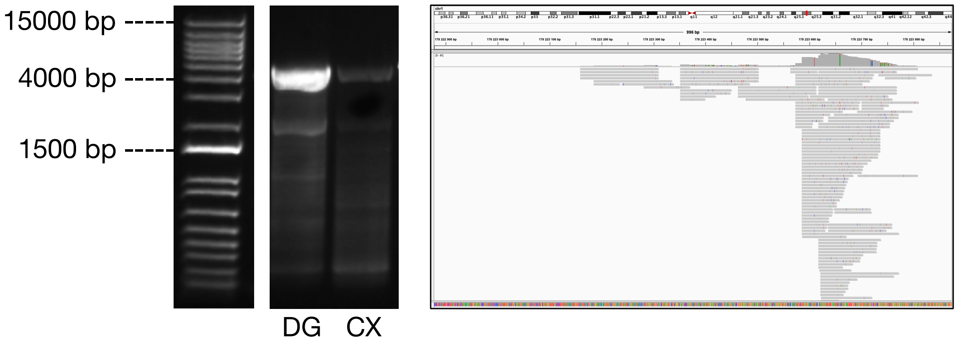

**L1-6550** (Somatic)

| Location | Strand | Number of tagmentation events | Number of single neurons | Number of sample plates |
|---|---|---|---|---|
| chr2:194354793 | + | 2 | 3 | 1 |

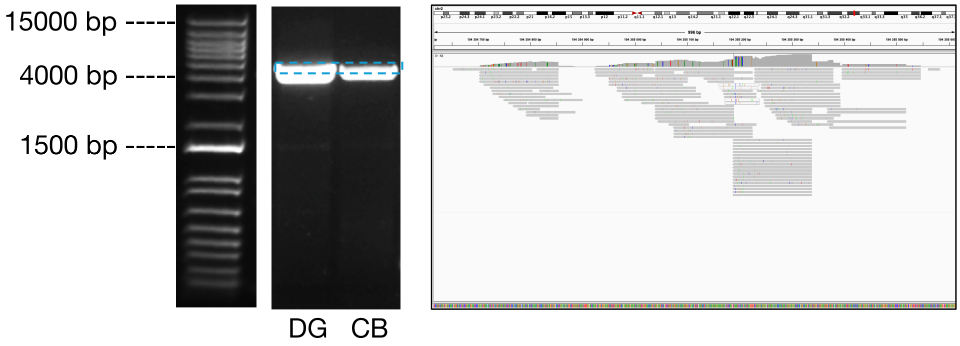


**L1-6782** (Somatic)

| Location | Strand | Number of tagmentation events | Number of single neurons | Number of sample plates |
|---|---|---|---|---|
| chr2:240309527 | - | 1 | 2 | 2 |

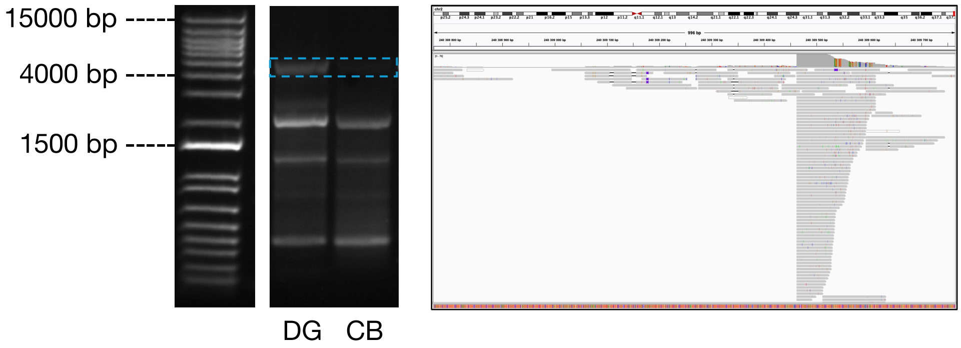


**L1-10286** (Somatic)

| Location | Strand | Number of tagmentation events | Number of single neurons | Number of sample plates |
|---|---|---|---|---|
| chr6:40069543 | + | 9 | 14 | 4 |

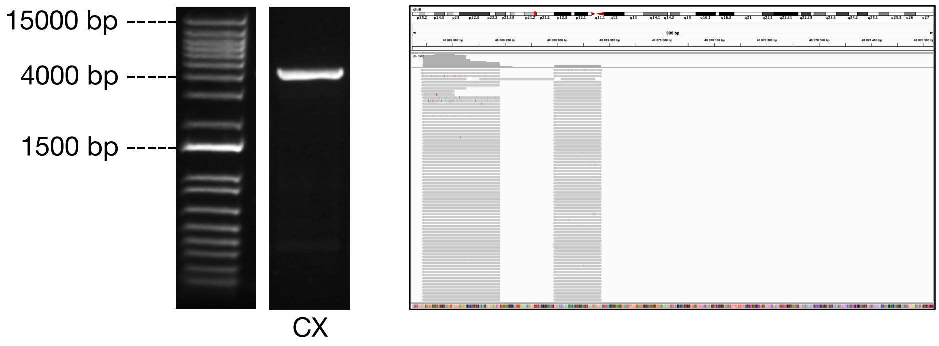

**<u>L1-11813</u>** (Somatic)

| Location | Strand | Number of tagmentation events | Number of single neurons | Number of sample plates |
|---|---|---|---|---|
| chr8:49993522 | - | 2 | 2 | 2 |

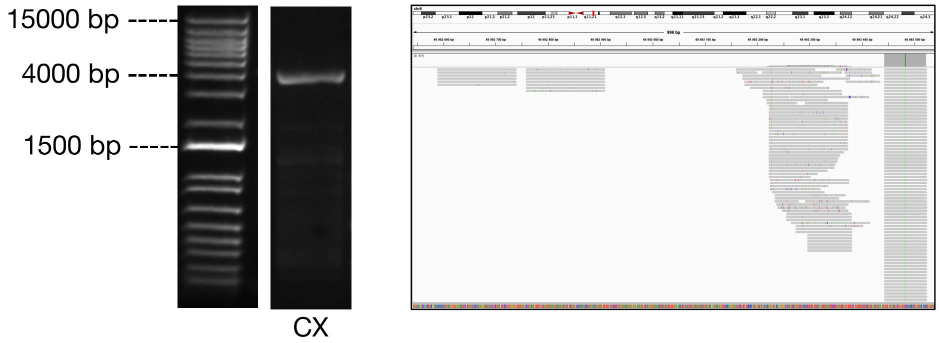


**<u>L1-134</u>** (Somatic)

| Location | Strand | Number of tagmentation events | Number of single neurons | Number of sample plates |
|---|---|---|---|---|
| chr1:23630666 | + | 2 | 1 | 1 |

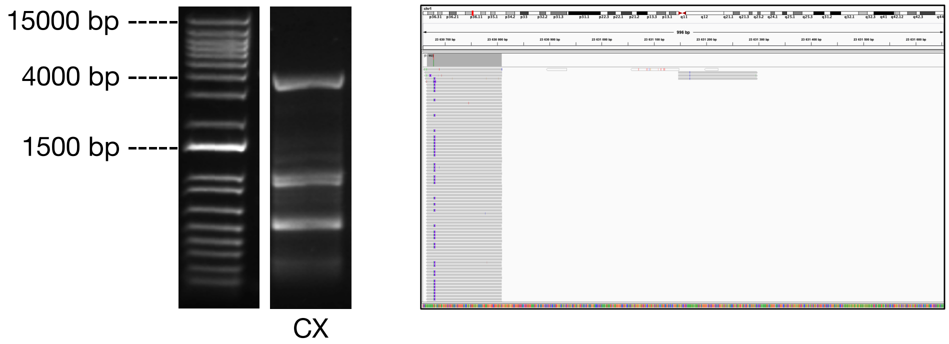


**<u>L1-2231</u>** (Somatic)

| Location | Strand | Number of tagmentation events | Number of single neurons | Number of sample plates |
|---|---|---|---|---|
| chr12:4325764 | + | 1 | 1 | 1 |

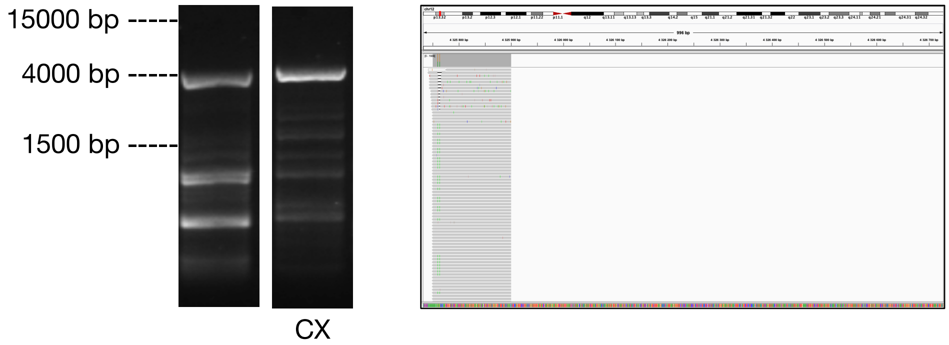

**L1-3110** (Somatic)

| Location | Strand | Number of tagmentation events | Number of single neurons | Number of sample plates |
|---|---|---|---|---|
| chr13:74566533 | - | 1 | 1 | 1 |

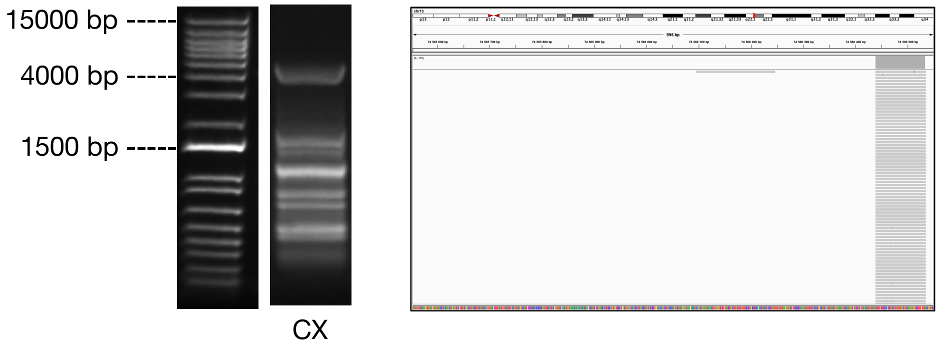


**L1-4164** (Somatic)

| Location | Strand | Number of tagmentation events | Number of single neurons | Number of sample plates |
|---|---|---|---|---|
| chr16:14673010 | - | 2 | 8 | 2 |

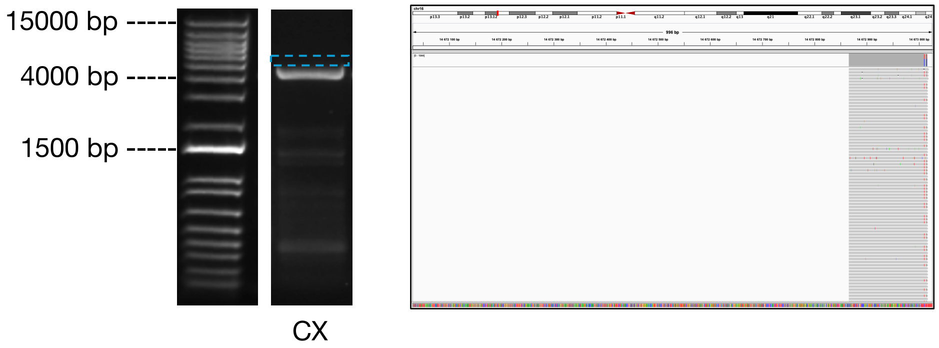


**L1-5568** (Somatic)

| Location | Strand | Number of tagmentation events | Number of single neurons | Number of sample plates |
|---|---|---|---|---|
| chr19:46154402 | - | 1 | 1 | 1 |

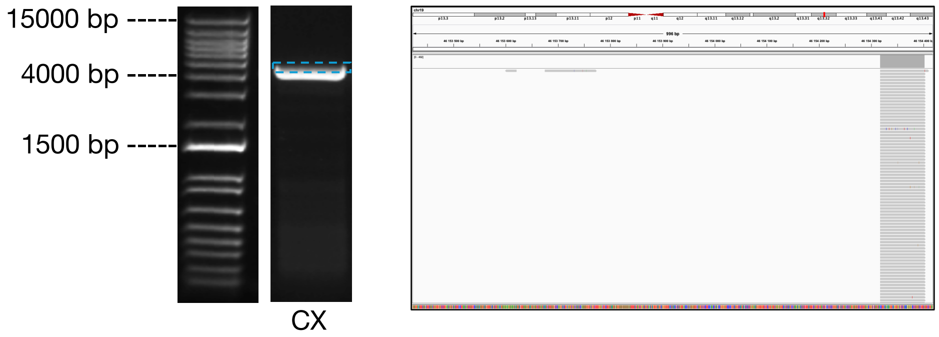

**L1-5972** (Somatic)

| Location | Strand | Number of tagmentation events | Number of single neurons | Number of sample plates |
|---|---|---|---|---|
| chr2:75303494 | + | 1 | 1 | 1 |

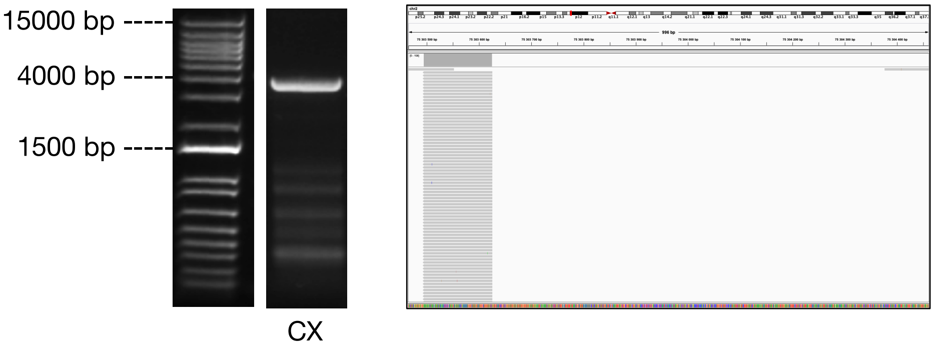


**L1-606** (Somatic)

| Location | Strand | Number of tagmentation events | Number of single neurons | Number of sample plates |
|---|---|---|---|---|
| chr1:155644953 | + | 1 | 1 | 1 |

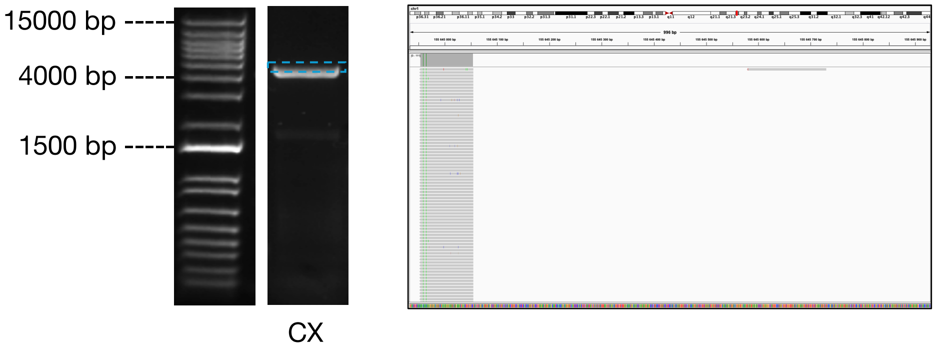


**L1-6156** (Somatic)

| Location | Strand | Number of tagmentation events | Number of single neurons | Number of sample plates |
|---|---|---|---|---|
| chr2:116517838 | + | 1 | 2 | 1 |

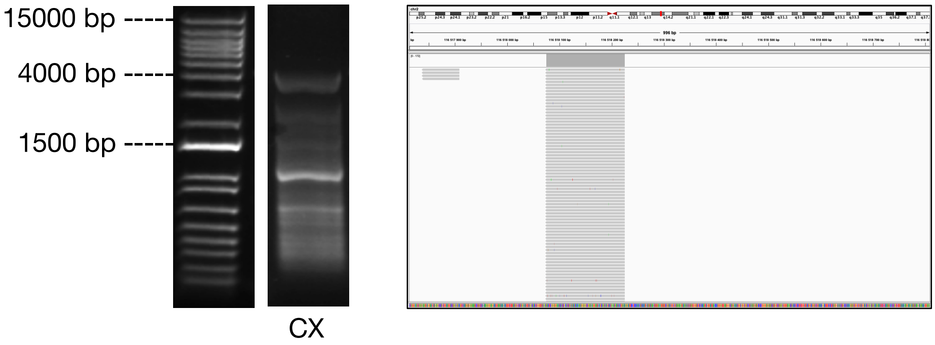

**<u>L1-6198</u>** (Somatic)

| Location | Strand | Number of tagmentation events | Number of single neurons | Number of sample plates |
|---|---|---|---|---|
| chr2:122758286 | - | 1 | 1 | 1 |

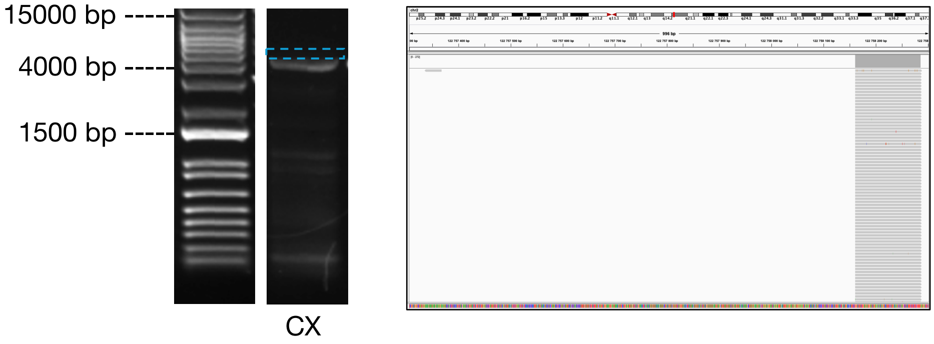


**<u>L1-7027</u>** (Somatic)

| Location | Strand | Number of tagmentation events | Number of single neurons | Number of sample plates |
|---|---|---|---|---|
| chr20:46603183 | + | 1 | 1 | 1 |

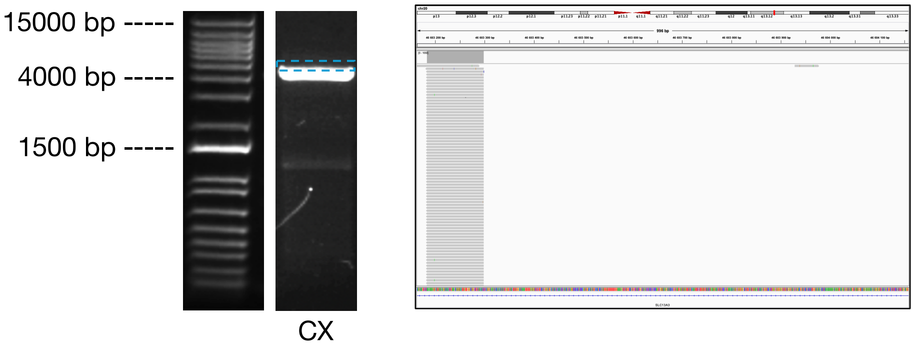


**<u>L1-850</u>** (Somatic)

| Location | Strand | Number of tagmentation events | Number of single neurons | Number of sample plates |
|---|---|---|---|---|
| chr1:208639202 | - | 1 | 3 | 2 |

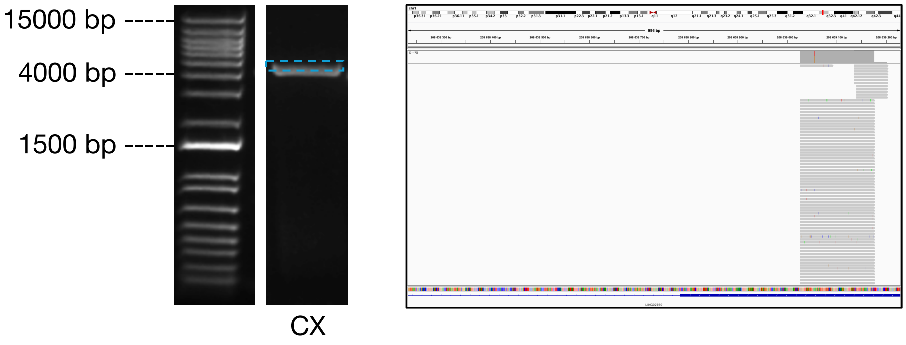

**L1-2734** (Somatic)

| Location | Strand | Number of tagmentation events | Number of single neurons | Number of sample plates |
|---|---|---|---|---|
| chr12:107783208 | - | 2 | 1 | 1 |

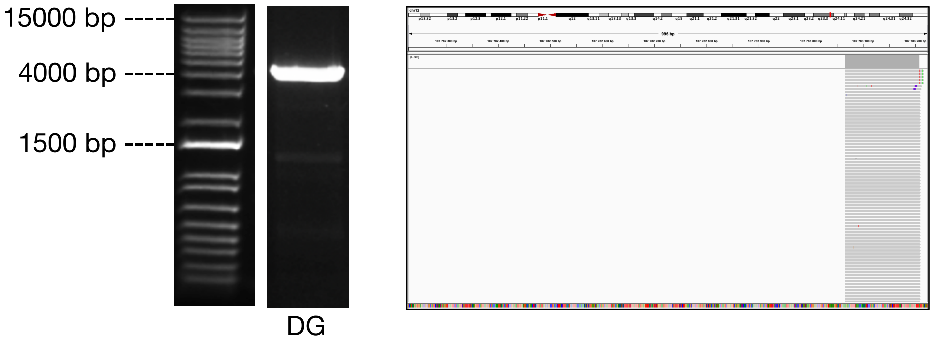


**L1-2829** (Somatic)

| Location | Strand | Number of tagmentation events | Number of single neurons | Number of sample plates |
|---|---|---|---|---|
| chr12:127138676 | - | 2 | 1 | 1 |

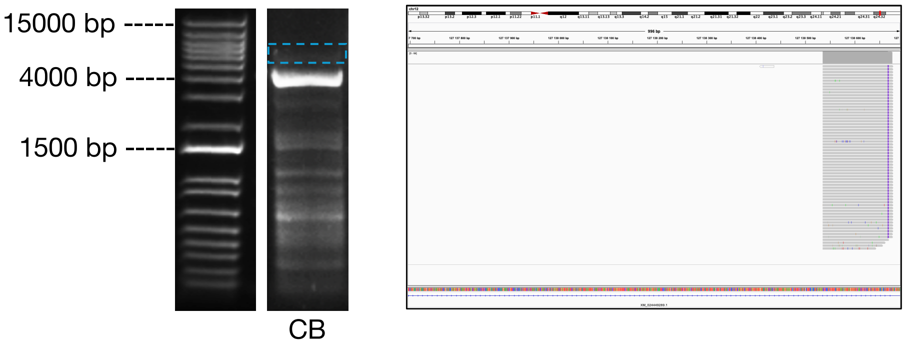

**Donor ND391**

<u>**L1-206**</u> (Polymorphic Germline)

| Location | Strand | Number of tagmentation events | Number of single neurons | Number of sample plates |
|---|---|---|---|---|
| chr15:100390956 | - | 371* | 4685 | 25 |

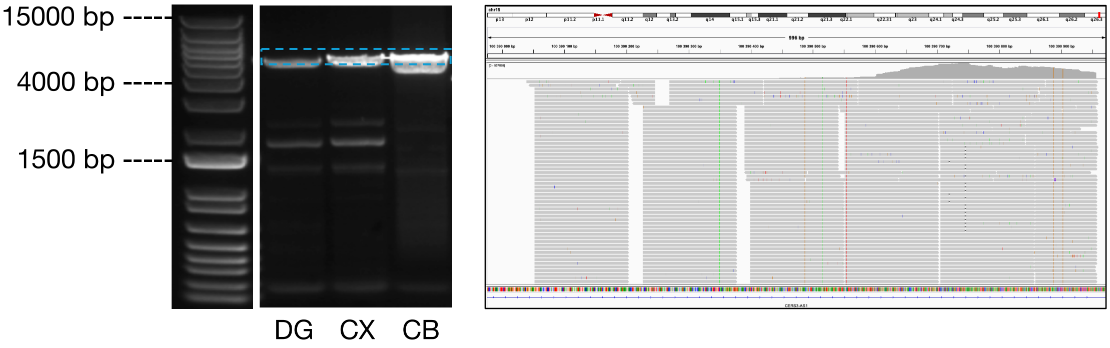


*Lower bound, donor sequenced across multiple experiments.

<u>**L1-133**</u> (Somatic)

| Location | Strand | Number of tagmentation events | Number of single neurons | Number of sample plates |
|---|---|---|---|---|
| chr2:23315448 | + | 188* | 2398 | 25 |

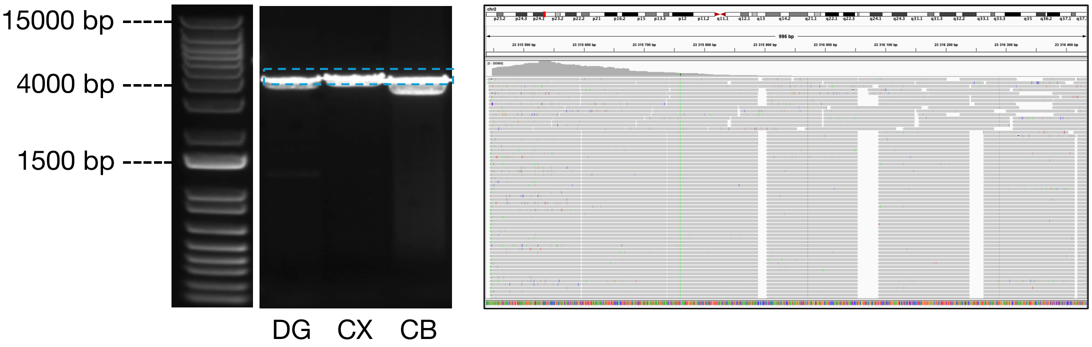


*Lower bound, donor sequenced across multiple experiments.

<u>**L1-503**</u> (Polymorphic Germline)

| Location | Strand | Number of tagmentation events | Number of single neurons | Number of sample plates |
|---|---|---|---|---|
| chr6:62658290 | + | 213* | 5087 | 25 |

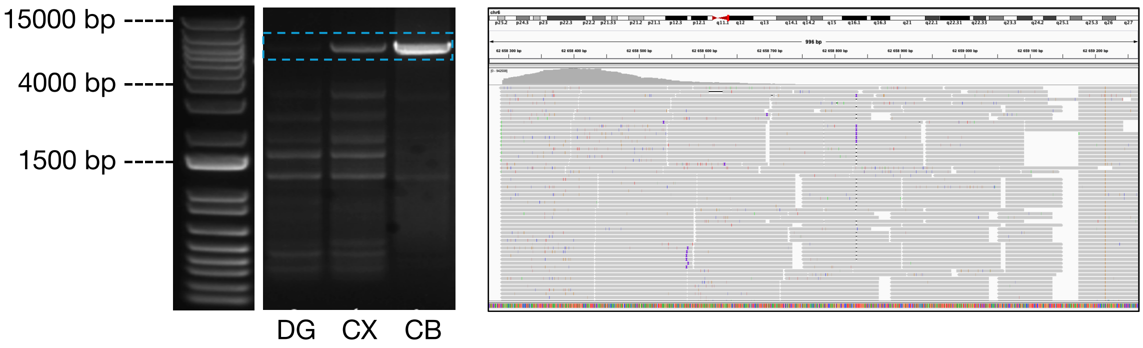


*Lower bound, donor sequenced across multiple experiments.

**L1-551** (Polymorphic Germline)

| Location | Strand | Number of tagmentation events | Number of single neurons | Number of sample plates |
|---|---|---|---|---|
| chr7:63032551 | + | 237* | 6685 | 25 |

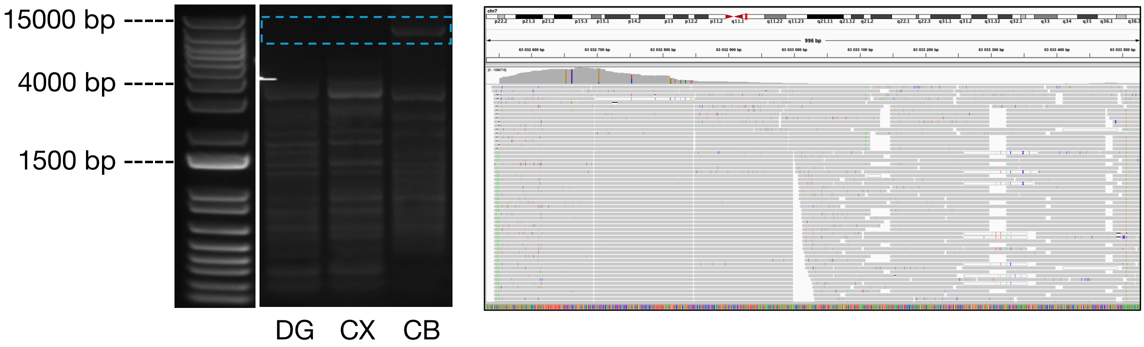


*Lower bound, donor sequenced across multiple experiments.

**L1-654** (Polymorphic Germline)

| Location | Strand | Number of tagmentation events | Number of single neurons | Number of sample plates |
|---|---|---|---|---|
| chrX:111986343 | - | 156* | 3,993 | 25 |

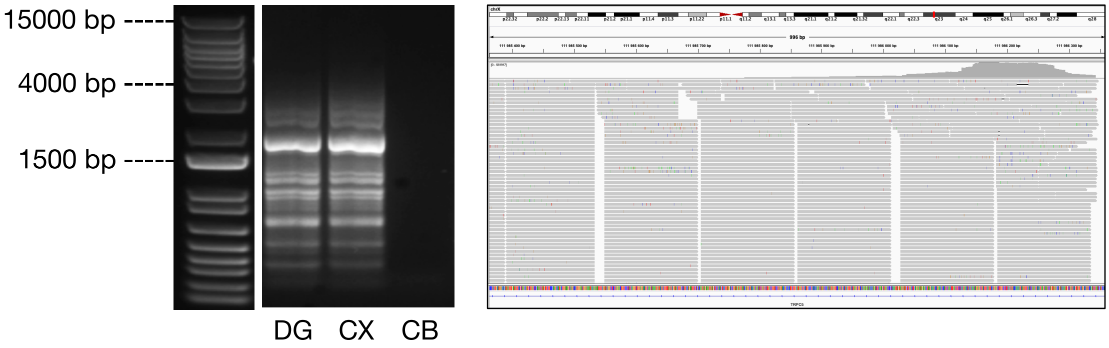


*Lower bound, donor sequenced across multiple experiments.

**L1-248** (Somatic)

| Location | Strand | Number of tagmentation events | Number of single neurons | Number of sample plates |
|---|---|---|---|---|
| chr6:147037216 | + | 195* | 2,385 | 25 |

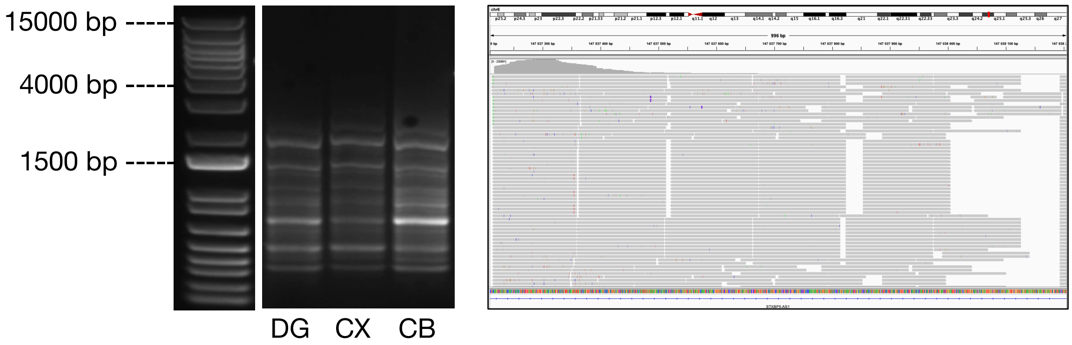


*Lower bound, donor sequenced across multiple experiments.

**L1-197** (Somatic)

| Location | Strand | Number of tagmentation events | Number of single neurons | Number of sample plates |
|---|---|---|---|---|
| chr3:144710822 | - | 4 | 8 | 3 |

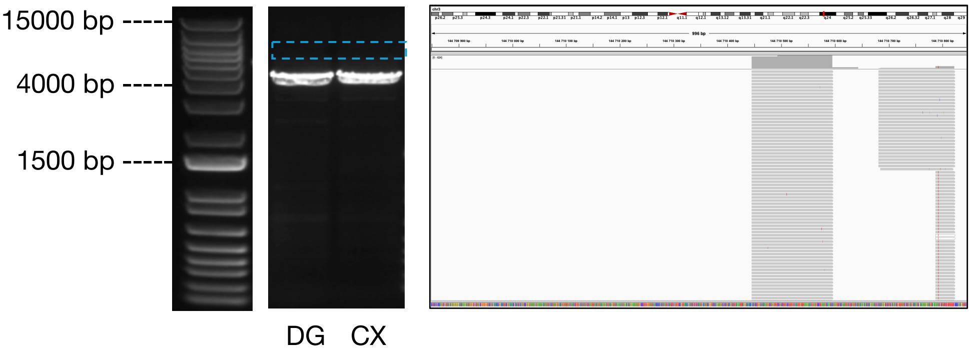


**L1-210** (Somatic)

| Location | Strand | Number of tagmentation events | Number of single neurons | Number of sample plates |
|---|---|---|---|---|
| chr4:173114443 | + | 165* | 2,346 | 23 |

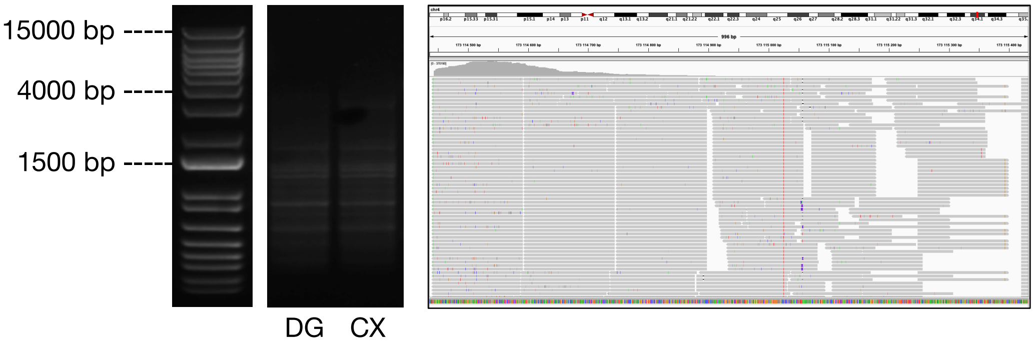

**L1-217** (Somatic)

| Location | Strand | Number of tagmentation events | Number of single neurons | Number of sample plates |
|---|---|---|---|---|
| chr5:41428432 | + | 12* | 52 | 14 |

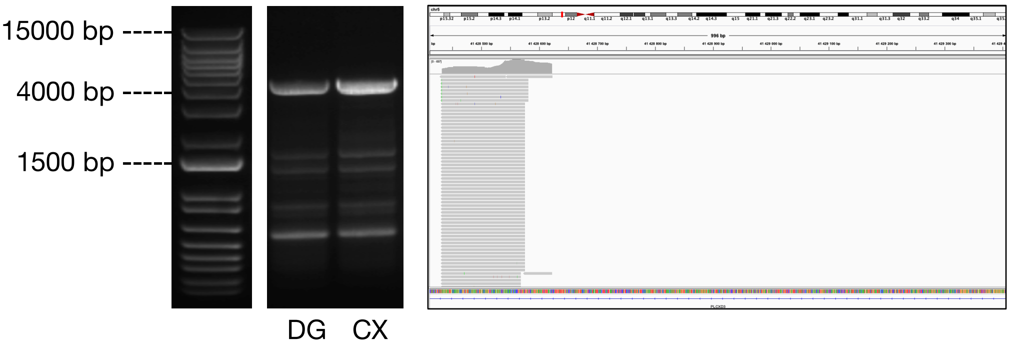


*Lower bound, donor sequenced across multiple experiments.

**L1-276** (Somatic)

| Location | Strand | Number of tagmentation events | Number of single neurons | Number of sample plates |
|---|---|---|---|---|
| chr8:110620382 | + | 11* | 50 | 18 |

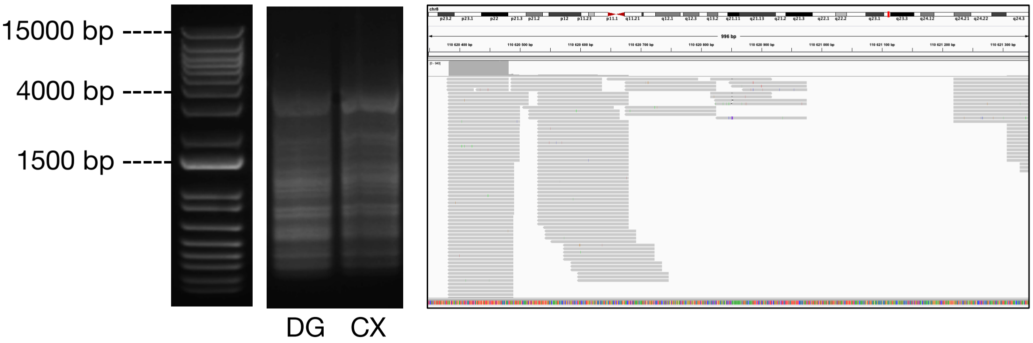


*Lower bound, donor sequenced across multiple experiments.

**L1-30** (Somatic)

| Location | Strand | Number of tagmentation events | Number of single neurons | Number of sample plates |
|---|---|---|---|---|
| chr10:81653425 | - | 3 | 6 | 6 |

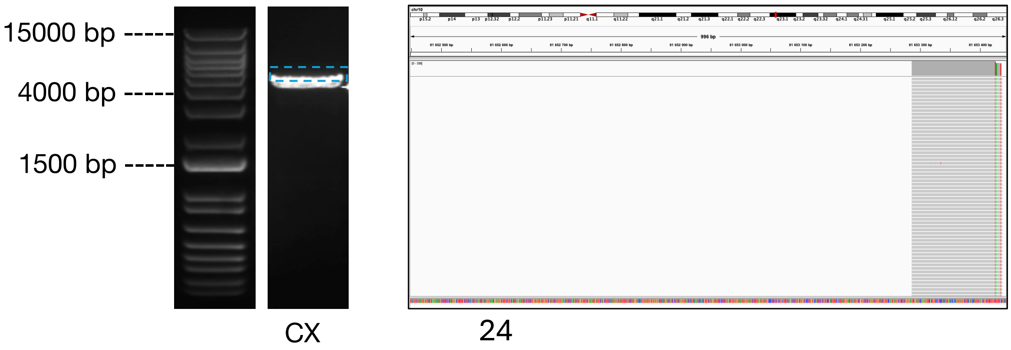

**L1-59** (Somatic)

| Location | Strand | Number of tagmentation events | Number of single neurons | Number of sample plates |
|---|---|---|---|---|
| chr12:86021706 | + | 3 | 8 | 6 |

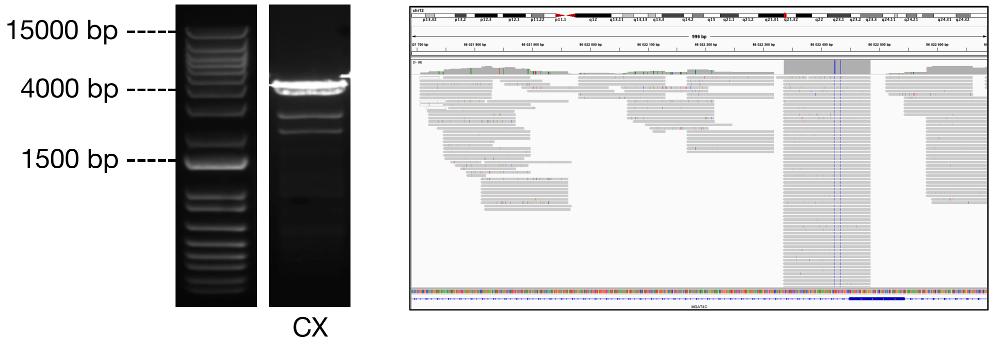


**L1-78** (Somatic)

| Location | Strand | Number of tagmentation events | Number of single neurons | Number of sample plates |
|---|---|---|---|---|
| chr13:112611819 | + | 4 | 12 | 6 |

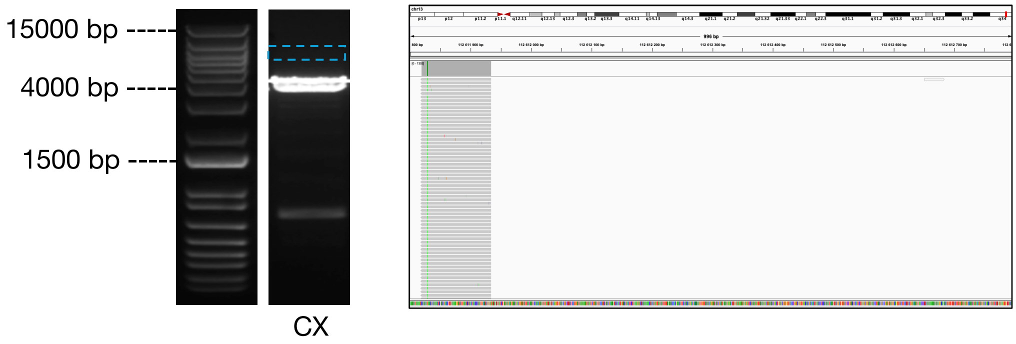


**L1-125** (Somatic)

| Location | Strand | Number of tagmentation events | Number of single neurons | Number of sample plates |
|---|---|---|---|---|
| chr19:14149908 | - | 4 | 12 | 5 |

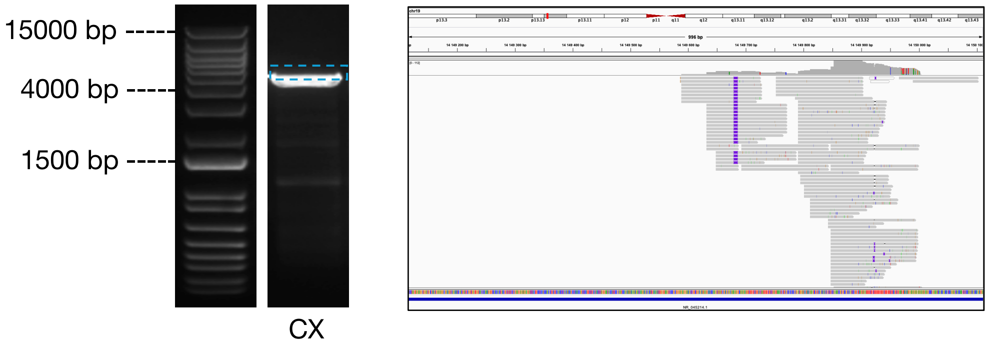

**L1-202** (Somatic)

| Location | Strand | Number of tagmentation events | Number of single neurons | Number of sample plates |
|---|---|---|---|---|
| chr4:11077197 | - | 16 | 22 | 8 |

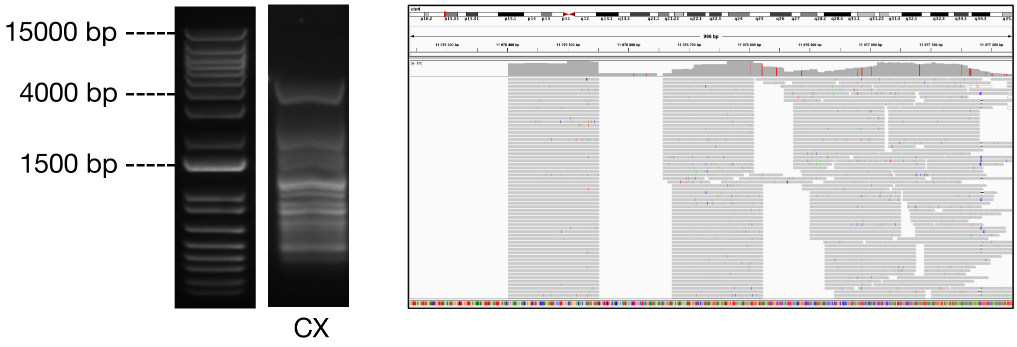


**L1-262** (Somatic)

| Location | Strand | Number of tagmentation events | Number of single neurons | Number of sample plates |
|---|---|---|---|---|
| chr7:154572544 | - | 6 | 12 | 8 |

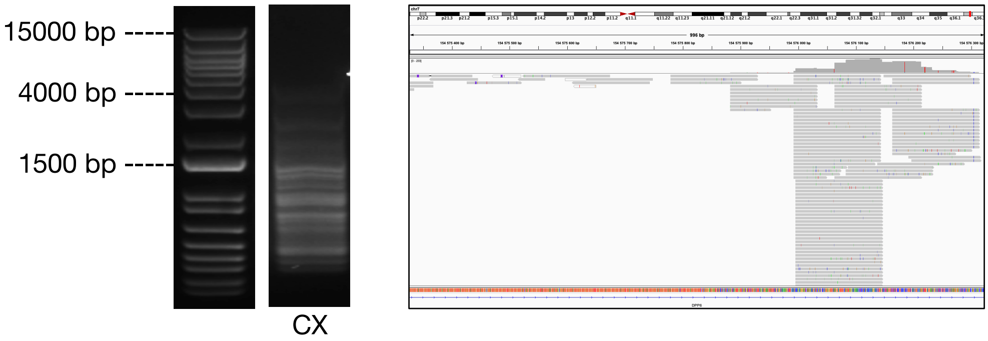


**L1-275** (Somatic)

| Location | Strand | Number of tagmentation events | Number of single neurons | Number of sample plates |
|---|---|---|---|---|
| chr8:108701088 | + | 1 | 1 | 1 |

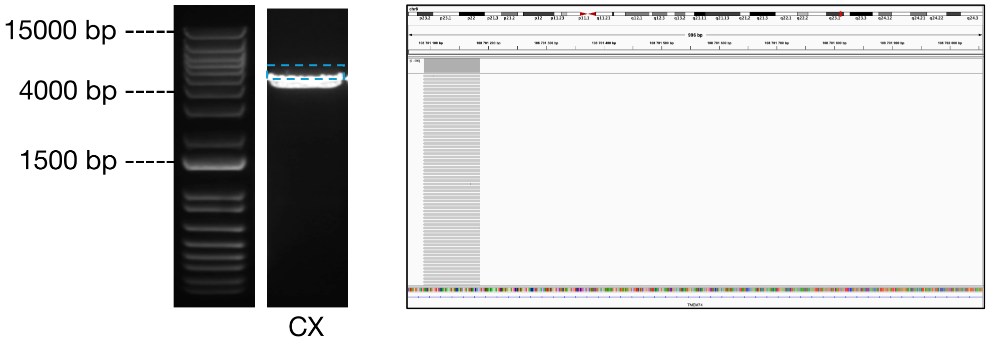

**L1-93** (Somatic)

| Location | Strand | Number of tagmentation events | Number of single neurons | Number of sample plates |
|---|---|---|---|---|
| chr15:24824702 | - | 1 | 1 | 1 |

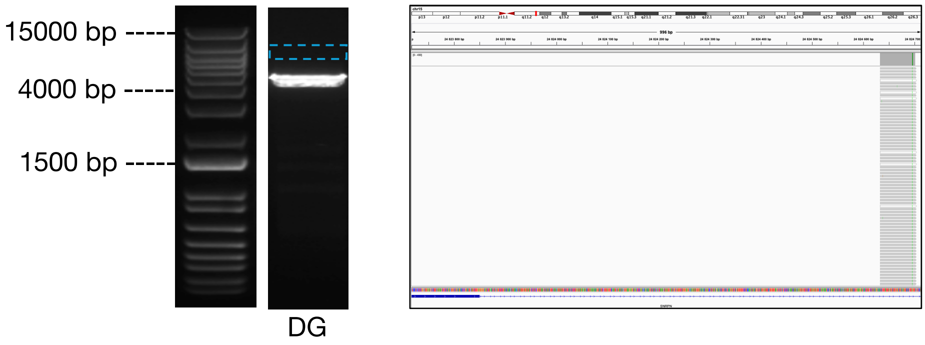


**L1-104** (Somatic)

| Location | Strand | Number of tagmentation events | Number of single neurons | Number of sample plates |
|---|---|---|---|---|
| chr17:2282461 | + | 1 | 1 | 1 |

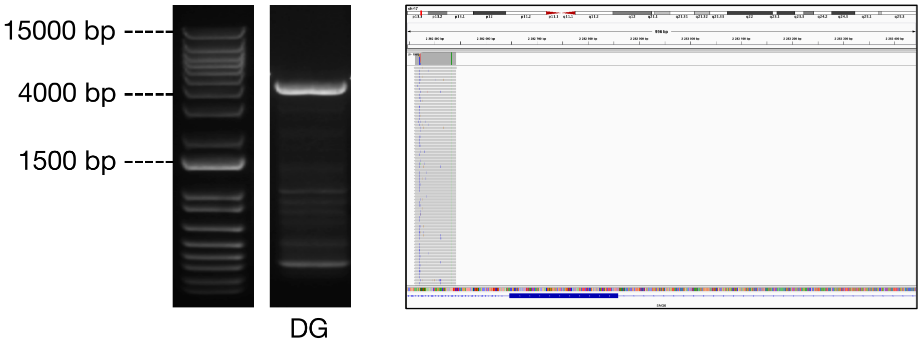


**L1-77** (Somatic)

| Location | Strand | Number of tagmentation events | Number of single neurons | Number of sample plates |
|---|---|---|---|---|
| chr13:107891888 | + | 2 | 3 | 2 |

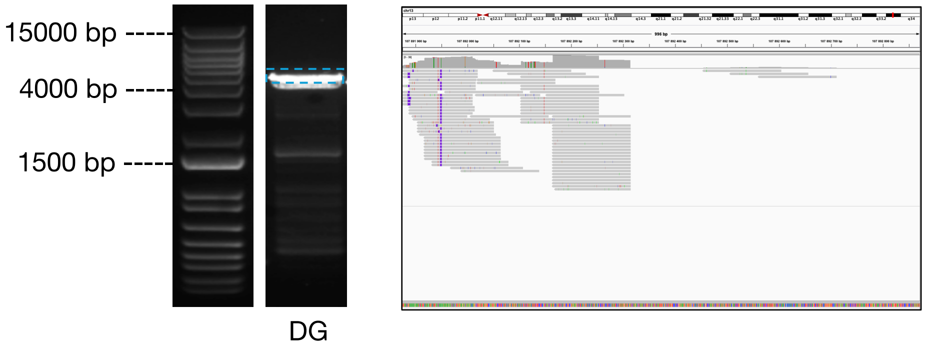

**L1-169** (Somatic)

| Location | Strand | Number of tagmentation events | Number of single neurons | Number of sample plates |
|---|---|---|---|---|
| chr21:15989357 | - | 6 | 9 | 2 |

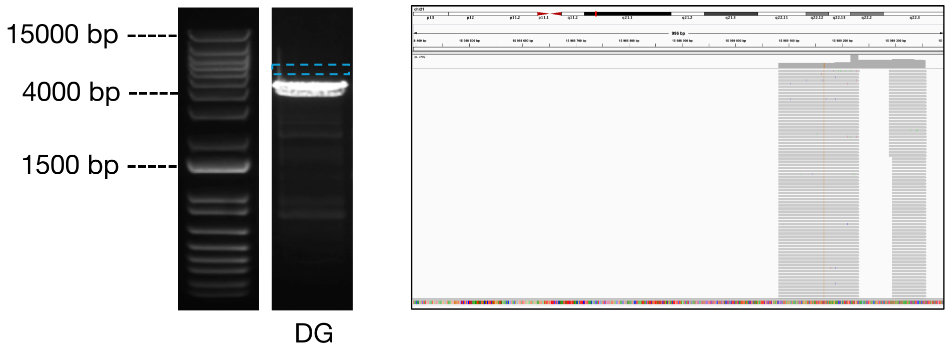


**L1-258** (Somatic)

| Location | Strand | Number of tagmentation events | Number of single neurons | Number of sample plates |
|---|---|---|---|---|
| chr7:124345655 | - | 3 | 5 | 3 |

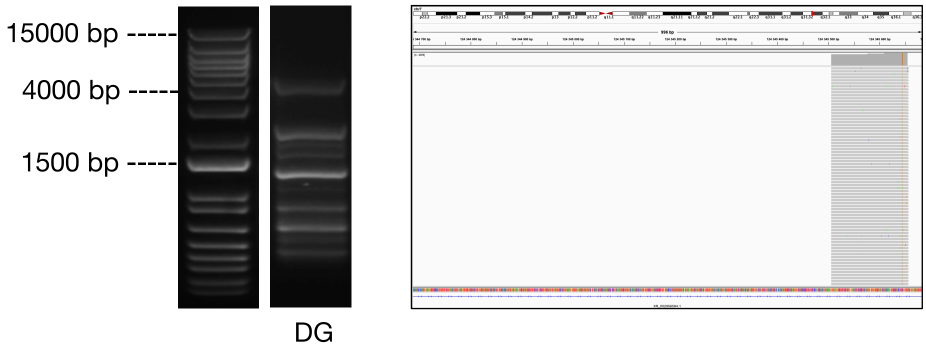


**L1-105** (Somatic)

| Location | Strand | Number of tagmentation events | Number of single neurons | Number of sample plates |
|---|---|---|---|---|
| chr17:5940326 | + | 1 | 9 | 2 |

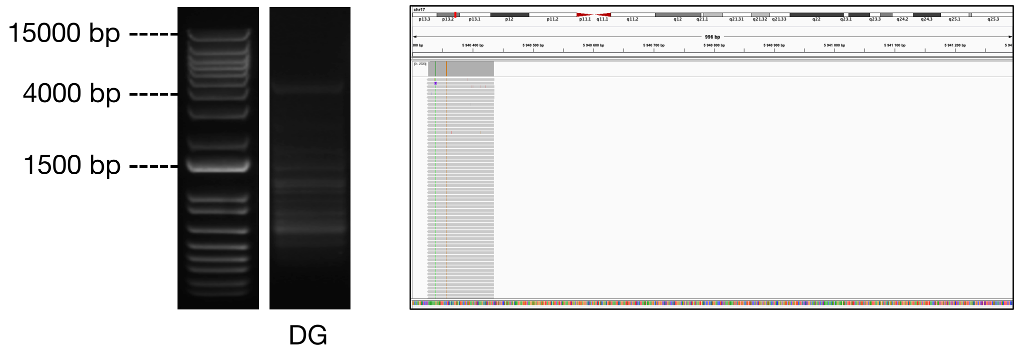

### Categorization of L1s for post-PCR validation rate estimations

We categorized each L1 somatic insertions subjected to PCR validation according to the electrophoresis result and LUSTRE sequencing reads:

PCR validated: PCR supports the existence of a larger DNA fragment than expected. May be separated from the main band to different degree.
PCR inconclusive: No evidence by PCR that a larger DNA fragment was amplified. Band is consistent with the expected size and there is no second larger band observed.
PCR missing: No observable band at all or only clear off-target priming.
IGV: No indication of inconsistencies after manual inspection of read coverage using the Integrated Genome Viewer (IGV). Note that very few insertions fail this criteria.

We assigned each insertion a combination of PCR validated/PCR inconclusive and IGV status. Four different grouping criteria was then used to calculate a validation rate shown in Figure 1:

- PCR validated and IGV
- PCR validated (meaning PCR supports the insertions but IGV has indications of inconsistencies)
- PCR validated/PCR inconclusive and IGV
- All PCR (including PCR missing) and IGV

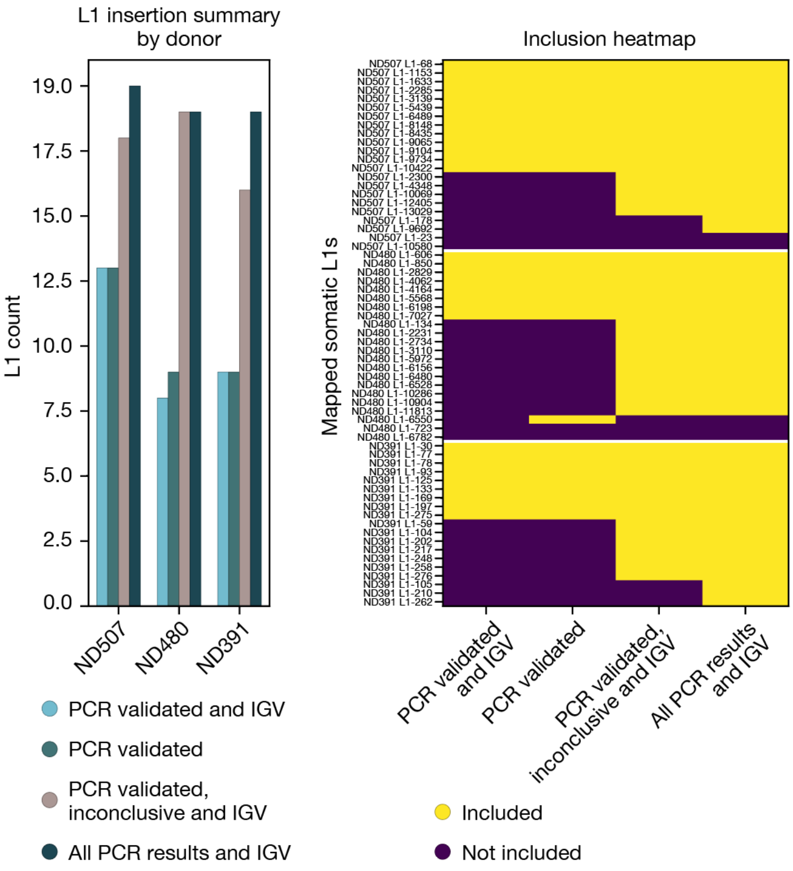


**Figure 1. Summary of categorization for L1 validations.**
Left: Total number of L1 insertions included in group.
Right: L1 insertions included in each group.

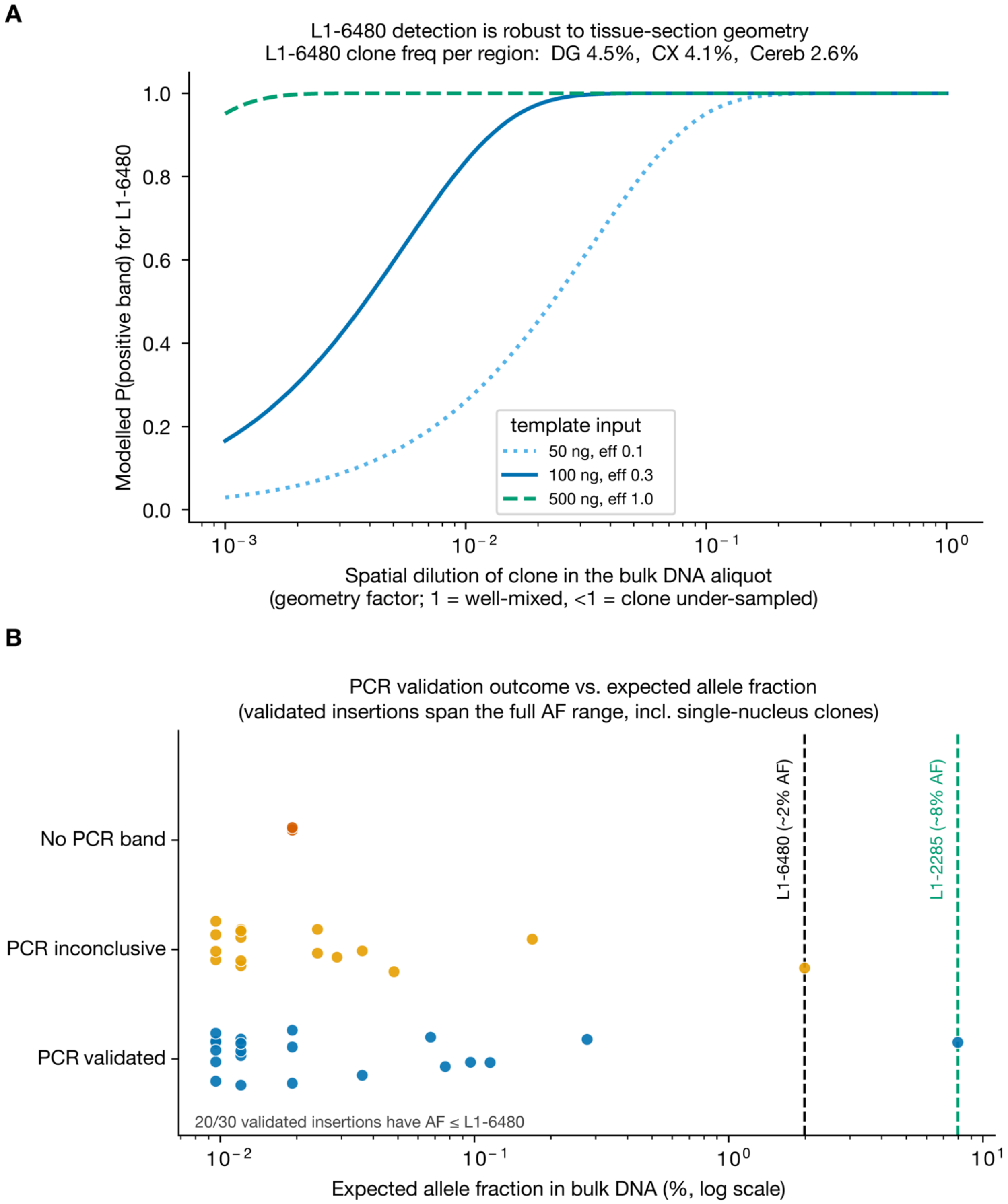
A
L1-6480 detection is robust to tissue-section geometry
L1-6480 clone freq per region: DG 4.5%, CX 4.1%, Cereb 2.6%
Modelled P(positive band) for L1-6480
1.0
0.8
0.6
0.4
0.2
0.0
template input
50 ng, eff 0.1
100 ng, eff 0.3
500 ng, eff 1.0
$10^{-3}$
$10^{-2}$
$10^{-1}$
$10^{0}$
Spatial dilution of clone in the bulk DNA aliquot
(geometry factor; 1 = well-mixed, <1 = clone under-sampled)
B
PCR validation outcome vs. expected allele fraction
(validated insertions span the full AF range, incl. single-nucleus clones)
No PCR band
PCR inconclusive
PCR validated
L1-6480 (~2% AF)
L1-2285 (~8% AF)
20/30 validated insertions have AF ≤ L1-6480
$10^{-2}$
$10^{-1}$
$10^{0}$
$10^{1}$
Expected allele fraction in bulk DNA (%, log scale)

**Figure 2. Modelled PCR sensitivity and summary of PCR validation outcome compared to allele fraction.**

(A) Modelled P(band) for L1-6480 as a function of the spatial dilution of the clone in the bulk DNA aliquot (geometry factor: 1 = well-mixed, <1 = clone under-sampled by a tissue punch), for three representative assay settings (50 ng/ε 0.1; 100 ng/ε 0.3; 500 ng/ε 1.0). Detection power is retained until the clone is diluted more than ~30-fold. L1-6480 is in any case detected in all three regions (dentate gyrus 4.5%, cortex 4.1%, cerebellum 2.6% clone frequency; title), so it is not spatially confined.

(B) Expected allele fraction in bulk DNA for every PCR-tested somatic grouped by outcome: PCR-validated (blue), PCR inconclusive (orange), no band (vermillion). Dashed vertical lines mark L1-6480 (~2% AF) and L1-2285 (~8% AF). Validated insertions extend down to single-nucleus clones (~0.01% AF); 20 of 30 validated insertions have an AF at or below L1-6480's 2%, and L1-6480's AF exceeds two-thirds of validated insertions — so validation success does not track bulk allele fraction. AF: Allele Fraction.

### Validation of L1 insertion sites by bulk whole-genome sequencing

We sequenced 12 WGS libraries from genomic DNA: one per brain region for each of the three donors included in the LUSTRE profiling, and three from an additional fourth donor (NT022) who was not profiled with LUSTRE and included as a negative control. We achieved even coverage across the genome with at least 30X over 90% of the genome and an average above 60X in every library and a median insert size of approximately 300 bp, where mean autosomal depth was 70-115X (Figure 3A-C). GC-dependent coverage varied between libraries (Figure 3D).

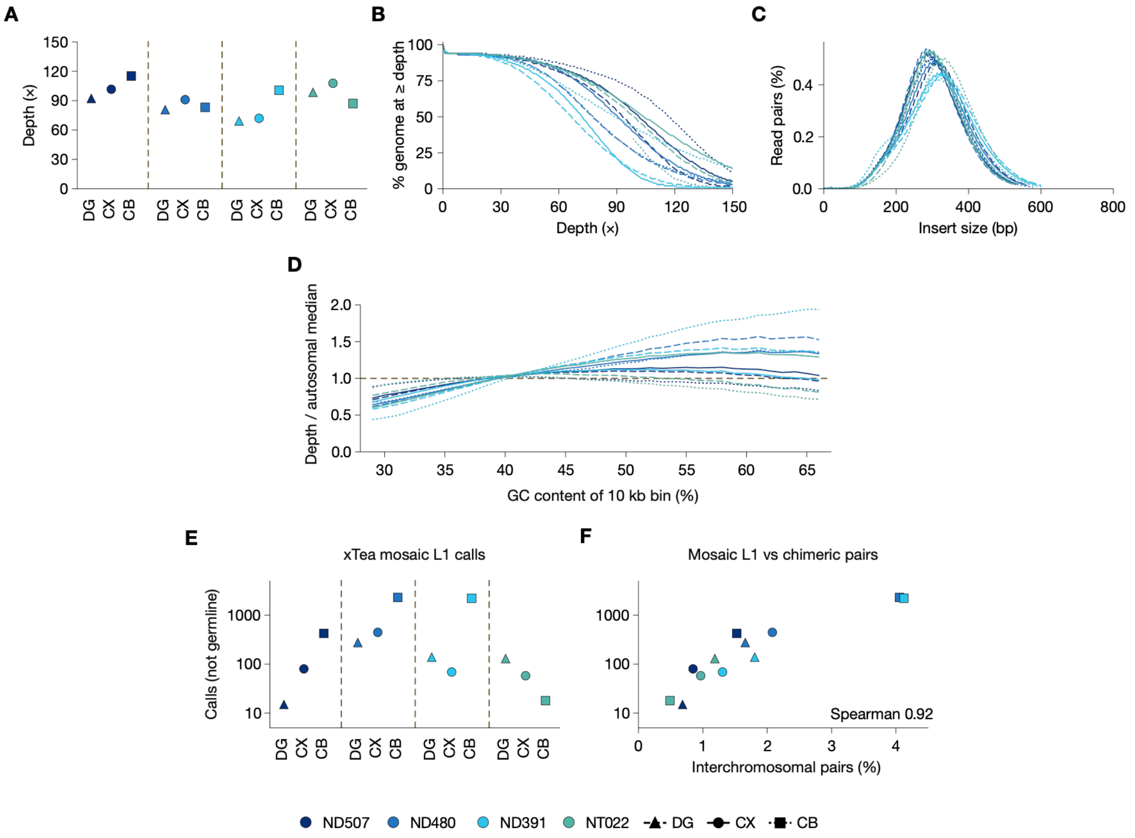


**Figure 3. Quality metrics and mosaic L1 calls of matched whole-genome sequencing data.**
(A) Mean WGS coverage on autosomal chromosomes.
(B) Cumulative coverage across all samples, colored by donor.
(C) Insert sizes across all samples, colored by donor.
(D) GC amplification bias across all samples, colored by donor.
(E) Number of xTea mosaic L1 calls per region and donor.
(F) Correlation between the number of xTea mosaic L1 calls and percentage of interchromosomal ("chimeric") pairs (n = 12 samples).

*De novo* calling of mosaic L1s with xTea returned between 15 and 2,300 calls per library. Importantly, the five most abundant insertions across the three donors were validated using xTea mosaic. The number of calls followed the fraction of interchromosomal read pairs (Spearman's $\rho = 0.92$, $n = 12$; Figure 3E–F), indicating that most calls arose from chimeric library fragments and we therefore did not include them. We instead searched each sample for split reads and discordant read pairs at the intersection of insertion sites identified by LUSTRE (STAR Methods). We then used the local depth and clone size of each L1 to estimate the number of expected supporting reads, and considered insertions with < 3 reads to be below the detection limit of WGS. Of 655 somatic L1s, 11 were above this limit and 8 were validated (L1-12844, L1-202 and L1-34 were not detected; Figure 4A–B). 4 further somatic L1s were validated by WGS although they were initially detected by LUSTRE each in a single nucleus. No insertion had supporting reads in NT022. The read depths of the L1 supporting WGS reads were generally higher compared to the detection by LUSTRE, and in some cases the WGS detected L1s in regions that were not detected by LUSTRE (Figure 4C). Observed counts likely exceed expectation in part because the support metric includes discordant pairs flanking the junction, yielding about 2.8 times the predicted reads even at heterozygous germline insertions (data not shown). LUSTRE may also underestimate clone size where the tissue it sampled differs in clonal composition from the WGS sample. Every validated somatic L1 supported by split reads placed the validated breakpoint within 17 bp of the site predicted by LUSTRE, consistent with the length of a target-site duplication (Figure 5).

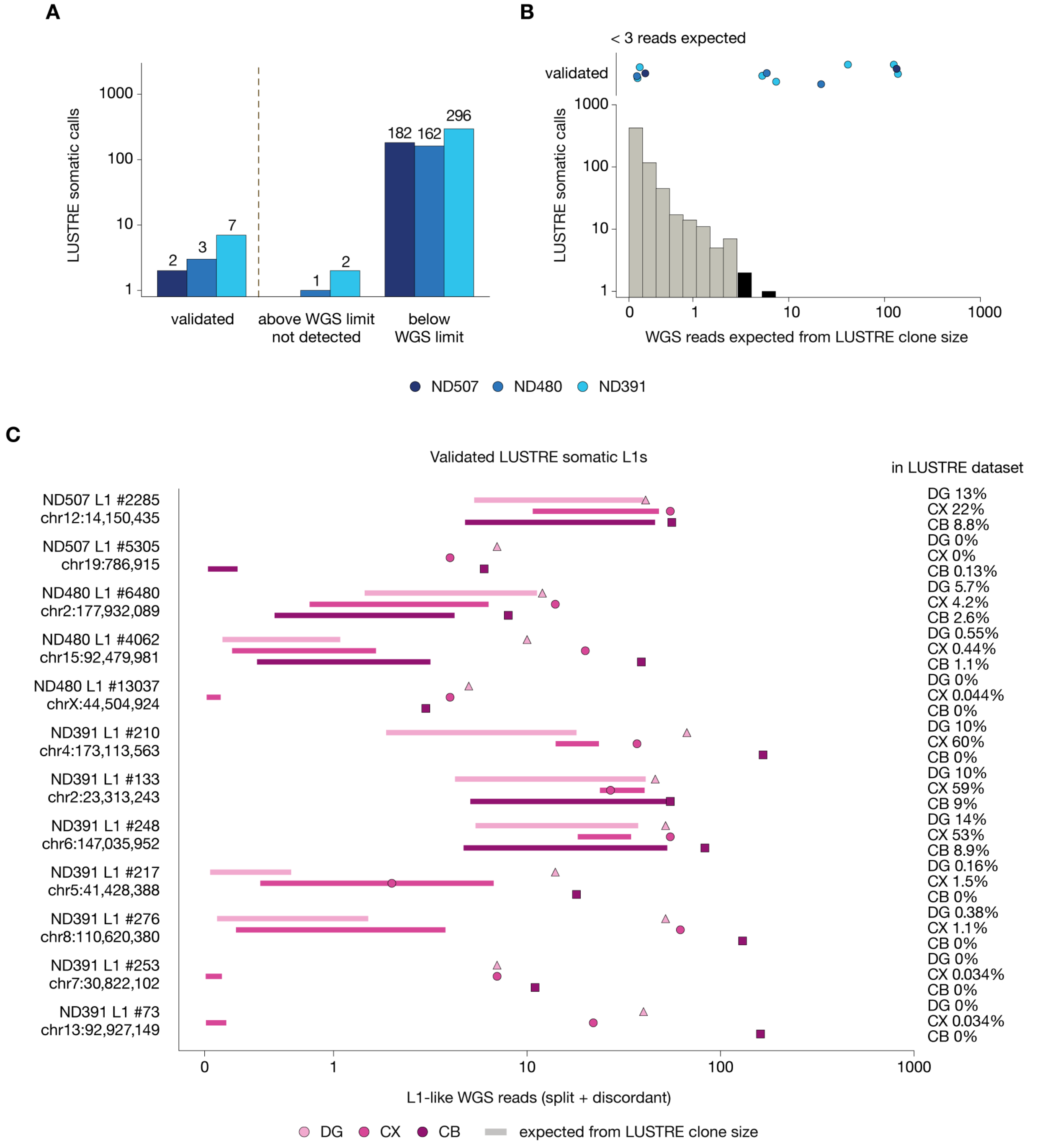

**Figure 4. WGS validation results.**

(A) Bar plot showing the number of somatic L1s validated per donor, the number of L1s above the detection limit and did not validate, and the number of L1s below the detection limit.

(B) Overview of the number of LUSTRE somatic calls, their expected number of WGS reads (histogram) and where the validated somatic L1s were found on this distribution (dot strip, top). Colored by donor.

(C) Overview of each somatic L1 validated by WGS: How many reads were found supporting the L1 event (markers) compared to the expected range from LUSTRE (bars). Missing bars indicate no somatic L1 detected by LUSTRE in that region. Right side shows the percentage of cells the somatic L1 was found in per region.

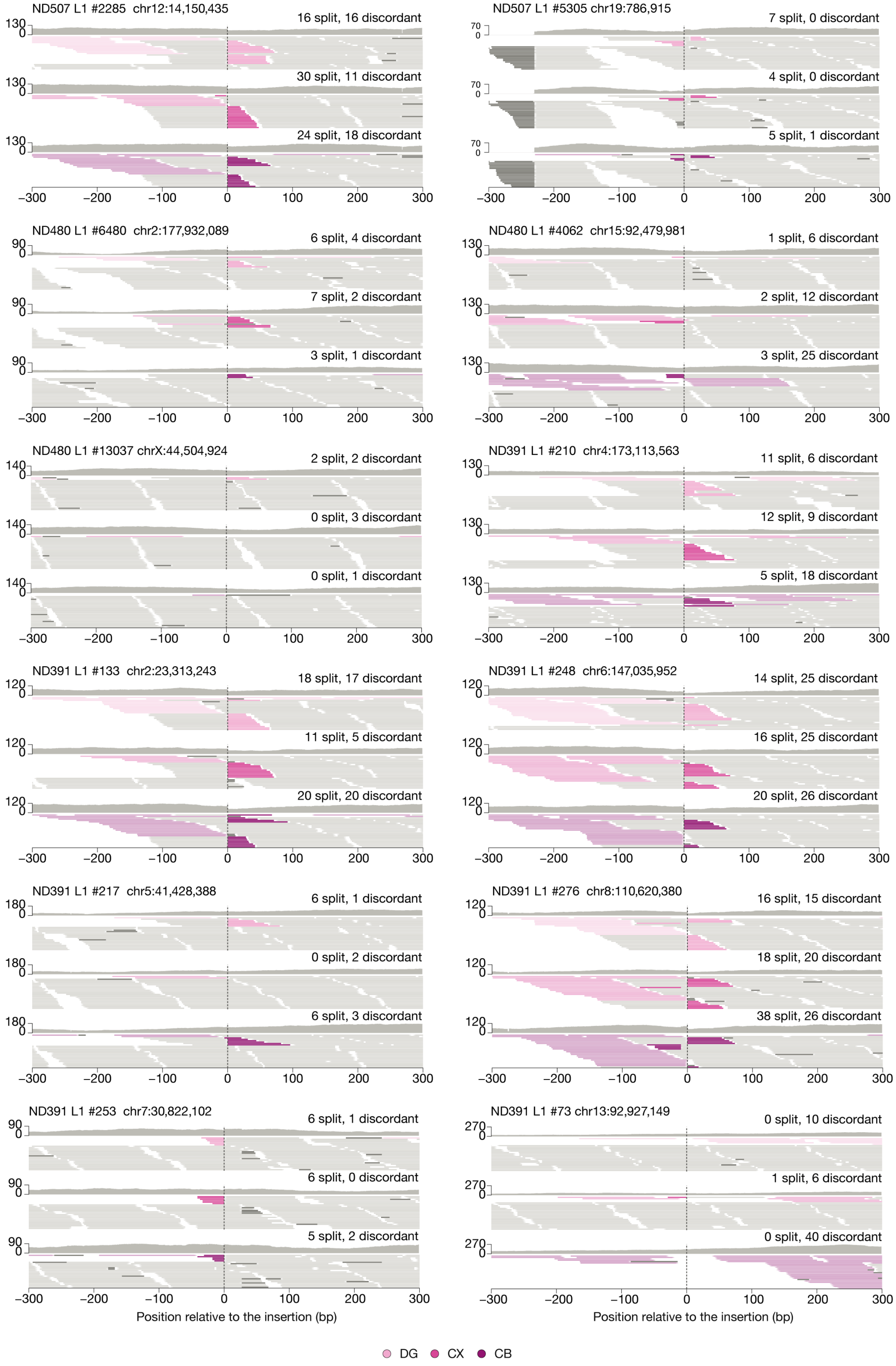
ND507 L1 #2285 chr12:14,150,435
16 split, 16 discordant
30 split, 11 discordant
24 split, 18 discordant
ND507 L1 #5305 chr19:786,915
7 split, 0 discordant
4 split, 0 discordant
5 split, 1 discordant
ND480 L1 #6480 chr2:177,932,089
6 split, 4 discordant
7 split, 2 discordant
3 split, 1 discordant
ND480 L1 #4062 chr15:92,479,981
1 split, 6 discordant
2 split, 12 discordant
3 split, 25 discordant
ND480 L1 #13037 chrX:44,504,924
2 split, 2 discordant
0 split, 3 discordant
0 split, 1 discordant
ND391 L1 #210 chr4:173,113,563
11 split, 6 discordant
12 split, 9 discordant
5 split, 18 discordant
ND391 L1 #133 chr2:23,313,243
18 split, 17 discordant
11 split, 5 discordant
20 split, 20 discordant
ND391 L1 #248 chr6:147,035,952
14 split, 25 discordant
16 split, 25 discordant
20 split, 26 discordant
ND391 L1 #217 chr5:41,428,388
6 split, 1 discordant
0 split, 2 discordant
6 split, 3 discordant
ND391 L1 #276 chr8:110,620,380
16 split, 15 discordant
18 split, 20 discordant
38 split, 26 discordant
ND391 L1 #253 chr7:30,822,102
6 split, 1 discordant
6 split, 0 discordant
5 split, 2 discordant
ND391 L1 #73 chr13:92,927,149
0 split, 10 discordant
1 split, 6 discordant
0 split, 40 discordant
−300 −200 −100 0 100 200 300
Position relative to the insertion (bp)
DG
CX
CB

**Figure 5. Visualization of L1 supporting reads.**
WGS reads supporting the insertion are visualized using the color of the region (order from top to bottom: DG: dentate gyrus, CX: cortex, CB: cerebellum). The darker shade is used to visualize the split reads, while the lighter shade is used to visualize the discordant reads. Read counts only quantify reads found within the [-300,300] bp window.

Among the 63 L1 insertions tested by both methods, WGS validated 5 that long-range PCR had not confirmed (four inconclusive, one without PCR product), including the most widespread somatic L1 in ND480 (L1-6480; Figure 6). Overall, one L1 above the WGS limit was detected by neither method. The remaining 54 were below the WGS limit, where 27 of those were validated by targeted long-range PCR.

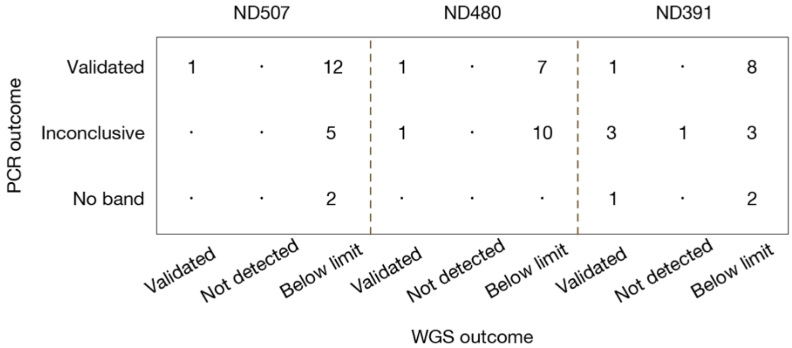


**Figure 6. Comparison between targeted PCR validation and WGS validation.**
This matrix shows the overlap between somatic L1s validated by either PCR, WGS or both. The 5 somatic L1s that failed the IGV check was not included in this comparison, of which none validated by WGS. The matrix therefore sums to 58.
