## Supplementary material for "Dynamics of somatic L1 retrotransposition in the human brain": Note S2

for mutation dating of L1s

Given a random sample of $n$ individuals from a larger population of size $N$, the Kingman's coalescent can be used to estimate coalescent times based on these data[1]. In this model, the genealogical tree coalesces into the most recent common ancestor (MRCA) in a probabilistic manner. Specifically, the waiting time, $W_j$, from when the sample has exactly $j$ ancestors to $j - 1$ ancestors is described by an exponentially distributed random variable with parameter

$$\frac{j(j-1)}{2},\ 2 \le j \ \le n.$$

Interesting quantities associated with a genealogical tree can be computed based on this waiting time, the tree height $T_n$ - the time to the MRCA - and the tree length, $L_n$ - the sum of all branch lengths of the tree. The tree height is the sum of all waiting times,

$$T_n = \sum_{j=2}^{n} W_j$$

and the tree length considers all the ancestors extant at that time,

$$L_n = \sum_{j=2}^{n} j\, W_j$$

We are also interested in estimating the height of specific mutations, to better understand when they occurred during development. This quantity is referred to as $T_m$.

To model the mutations, in our case somatic L1s, the model assumes the mutations occur by a Poisson process with constant rate $\theta/2$. The parameter $\theta$ essentially captures the accumulation of mutations across the entire population. It is further specified by $\theta = 2N\mu$, where $N$ is the population size and $\mu$ is the mutation rate per site per generation. In our case, the mutation rate is not well known. We set it to $10^{-9}$ (order-of-magnitude assumption) and the number of sites to $100$, roughly reflecting the number of active L1. A sensitivity analysis of these variables within this model is presented in Figure S10.

To estimate these parameters, we adapted two algorithms described previously[2]:

**Algorithm to estimate $T_n$, $N$, and $\theta$.**

1. Generate $N \sim Unif(10^2, 10^9)$. Where $Unif$ is the continuous uniform distribution.
2. Calculate $\theta = 2N\mu l$
3. Use the formulas above to obtain $T_n$ and $L_n$.
4. Keep $T_n$ and $N$ if $CDF(k\ |N\mu\, L_n) > 0.05$ and $CDF(k\ |N\mu L_n) < 0.95$. Where k is the number of segregating sites (i.e. insertions). Where $CDF$ is the cumulative distribution function for the Poisson distribution.

**Algorithm to estimate $T_m$.**

1. Set $A_m = m,\ A_n = n,\ T_n = 0,\ L_n = 0$.
2. Generate $W \sim Exp(\frac{A_m(A_m-1)}{2})$. Where $Exp$ is the exponential distribution.
3. Set $T_n = T_n + W$ and $L_n = L_n + A_n W$
4. Set $p = \frac{A_n(A_n - 1)}{A_m(A_m - 1)}$.
5. Set $A_m = A_m - 1$
6. With probability p, set $A_n = A_n - 1$
7. If $A_n > 1$, go to 2.
8. When $A_n = 1$, Accept $A_m$ and $T_n$ with probability $\frac{Pois(k \mid \frac{\theta L_n}{2})}{Pois(k \mid k)}$ or $\frac{\theta L_n}{2} > k$.
9. If rejected, go to 1.
10. If accepted: If $A_m = 1$, set $T_{nm} = 0$. Otherwise generate $T_{nm} = \sum_{j=2}^{A_m} W_j$, where $W_j \sim Exp(\frac{j(j-1)}{2})$. Return $T_m = T_n + T_{nm}$.
